# An epigenome-wide analysis of DNA methylation, racialized and economic inequities, and air pollution

**DOI:** 10.1101/2023.12.07.570610

**Authors:** Sarah Holmes Watkins, Christian Testa, Andrew J. Simpkin, George Davey Smith, Brent Coull, Immaculata De Vivo, Kate Tilling, Pamela D. Waterman, Jarvis T. Chen, Ana V. Diez-Roux, Nancy Krieger, Matthew Suderman, Caroline Relton

**Author notes:** Corresponding author: Sarah Holmes Watkins. address: Oakfield House, Oakfield Grove, Bristol, BS8 2BN, UK.

## Abstract

**Importance:** DNA methylation (DNAm) provides a plausible mechanism by which adverse exposures become embodied and contribute to health inequities, due to its role in genome regulation and responsiveness to social and biophysical exposures tied to societal context. However, scant epigenome-wide association studies (EWAS) have included structural and lifecourse measures of exposure, especially in relation to structural discrimination.

**Objective:** Our study tests the hypothesis that DNAm is a mechanism by which racial discrimination, economic adversity, and air pollution become biologically embodied.

**Design:** A series of cross-sectional EWAS, conducted in My Body My Story (MBMS, biological specimens collected 2008-2010, DNAm assayed in 2021); and the Multi Ethnic Study of Atherosclerosis (MESA; biological specimens collected 2010-2012, DNAm assayed in 2012-2013); using new georeferenced social exposure data for both studies (generated in 2022).

**Setting:** MBMS was recruited from four community health centers in Boston; MESA was recruited from four field sites in: Baltimore, MD; Forsyth County, NC; New York City, NY; and St. Paul, MN.

**Participants:** Two population-based samples of US-born Black non-Hispanic (Black NH), white non-Hispanic (white NH), and Hispanic individuals (MBMS; n=224 Black NH and 69 white NH) and (MESA; n=229 Black NH, n=555 white NH and n=191 Hispanic).

**Exposures:** Eight social exposures encompassing racial discrimination, economic adversity, and air pollution.

**Main **outcome**:** Genome-wide changes in DNAm, as measured using the Illumina EPIC BeadChip (MBMS; using frozen blood spots) and Illumina 450k BeadChip (MESA; using purified monocytes). Our hypothesis was formulated after data collection.

**Results:** We observed the strongest associations with traffic-related air pollution (measured via black carbon and nitrogen oxides exposure), with evidence from both studies suggesting that air pollution exposure may induce epigenetic changes related to inflammatory processes. We also found suggestive associations of DNAm variation with measures of structural racial discrimination (e.g., for Black NH participants, born in a Jim Crow state; adult exposure to racialized economic residential segregation) situated in genes with plausible links to effects on health.

**Conclusions and Relevance:** Overall, this work suggests that DNAm is a biological mechanism through which structural racism and air pollution become embodied and may lead to health inequities.

**Key points:** *Question:* Could DNAm be a mechanism by which adversity becomes embodied?

*Findings:* Traffic-related air pollution exposure may induce epigenetic changes related to inflammatory processes and there are suggestive associations with measures of structural racism

*Meaning:* DNAm may be a biological mechanism through which structural racism and air pollution become biologically embodied

## Introduction

Recent advances enabling large population-based epigenetic studies are permitting researchers to test hypotheses linking socially-patterned exposures, gene regulation, and health inequities ^1-3^. DNA methylation (DNAm) is a plausible biological mechanism by which adverse social exposures may become embodied ^4,5^, because 1) it plays an active role in genome regulation ^6-8, 2^) it changes in response to environmental exposures ^1,9^ and internal human physiology like ageing ^2^ and inflammation ^10^, and 3) induced changes can be long-lasting ^11-14^. There is a growing literature reporting associations between DNAm and environmental factors to which social groups are unequally exposed; a recent review ^15^ found associations between DNAm and measures of socio-economic position (SEP), including income, education, occupation, and neighbourhood measures; and illustrated timing and duration of exposure is important. Exposure to toxins, including air pollution, is often inequitable between social groups ^16,17^. A number of EWAS have identified associations with particulate matter ^18-21^ and oxides of nitrogen (NOx) ^21,22^; although there is little replication between studies, and some studies have failed to find effects of particulate matter ^22,23^, NOx ^23^, and residential proximity to roadways ^24^. Two EWAS have each found two (non-overlapping) DNAm sites associated with experience of racial discrimination, one in first generation Ghanaian migrants living in Europe ^3^, and one in African American women ^25^; given population and migration differences the lack of replication is perhaps not surprising.

However, no EWAS has yet examined associations between DNAm and exposure to racial discrimination and economic adversity, both at individual and structural levels, and measured at different points in the lifecourse, in the same group of people. This is important because it is not clear if the different timing, duration, and levels of these adverse exposures are embodied in different ways involving differing biological pathways. Supporting attention to these issues is a growing body of research documenting how exposure to health-affecting factors such as toxins, quality healthcare, education, fresh food, and green spaces are determined by the way dominant social groups have structured society, which in turn results in health inequities between dominant social groups and groups they have minoritized ^4,26^. Structural racism (the totality of ways in which society discriminates against racialized groups ^27^), for example, results in people of colour often disproportionately bearing the burden of adverse exposures and economic hardship ^4,28^, thus driving racialized health inequities ^29^. Associations between structural racism and cardiovascular health have been shown for discriminatory housing policies and continuing neighbourhood racial segregation ^30,31^; with the historical legacy of slavery ^32^; and with state-level institutional domains ^33^. Associations have also been shown for diabetes outcomes in the US ^34^ and globally ^35^.

Guided by the ecosocial theory of disease distribution ^4,5^, we tested the hypothesis that DNAm is a biological mechanism by which embodiment of structural racial discrimination, economic hardship, and air pollution may occur. We tested our study hypothesis using data from US-born participants in two US population based studies with similar exposure data: our primary study, the My Body My Story study (MBMS), and the Multi-Ethnic Study of Atherosclerosis study (MESA), which we use for evidence triangulation ^36^ due to differences between the two study populations.

## Methods

### Participants

This study utilises biological specimens obtained in 2010-2012 from MBMS and MESA, two US population-based studies that contain similar data on the study exposures. In 2021-2022, the study team newly conducted epigenetic assays for MBMS and added new georeferenced social exposure data. Full study descriptions are in the supplementary materials. Our analyses comprised 293 participants (224 Black and 69 white) from MBMS; and 975 participants from MESA who were US- born (229 Black, 555 white NH, additionally including 191 Hispanic).

### Social exposures

We tested the relationship between DNAm and eight variables relating to exposure to racial discrimination (both structural and self-reported), economic hardship, and air pollution; these are described in detail in Supplementary Table 2.

### DNA methylation

For detailed description of DNA extraction and DNAm data generation, please see the Supplementary materials. Briefly, for MBMS DNA was extracted from frozen blood spots in 2021, and data were generated using the Illumina Infinium MethylationEPIC Beadchip. For MESA, DNA was extracted from purified monocytes in 2012-2013 and data were generated using the Illumina Infinium HumanMethylation450 BeadChip. We used DNAm beta values for both studies, which measure DNAm on a scale of 0 (0% methylation) to 1 (100% methylation).

### Participant stratification

EWAS were stratified by self-reported membership of racialized groups, for two reasons. Firstly, for most of our exposures, different constructs are represented between the racialized groups; for example, being born in a Jim Crow state means something very different for individuals who identify as Black versus white. Secondly, stratification prevents potential confounding by racialized group due to exposure and a degree of genetic differences between groups. Racialized groups are social constructs that are changeable and dependent on local context ^37^; they are important to our research question because group membership is pertinent to the experience of social inequities perpetuated by structural racial discrimination.

### EWAS

All EWAS were conducted using linear regression models implemented using the R package meffil ^38^. Many exposures had low levels of missing data, complete case numbers for each EWAS can be found in Table 3. EWAS details can be found in the Supplementary materials; briefly, we adjusted for age, reported gender (MBMS)/sex (MESA), smoking status, blood cell count proportions, and batch effects.

### Sensitivity analysis

In MESA we conducted a sensitivity analysis to test whether our results were influenced by population stratification; details are in the Supplementary materials. Additional sensitivity analysis restricted the MESA analysis to participants recruited from the Baltimore and New York sites, because these cities bear the greatest similarity to the Boston area in terms of geographical location, city environment, and social histories.

### Meta-analysis

We meta-analysed associations with air pollution within MBMS and within MESA because air pollution is the only exposure we tested that we would hypothesize to have the same meaning, and therefore biological effect, for all individuals. We used METAL ^39^ to meta-analyse effect sizes and standard errors of the EWAS summary statistics of each racialized group, for black carbon/LAC and NOx.

### Functional relevance of sites passing the genome-wide threshold

For DNAm sites associated with an exposure, we used the UCSC genome browser to identify genomic regions. For sites within known genes, we used GeneCards (https://www.genecards.org/) and literature searches to identify putative gene functions. We used the EWAS Catalog to determine if associations between DNAm sites and other traits had been reported in previous studies. Where multiple DNAm sites were associated with an exposure we performed gene set enrichment analysis using the missMethyl R package ^40^.

**Figure 1:**
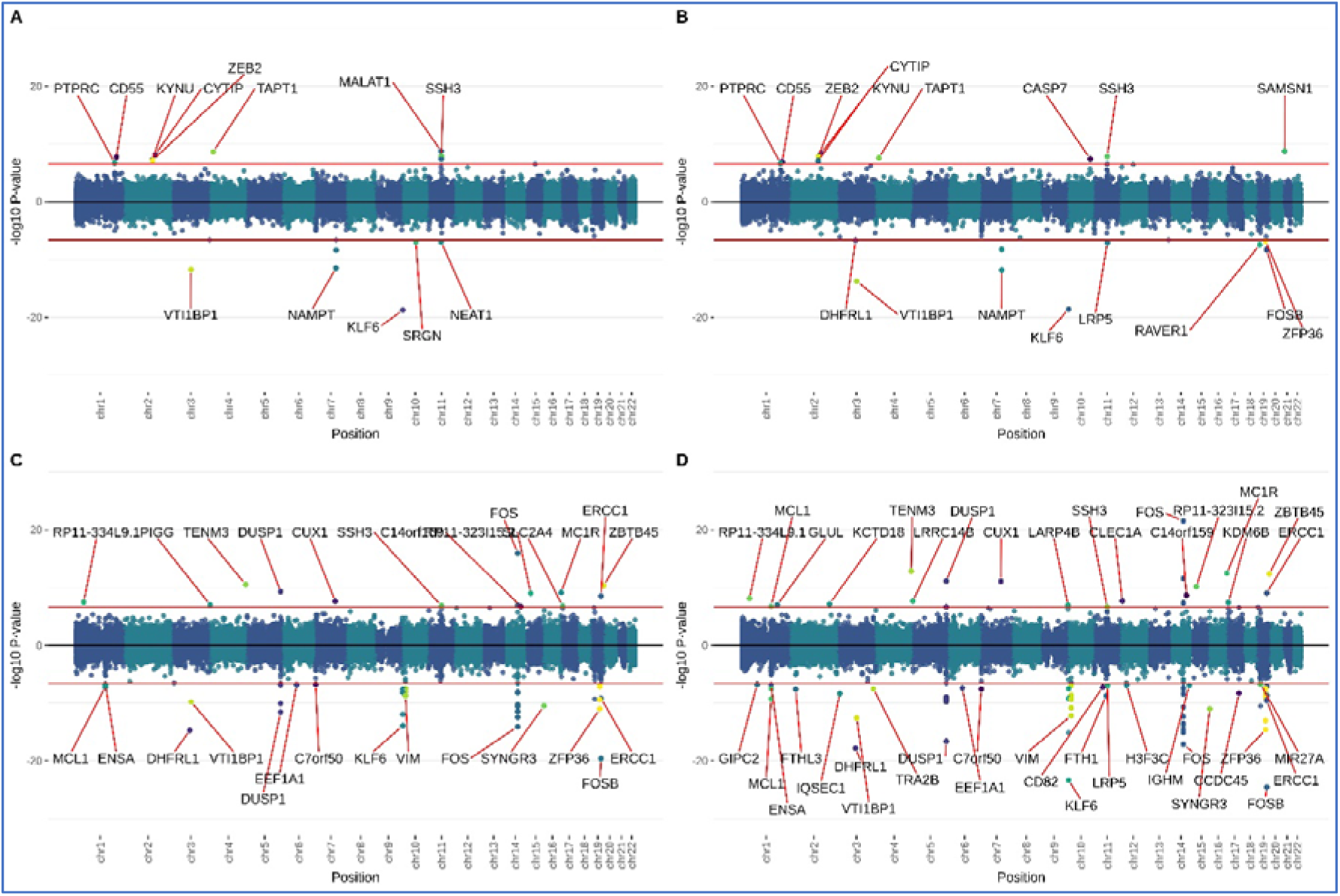
MESA air pollution meta-analysis miami plots. **A**: MESA full cohort LAC meta-analysis. **B:** MESA full cohort NOx meta-analysis. **C:** MESA subgroup LAC meta-analysis. **D:** MESA subgroup NOx meta-analysis.

### Biological enrichments of top sites

Following each EWAS we performed analyses to ascertain whether DNAm sites associated with our exposures indicate effects on particular biological pathways, processes or functions. Details are in the Supplementary materials; briefly, we conducted gene set enrichment analyses, and for enrichments of tissue-specific chromatin states, genomic regions and transcription factor binding sites (TFBS).

### Lookup of associations in *a priori* specified genomic locations

We hypothesised a priori that our EWAS would detect DNAm sites that have been robustly associated with our study exposures, or factors that might relate to our exposures, in previous studies. See Supplementary materials for details.

## Results

### Participant characteristics

Both cohorts include racialized groups that are underrepresented in epigenetic studies. Beyond this, substantial differences existed between the racialized groups within and across MBMS and MESA. Overall, MBMS participants were on average 21 years younger than MESA participants, had less variability in exposure to air pollution, and far more were current smokers. In both studies, Black NH compared to white NH participants had higher BMI, rates of smoking, impoverishment, lower education, rates of self-reported exposure to racial discrimination, and were more likely to be born in a Jim Crow state and live in a neighbourhood with extreme concentrations of low-income persons of colour. In MESA, Hispanic participants reported the lowest levels of personal and parental education.

**Table 1:** Characteristics of MBMS and MESA participants. ^1^Jim Crow states are the 21 US states (plus the District of Columbia) which permitted legal racial discrimination prior to the 1964 US Civil Rights Act. ^2^Participants’ ratio of household income in 2010 dollars to the US 2010 poverty line given household composition. ^3^Census tract measure of economic and racialized segregation, scored from -1 to 1 ^4^NOx measurements were used to construct a weighted score of roadway pollution ^5^Validated self-report questionnaire measuring the number of domains of exposure to racial discrimination. Score range 0- 9, categorised into 0, 1-2, 3 ^6^Validated self-report questionnaire measuring the number of domains of exposure to racial discrimination; combined with the attribution aspect from EDS (everyday discrimination scale) to enable comparability between EOD and MDS. Score range 0-5, categorised into 0, 1-2, 3+

| Variable |  | MBMS:<br>Black NH | MBMS:<br>white NH | MESA:<br>Black NH | MESA:<br>white NH | MESA:<br>Hispanic |
| --- | --- | --- | --- | --- | --- | --- |
| Total N |  | 224 | 69 | 229 | 555 | 191 |
| **Sociodemographic characteristics** |  |  |  |  |  |  |
| Age: mean (SD) |  | 49.02 (7.8) | 48.7 (8.3) | 71 (8.9) | 70.1 (9.5) | 68.5 (8.9) |
| Gender: N (%) women |  | 135<br>(60.3%) | 49 (71%) | 133<br>(58.1%) | 264<br>(47.6%) | 86 (45%) |
| BMI: mean (SD) |  | 32.1 (7.7) | 29.7 (7.2) | 30.6 (5.7) | 28.7 (5.3) | 30.8 (5.5) |
| Smoking: N (%) | Current | 115<br>(51.3%) | 24 (34.8%) | 31 (13.7%) | 44 (8%) | 16 (8.6%) |
|  | Former | 31 (13.8%) | 23 (33.3%) | 101<br>(44.9%) | 262<br>(47.7%) | 80 (43%) |
|  | Never | 78 (34.8%) | 22 (31.9%) | 93 (41.5%) | 243<br>(44.3%) | 90 (48.4%) |
|  | Missing | 0 | 0 | 4 (1.7%) | 6 (1.1%) | 5 (2.6%) |
| **Childhood exposure to racialized and economic adversity:** |  |  |  |  |  |  |
| Born in a Jim Crow state<sup>1</sup>: N (%) yes |  | 71 (31.7%) | 2 (3%) | 165<br>(72.1%) | 166<br>(29.9%) | 19 (9.9%) |
| Parent's highest education: N (%) | <High school | 29 (18.4%) | 8 (14%) | 95 (42.2%) | 161<br>(29.3%) | 129 (69.7%) |
|  | >= High school and <4yr college | 94 (59.5%) | 24 (42.1%) | 106<br>(47.3%) | 258 (47%) | 51 (27.6%) |
|  | 4+ years college | 35 (22.2%) | 25 (43.9%) | 24 (10.7%) | 130<br>(23.7%) | 5 (2.7%) |
|  | Missing | 66 (29.5%) | 12 (17.4%) | 4 (1.7%) | 6 (1.1%) | 6 (3.1%) |
| Participant's education: N (%) | <High school | 34 (15.2%) | 8 (11.6%) | 23 (10%) | 21 (3.8%) | 35 (18.3%) |
|  | >= High school and <4yr college | 161<br>(71.9%) | 33 (47.8%) | 175<br>(76.4%) | 413<br>(74.4%) | 140 (73.3%) |
|  | 4+ years college | 29 (12.9%) | 28 (40.6%) | 31 (13.5%) | 121<br>(21.8%) | 16 (8.4%) |
|  | Missing: | 0 | 0 | 0 | 0 | 0 |
| **Adult exposure to racialized and economic adversity:** |  |  |  |  |  |  |
| Household income to poverty ratio<sup>2</sup>: mean (SD) |  | 2.2 (2.2) | 2.9 (2.3) | 3.9 (2.3) | 4.8 (2.9) | 3.3 (2.1) |
| Missing |  | 34 (15.2%) | 3 (4.3%) | 9 (3.9%) | 24 (4.3%) | 7 (3.7%) |
| Index of Concentration at the Extremes for racialized economic segregation<sup>3</sup>: mean(SD) |  | -0.07 (0.2) | 0.19 (0.2) | -0.11 (0.2) | 0.16 (0.2) | 0.09 (0.2) |
| Missing |  | 0 | 0 | 2 (0.9%) | 4 (0.7%) | 12 (6.3%) |
| Black carbon (µg/m3): mean (SD) |  | 0.64 (0.1) | 0.63 (0.17) |  |  |  |
| Missing |  | 0 | 0 |  |  |  |
| Light absorption coefficient (10<sup>-5</sup>/m): mean (SD) |  |  |  | 0.89 (0.35) | 0.6 (0.3) | 0.7 (0.4) |
| Missing |  |  |  | 14 (6.1%) | 23 (4.1%) | 11 (5.6%) |
| Pollution Proximity Index<sup>4</sup> (scale of 0-5): mean (SD) |  | 4.3 (1.1) | 3.9 (1.4) |  |  |  |
| *Missing* |  | 5 (2.2%) | 0 |  |  |  |
| Oxides of nitrogen (NOx, parts per billion): mean (SD) |  |  |  | 31.9 (16.2) | 21.55 (12.2) | 27 (16.4) |
| *Missing* |  |  |  | 14 (6.1%) | 23 (4.1%) | 11 (5.6%) |
| Experiences of Discrimination (EOD, N of domains)<sup>5</sup>: N (%) | 0 | 30 (13.4%) | 35 (50.7%) |  |  |  |
|  | 1-2 | 52 (23.2%) | 24 (34.8%) |  |  |  |
|  | 3+ | 140 (62.5%) | 10 (14.5%) |  |  |  |
|  | *Missing* | 2 (0.9%) | 0 |  |  |  |
| Major Discrimination Scale (MDS, N of domains)<sup>6</sup>: N (%) | 0 |  |  | 129 (56.6%) | 534 (96.4%) | 131 (68.6%) |
|  | 1-2 |  |  | 79 (34.6%) | 20 (3.6%) | 53 (27.7%) |
|  | 3+ |  |  | 20 (8.8%) | 0 | 7 (3.7%) |
|  | *Missing* |  |  | 1 (0.4%) | 1 (0.2%) | 0 |
| **Predicted cell count proportions** |  |  |  |  |  |  |
| B Cell |  | 0.08 (0.02) | 0.06 (0.01) | 0.04 (0.03) | 0.03 (0.02) | 0.03 (0.02) |
| CD4+T cells |  | 0.17 (0.05) | 0.15 (0.04) | 0.04 (0.02) | 0.03 (0.03) | 0.03 (0.01) |
| CD8+T cells |  | 0.005 (0.02) | 0.002 (0.01) | 0.002 (0.006) | 0.0007 (0.003) | 0.0006 (0.003) |
| Monocytes |  | 0.124 (0.02) | 0.116 (0.02) | 0.9 (0.05) | 0.91 (0.04) | 0.92 (0.04) |
| Neutrophils |  | 0.55 (0.1) | 0.62 (0.08) | 0.0003 (0.002) | 0.0008 (0.005) | 0.0009 (0.007) |
| Natural Killer |  | 0.1 (0.04) | 0.09 (0.04) | 0.015 (0.01) | 0.012 (0.01) | 0.01 (0.01) |
| Eosinophils |  | 0.007 (0.02) | 0.003 (0.009) | 0.01 (0.01) | 0.01 (0.01) | 0.01 (0.01) |

### EWAS results and biological interpretation

In MBMS, among the Black NH participants one DNAm site, in ZNF286B, was associated with being born in a Jim Crow state. Another DNAm site, PLXND1, was associated with participants having less than high school education. Among white NH participants, no associations passed the genome-wide threshold. See Table 2 for details of gene functions; Table 3 for numbers of associated EWAS sites; and Supplementary figures 1 and 2 for Miami plots. In MESA, two DNAm sites were associated with racialized economic segregation – one in Black NH participants (in FUT6) and one in white NH participants (a CpG previously associated with BMI); and in Black NH participants one DNAm site (in PDE4D) was associated with an MDS score of 0. The majority of associations in MESA were related to air pollution exposure – among Black NH participants, 12 sites with LAC and 22 sites with NOx. Notably, many of these sites are clustered in genes with putative roles in immune responses and are known to interact with one another, including KLF6, MIR23A, FOS, FOSB, ZFP36 and DUSP1. Among the MESA white NH participants, four DNAm sites were associated with both LAC and NOx, and an additional 3 uniquely associated with LAC. Associations of 53 DNAm sites with birth in a Jim Crow state were the result of confounding by air pollution (see Supplementary Materials). Among Hispanic participants, one site was associated with LAC (NPNT) and one with NOx (ADPRHL1). See Table 3 for numbers of associations for all EWAS performed and Supplementary Figures 3-5 for corresponding Miami plots.

**Table 2:**
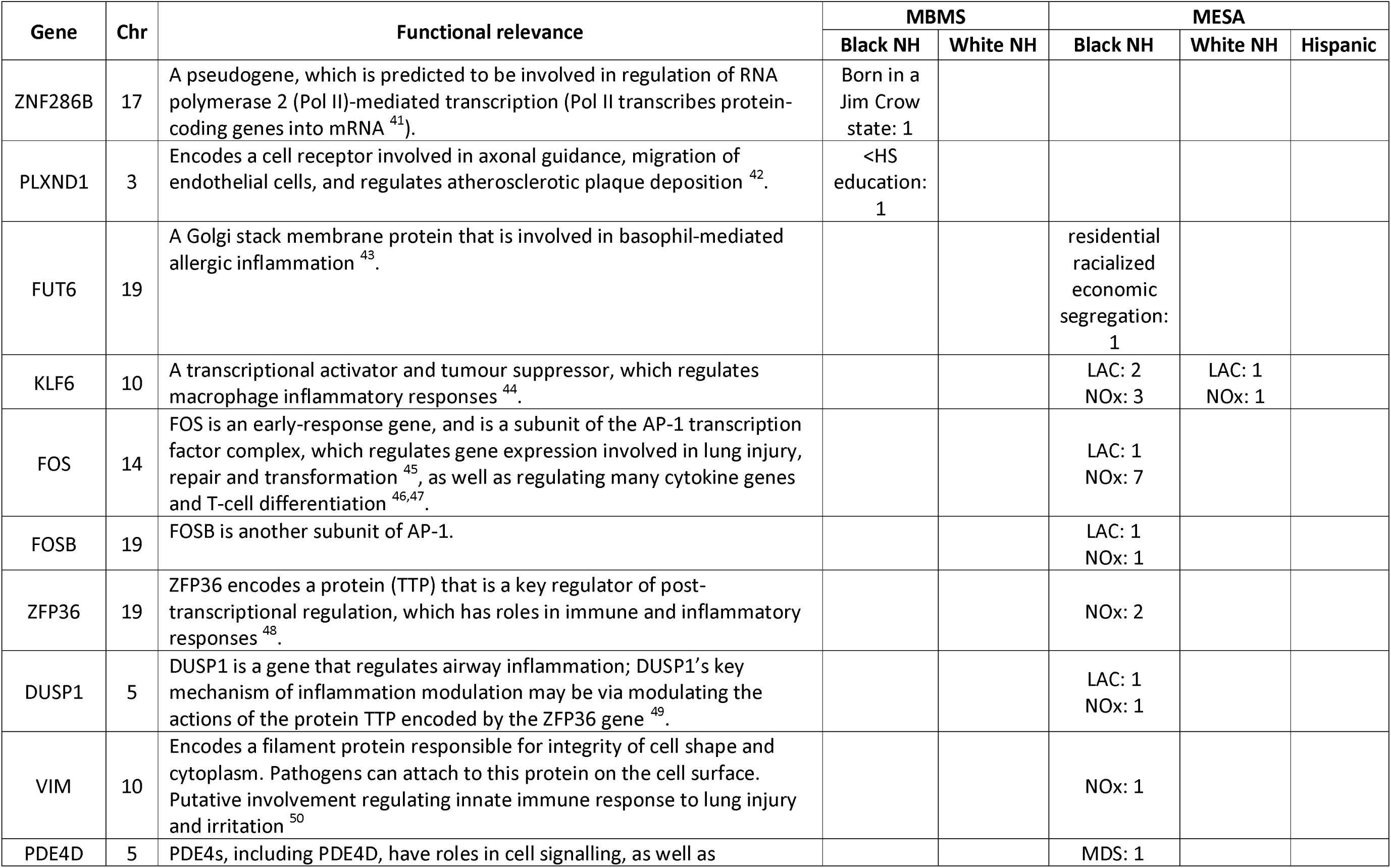
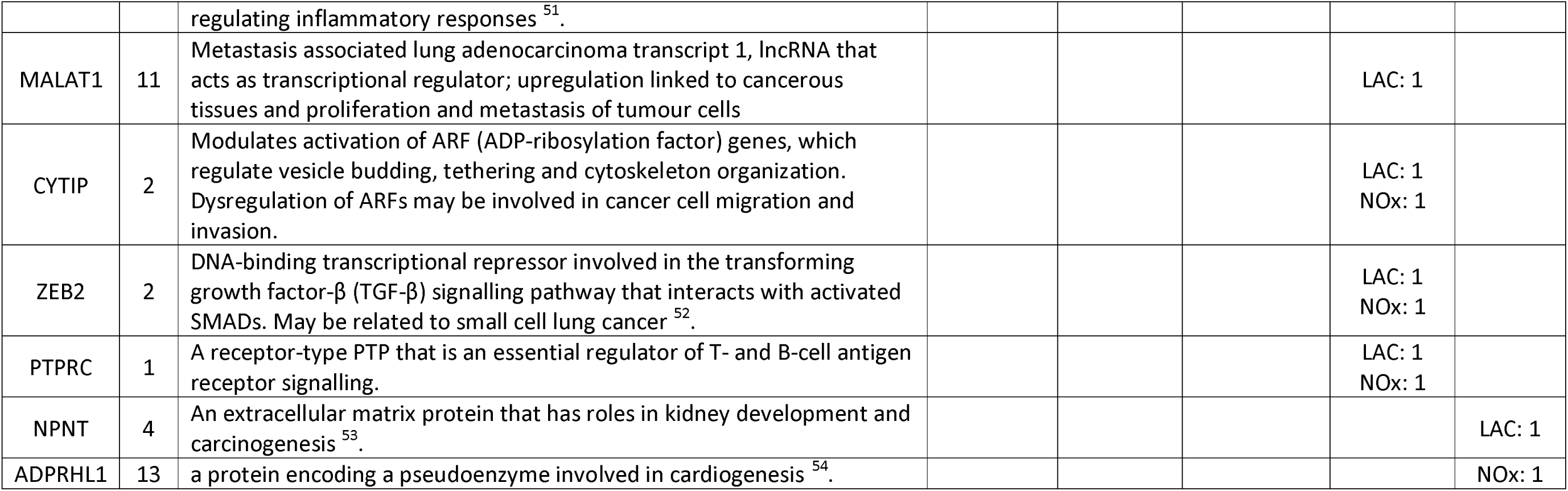
Putative functions of genes in which the top exposure-associated DNAm sites sit; with details of how many sites within that gene were identified, and in which main analysis EWAS y were identified.

| Gene | Chr | Functional relevance | MBMS |  | MESA |  |  |
| --- | --- | --- | --- | --- | --- | --- | --- |
|  |  |  | Black NH | White NH | Black NH | White NH | Hispanic |
| ZNF286B | 17 | A pseudogene, which is predicted to be involved in regulation of RNA polymerase 2 (Pol II)-mediated transcription (Pol II transcribes protein-coding genes into mRNA <sup>41</sup>). | Born in a Jim Crow state: 1 |  |  |  |  |
| PLXND1 | 3 | Encodes a cell receptor involved in axonal guidance, migration of endothelial cells, and regulates atherosclerotic plaque deposition <sup>42</sup>. | <HS education: 1 |  |  |  |  |
| FUT6 | 19 | A Golgi stack membrane protein that is involved in basophil-mediated allergic inflammation <sup>43</sup>. |  |  | residential racialized economic segregation: 1 |  |  |
| KLF6 | 10 | A transcriptional activator and tumour suppressor, which regulates macrophage inflammatory responses <sup>44</sup>. |  |  | LAC: 2<br>NOx: 3 | LAC: 1<br>NOx: 1 |  |
| FOS | 14 | FOS is an early-response gene, and is a subunit of the AP-1 transcription factor complex, which regulates gene expression involved in lung injury, repair and transformation <sup>45</sup>, as well as regulating many cytokine genes and T-cell differentiation <sup>46,47</sup>. |  |  | LAC: 1<br>NOx: 7 |  |  |
| FOSB | 19 | FOSB is another subunit of AP-1. |  |  | LAC: 1<br>NOx: 1 |  |  |
| ZFP36 | 19 | ZFP36 encodes a protein (TTP) that is a key regulator of post-transcriptional regulation, which has roles in immune and inflammatory responses <sup>48</sup>. |  |  | NOx: 2 |  |  |
| DUSP1 | 5 | DUSP1 is a gene that regulates airway inflammation; DUSP1's key mechanism of inflammation modulation may be via modulating the actions of the protein TTP encoded by the ZFP36 gene <sup>49</sup>. |  |  | LAC: 1<br>NOx: 1 |  |  |
| VIM | 10 | Encodes a filament protein responsible for integrity of cell shape and cytoplasm. Pathogens can attach to this protein on the cell surface. Putative involvement regulating innate immune response to lung injury and irritation <sup>50</sup> |  |  | NOx: 1 |  |  |
| PDE4D | 5 | PDE4s, including PDE4D, have roles in cell signalling, as well as regulating inflammatory responses<sup>51</sup>. |  |  | MDS: 1 |  |  |
| MALAT1 | 11 | Metastasis associated lung adenocarcinoma transcript 1, lncRNA that acts as transcriptional regulator; upregulation linked to cancerous tissues and proliferation and metastasis of tumour cells |  |  |  | LAC: 1 |  |
| CYTIP | 2 | Modulates activation of ARF (ADP-ribosylation factor) genes, which regulate vesicle budding, tethering and cytoskeleton organization. Dysregulation of ARFs may be involved in cancer cell migration and invasion. |  |  |  | LAC: 1<br>NOx: 1 |  |
| ZEB2 | 2 | DNA-binding transcriptional repressor involved in the transforming growth factor-β (TGF-β) signalling pathway that interacts with activated SMADs. May be related to small cell lung cancer<sup>52</sup>. | | | | LAC: 1<br>NOx: 1 | |
| PTPRC | 1 | A receptor-type PTP that is an essential regulator of T- and B-cell antigen receptor signalling. |  |  |  | LAC: 1<br>NOx: 1 |  |
| NPNT | 4 | An extracellular matrix protein that has roles in kidney development and carcinogenesis<sup>53</sup>. |  |  |  |  | LAC: 1 |
| ADPRHL1 | 13 | a protein encoding a pseudoenzyme involved in cardiogenesis<sup>54</sup>. |  |  |  |  | NOx: 1 |

**Table 3:** Summary of the number of DNAm sites passing the genome-wide threshold in each individual EWAS in MBMS (threshold 2.4e-7) and MESA (threshold 9e-8). The list of specific DNAm sites passing the genome-wide threshold can be found in supplementary table 4. ^1^The EWAS was not run for Jim Crow birth state for white NH participants in MBMS, due to small cell numbers. ^2^See text; these 53 sites were driven by air pollution differences between individuals born and not born in a Jim Crow state. ^3^The two EWAS for parental education were not run for Hispanic participants in MESA, due to small cell numbers. ^4^The EWAS was not run for MDS (score of 1-2 vs 3+) for white NH participants in MESA, as no participants had a score of 3 or more.

|  | MBMS |  |  |  | MESA |  |  |  |  |  |
| --- | --- | --- | --- | --- | --- | --- | --- | --- | --- | --- |
|  | Black NH |  | white NH |  | Black NH |  | white NH |  | Hispanic |  |
|  | N | N sites | N | N sites | N | N sites | N | N sites | N | N sites |
| Birth in a Jim Crow state | 224 | 1 | NA<sup>1</sup> | NA<sup>1</sup> | 225 | 0 | 549 | 53<sup>2</sup> | 186 | 0 |
| Parent's highest education (high vs low) | 64 | 0 | 33 | 0 | 117 | 0 | 288 | 0 | NA<sup>3</sup> | NA<sup>3</sup> |
| Parent's highest education (high vs mid) | 129 | 0 | 49 | 0 | 128 | 0 | 384 | 0 | NA<sup>3</sup> | NA<sup>3</sup> |
| Participant's education (high vs low) | 63 | 1 | 36 | 0 | 54 | 0 | 142 | 0 | 51 | 0 |
| Participant's education (high vs mid) | 190 | 0 | 61 | 0 | 202 | 0 | 534 | 0 | 156 | 0 |
| Household poverty to income ratio | 190 | 0 | 66 | 0 | 218 | 0 | 528 | 0 | 180 | 0 |
| Racialized economic segregation | 224 | 0 | 69 | 0 | 223 | 1 | 545 | 1 | 174 | 0 |
| Black carbon | 224 | 0 | 69 | 0 | 211 | 12 | 526 | 7 | 175 | 1 |
| Nitrogen oxides | 219 | 0 | 69 | 0 | 211 | 22 | 526 | 4 | 175 | 1 |
| EOD<sup>5</sup> (1-2 vs 0) | 82 | 0 | 59 | 0 |  |  |  |  |  |  |
| EOD<sup>5</sup> (1-2 vs 3+) | 192 | 0 | 34 | 0 |  |  |  |  |  |  |
| MDS<sup>6</sup> (1-2 vs 0) |  |  |  |  | 204 | 1 | 554 | 0 | 184 | 0 |
| MDS<sup>6</sup> (1-2 vs 3+) |  |  |  |  | 97 | 0 | NA<sup>4</sup> | NA<sup>4</sup> | 60 | 0 |

### MESA subgroup analysis

The main impact of removing the Minnesota and Forsyth County sites (which both had very low levels of air pollution) was to remove the confounding structure between air pollution and Jim Crow birth state among white NH participants. It also increased the similarity of air pollution associations between the Black NH and white NH participants; for example, of the 19 DNAm sites associated with NOx among white NH participants, 12 passed the genome-wide threshold in the Black NH participant EWAS. Numbers of associated sites are in Table 4. Miami plots for this MESA subgroup can be found in Supplementary figures 6, 7 and 8.

**Table 4:** Summary of EWAS results for MESA subgroup analysis. ^1^The EWAS for Jim Crow birth state was not run for Hispanic participants due to small cell numbers. ^2^The EWAS for parental and participant education were not run for Hispanic participants, due to small cell numbers. ^3^The EWAS was not run for MDS (score of 1-2 vs 3+) for white NH and Hispanic participants in MESA, as no participants had a score of 3 or more.

|  | Black NH |  | white NH |  | Hispanic |  |
| --- | --- | --- | --- | --- | --- | --- |
|  | N | N sites | N | N sites | N | N sites |
| Birth in a Jim Crow state | 221 | 0 | 237 | 0 | NA<sup>1</sup> | NA<sup>1</sup> |
| Parent's highest education (high vs low) | 115 | 0 | 134 | 0 | NA<sup>2</sup> | NA<sup>2</sup> |
| Parent's highest education (high vs mid) | 125 | 0 | 164 | 0 | NA<sup>2</sup> | NA<sup>2</sup> |
| Participant's education (high vs low) | 54 | 0 | 55 | 0 | NA<sup>2</sup> | NA<sup>2</sup> |
| Participant's education (high vs mid) | 198 | 0 | 227 | 0 | NA<sup>2</sup> | NA<sup>2</sup> |
| Household poverty:income ratio | 214 | 0 | 227 | 1 | 67 | 0 |
| Racialized economic segregation | 219 | 1 | 233 | 0 | 59 | 0 |
| Light Absorption Coefficient | 208 | 10 | 231 | 6 | 57 | 0 |
| Nitrogen oxides | 208 | 20 | 231 | 19 | 57 | 0 |
| Major Discrimination Scale (1-2 vs 0) | 200 | 1 | 236 | 0 | 67 | 0 |
| Major Discrimination Scale (1-2 vs 3+) | 94 | 0 | NA<sup>3</sup> | NA<sup>3</sup> | NA<sup>3</sup> | NA<sup>3</sup> |

### Meta-analysis

Meta-analysis in MBMS did not yield any sites passing the genome-wide threshold. In MESA we see approximately similar numbers of associations as with the Black NH subgroup (17 for LAC and 18 for NOx); see Supplementary Table 3. When we restricted to participants recruited at the Baltimore and New York sites, a much larger number of DNAm sites passed the genome-wide threshold (51 for LAC and 79 for NOx); this may be because Minnesota and Forsyth County sites had very low variance in pollution levels. The MESA sensitivity meta-analysis identified multiple associations linked to DUSP1, FOS, KLF6, MCL1, and VIM; genes that have putative roles in inflammation and immunity.

## Biological enrichments of exposure associations

### Gene ontology

We observed no evidence for gene set enrichments for any Gene Ontology terms among the top 100 sites of the main EWAS we conducted. However, we did observe that the 22 sites associated with NOx above the genome-wide threshold among MESA Black NH participants were enriched for the gene ontology terms ‘response to glucocorticoid’ and ‘response to corticosteroid’ (FDR>0.05). We also observed that the MESA meta-analysis of NOx among all participants was associated with 13 Gene Ontology terms (FDR>0.05) related to blood-based immune response.

### EWAS catalog

We observed a number of relevant enrichments among sites identified in our EWAS. Details of the associations (p < 0.05, Fisher’s exact test) can be found in the Supplementary Materials and Supplementary figures 6-16. Briefly, in MBMS, we see enrichment for inflammation for both NOx and LAC EWAS among Black NH participants, and in the NOx meta-analysis. In MESA, we observed consistent enrichment for infection and cancer among Black NH and white NH participants, and also in the meta-analyses. We observed enrichment for inflammation among Hispanic participants. We also found among both MESA Black NH and Hispanic participants, the racialized economic segregation EWAS was enriched for neurological traits. Among both the white NH and Hispanic participants, household poverty to income ratio EWAS was enriched for SEP and education. In the MESA subgroup analysis, enrichment for prenatal exposures was observed for the Jim Crow birth state EWAS among the Black NH and Hispanic participants.

### Enrichment for genomic features

When we looked at enrichment of genomic locations of the top 100 sites (p < 0.05, Fisher’s exact test), we found that among MBMS Black NH participants, NOx was the only exposure with associated CpGs being located in active genomic regions (please see Supplementary materials for details). In the MBMS meta-analyses, NOx was enriched for regions related to gene promoters. Among MESA Black NH participants, we observe enrichment for regions related to transcription and genome regulation in the LAC and NOx EWAS. We also observed enrichment relating to transcription regulation for the birth in a Jim Crow state EWAS. Among MESA white NH participants, we observed enrichment for transcription regulation for both measures of air pollution. Among MESA Hispanic participants, LAC exposure shows some associations with active genomic regions. When we restrict MESA to the New York and Baltimore sites, we see a similar set of enrichments; and in the MESA meta-analyses we see consistent enrichment related to transcription regulation and promotors. Notably, genomic feature enrichments for NOx among both MBMS and MESA Black NH participants involved similar genomic locations (CpG islands and shores) and chromatin states (related to promotors), as well as 6 of a possible 9 TFBS.

### Lookup of associations in a priori specified genomic locations

We did not observe any associations in our EWAS results for sites identified in previous EWAS of related exposures.

## Discussion

The series of EWAS we conducted on a range of adverse exposures at different levels and at different points in the lifecourse, drawing on two different population-based studies with similar exposure data, provide evidence that DNAm may be a biological pathway by which societal context shapes health inequities. This work has shown for the first time associations between DNAm and multiple levels of structural discrimination, in genes that are biologically plausible routes of embodiment involving gene regulation, including inflammation. Additionally, our EWAS and meta-analyses of air pollution showed clear association between two road traffic-related measures of air pollution, and DNAm of multiple CpGs in multiple genes that have been consistently associated with inflammation and infection, suggesting that the material environment people live may induce inflammatory changes. Our study has added to the existing literature on air pollution; there are few EWAS studies looking at NOx (n=3), and none so far looking at black carbon. In total, this work highlights the need for researchers to consider multiple levels of discrimination and adversity across the lifecourse, especially structural inequities in the material world in which people live, to fully elucidate drivers and biological mechanisms of inequitable health.

Associations detected at the genome-wide level in MBMS related more closely to early-life exposures (being born in a Jim Crow state, and low educational attainment); in MESA they related more to current experiences and exposures (air pollution, racialized economic segregation, and experiences of discrimination), possibly reflecting the relatively older age of the MESA participants. The much stronger associations with air pollution in MESA compared to MBMS could potentially be due to: (1) the use of purified monocytes in MESA, with a single cell type making associations easier to detect; (2) less variation in exposure to air pollution in MBMS compared MESA; (3) longer duration of air pollution exposure in MESA (due to older age of the participants); or (4) reduced statistical power in MBMS, due to lower quantities of DNA.

Notably, inflammation was the predominant pathway indicated in the air pollution analyses, both via putative gene functions and enrichment analyses. These findings underscore that while there is a large psychosocial literature on inflammation being a mechanism by which discrimination harms health ^28,55,56^, it is also critical to consider inequities in biophysical exposures in the material world as an important driver of this inflammation. Overall, air pollution sites tend to be enriched for inflammation in MBMS and infection in MESA; this could represent different mechanisms of the same process due to the different blood cell types sampled in the two cohorts; with monocytes being specialised in infection prevention, and neutrophils (the highest proportion cell in whole blood) being specialised in inflammatory responses.

Our study identified a greater number of associations with air pollution measures than previous work in MESA ^20,57^; this is likely due to the fact that we do not adjust for recruitment site (which would reduce variation in the exposure because exposure is location-dependent); and previous analyses have adjusted for racialized group membership, which is also associated with air pollution exposure; this may have masked the effects that we have detected. This joins other research that has demonstrated the importance of considering spatial effects of air pollution ^58^.

A limitation of our study is that we cannot infer causality. Although it would be possible to conduct Mendelian randomization instrumenting cis-mQTLs, we did not conduct this analysis because we think the results would be highly speculative. Additionally, the MESA sample we used may have been subject to selection bias, because (1) individuals who had experienced prior cardiovascular events were excluded from recruitment, and (2) a number of participants died between Exam 1 and Exam 5. If adversity and discrimination are associated with these cardiovascular events and mortality, associations could be biased in MESA.

## Conclusions

We think this work provides direction for future epigenetic studies to consider the role of inequitable adverse social and biophysical exposures across the lifecourse, including but not limited to structural discrimination. Our results suggest inflammation may be a key biological pathway by which inequities become embodied, in our case driven primarily by exposure to air pollution, and not self-reported racial discrimination. These findings accordingly suggest that attention to how social inequities shape biophysical as well as social exposures is crucial for understanding how societal inequities can become embodied, via pathways involving DNAm.

## Declaration statements

### Ethics approval

The study protocol, involving use of both the MBMS and MESA data, was approved by the Harvard T.H. Chan School of Public Health Office of Human Research Administration (Protocol # IRB19-0524; June 10, 2019).

The original MBMS study protocol, implemented in accordance with the Helsinki Declaration of 1975, as revised in 2000, was approved by the Harvard School of Public Health Office of Human Research Administration (protocol #11950–127, which covered 3 of the 4 health centers through reciprocal IRB agreements), and was also separately approved by the fourth community health center’s Institutional Review Board. All participants provided written informed consent.

Information regarding the MESA protocols and their IRB approvals and other information, is available at: www.mesa-nhlbi.org.

### Data sharing

This study (<u>NIH Grant number R01MD014304</u>) relied on three sources of data, each of which is subject to distinct data sharing stipulations: (1) the non-public data from the “My Body, My Story” (MBMS) study; (2) the non-public data from the Multi-Ethnic Study of Atherosclerosis (MESA; data use agreement G638); and (3) the public de-identified data from the US Census, the American Community Survey, and the State Policy Liberalism Index. We provide descriptions of these data sharing stipulations and access to these data below; this information is also available at: https://www.hsph.harvard.edu/nancy-krieger/data-sharing-resources/

- ICE metrics relating to racial composition, income distribution, and housing tenure that were derived from sources in the public domain i.e. the US Census and the American Community Survey are available at the census tract level now on <u>GitHub</u>.
- Code used to construct the variables is available on <u>GitHub</u> and <u>here</u>.
- The State Policy Liberalism Index data used in our study is also publicly available and can be obtained from the <u>Harvard Dataverse</u>. Reference: Caughey, Devin; Warshaw, Christopher, 2014, “The Dynamics of State Policy Liberalism, 1936-2014”, http://dx.doi.org/10.7910/DVN/ZXZMJB Dataverse [Distributor] V1 [Version].
- De-identified data from the My Body My Story study used for this project will be made available only for purposes approved by the study PI, as stipulated by the study’s informed consent protocol. The application form to obtain these data will be made available via this website after completion of this project in late Fall 2024.
- Data from the <u>Multi-Ethnic Study of Atherosclerosis (MESA)</u> must be obtained directly from the MESA website via their application protocol.
- The scripts to run the EWAS and downstream analyses are available on <u>GitHub</u>.
- EWAS summary statistics will be uploaded to the <u>EWAS catalog</u> website upon publication.

### Author contributions

- SHW performed quality checks and normalisation of MBMS DNAm data, co-designed and conducted the analyses, wrote the first manuscript draft, produced tables and figures, and prepared study materials to be shared via the data repository (software code).
- MS provided advice on data QC, co-designed the analyses, and contributed to interpreting the results.
- CR co-led obtaining funds for the research project, co-designed the analyses, and contributed to interpreting the results.
- NK led conceptualization of the study, contributed to designing the analyses, and co-led obtaining funds for the research project.
- CT accessed the electronic public use data and generated the study variables derived from these data, contributed to designing the analyses and interpreting the results, and prepared the study materials to be shared via the data repository (data dictionary; software code).
- JTC contributed to designing the analyses and interpreting results.
- PDW facilitated finalizing all human subject approvals and data use agreements and also the data transfer of the MBMS epigenetic data from HSPH to Bristol, geocoded the place of birth data, extracted the historical census data from PDF files, and contributed to interpreting results.
- AS, BC, KT, and GDS contributed to designing the analyses and interpreting the results.
- IDV led and supervised the assays to extract epigenetic data from the MBMS blood spots and contributed to designing the analyses and interpreting the results.
- ADR facilitated interpretation of the MESA data and contributed to designing the analyses and interpreting the study results.
- All co-authors provided critical intellectual content to and approved the submitted manuscript.

## Acknowledgements

We thank, with written permission: (1) Nykesha Johnson for her insightful comments on this manuscript (2) Linda V. Nguyen (Department of Medicine, Brigham and Women’s Hospital and Harvard Medical School, Boston, MA, USA; May 1, 2023) for her contributions to extracting the DNA from the MBMS blood spots and preparing it for shipment to the Bristol team members for analysis, (3) Emily Wright, PhD (Harvard TH Chan School of Public Health; May 1, 2023), for her assistance, when a doctoral student, with linking area-based data to the MBMS study participant records; and (4) Karen Ho (May 17, 2023), (5) Louise Falk (May 18, 2023), (6) Sophie FitzGibbon (May 24, 2023), and (7) Catherine Slack (May 19, 2023) for their role in generating the MBMS Illumina data.

We are extremely grateful to all the MBMS participants who took part in the original MBMS study, the staff at the four participating community health centers, and the researchers who conducted participant recruitment and interviews. We are likewise grateful to all the MESA participants and investigators whose data and dedicated work enabled us to have incorporate MESA data into our research project, including Amanda Gassett, MS for providing us with the air pollution dataset used from MESA Air (written permission: July 20, 2023); a full list of participating MESA investigators and institutions can be found at: http://www.mesa-nhlbi.org. This paper has been reviewed and approved by the MESA Publications and Presentations Committee (DATE).

