## Supplementary figures for "An epigenome-wide analysis of DNA methylation, racialized and economic inequities, and air pollution"

### Miami plots

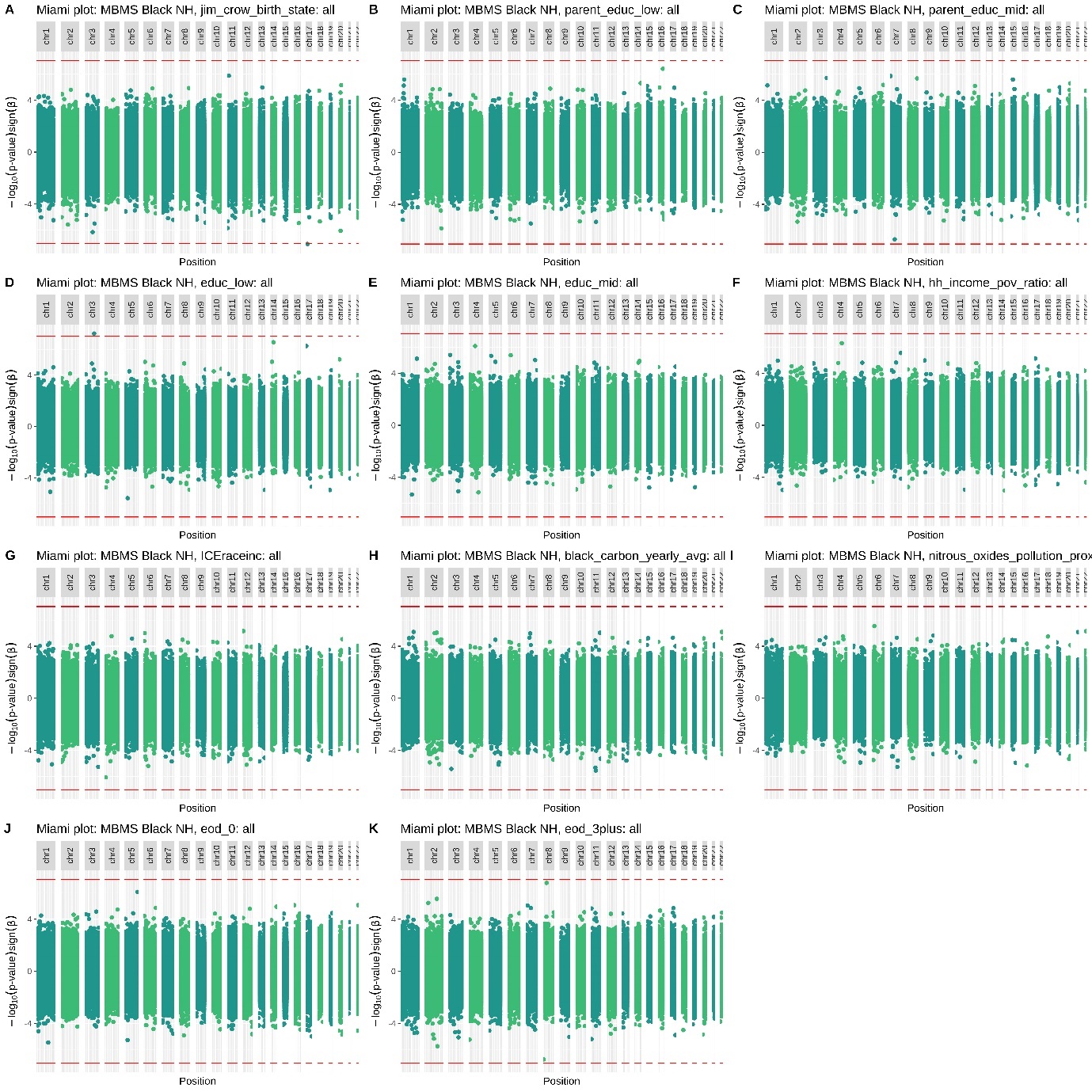

Figure 1: MBMS Black NH Miami plots

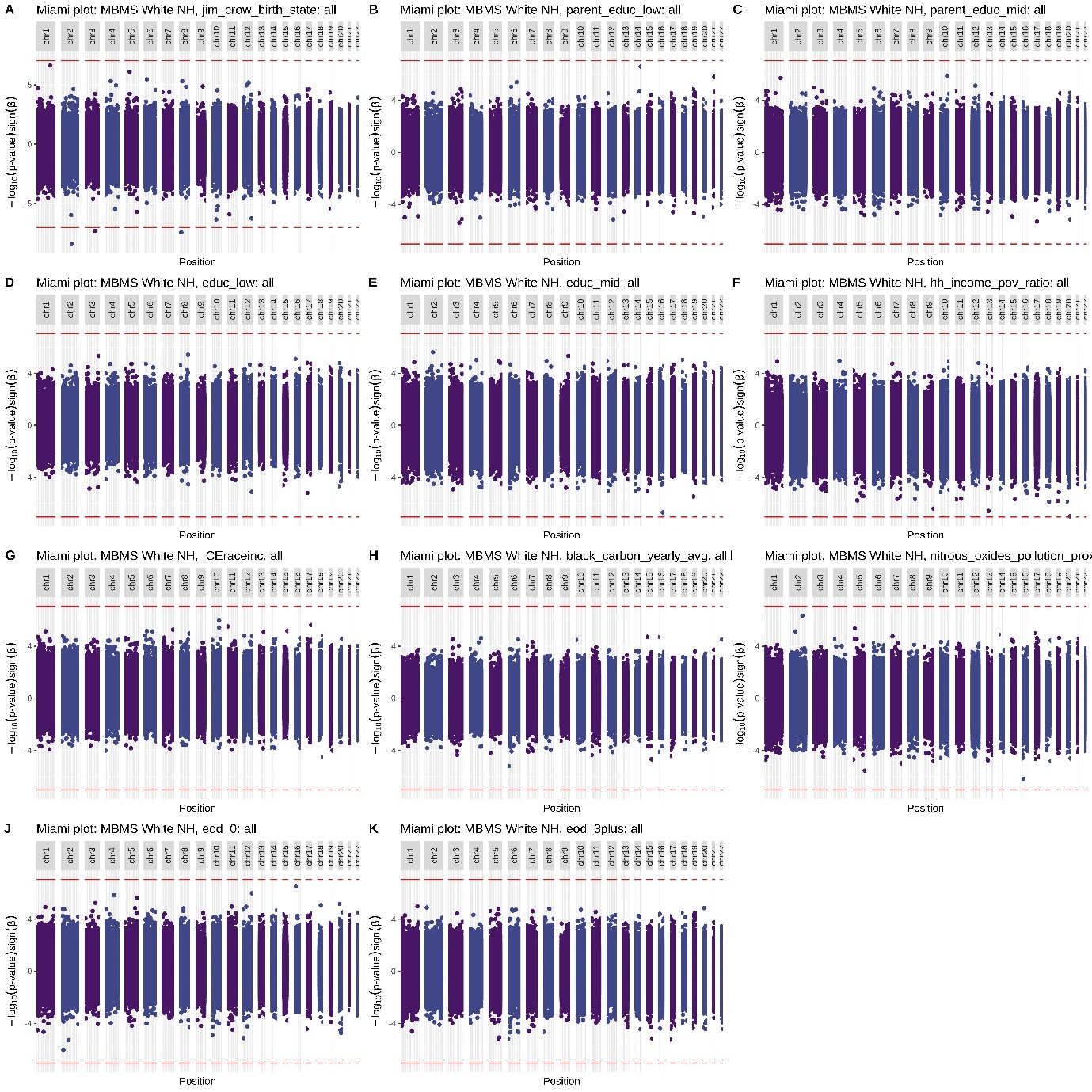

Figure 2: MBMS white NH Miami plots

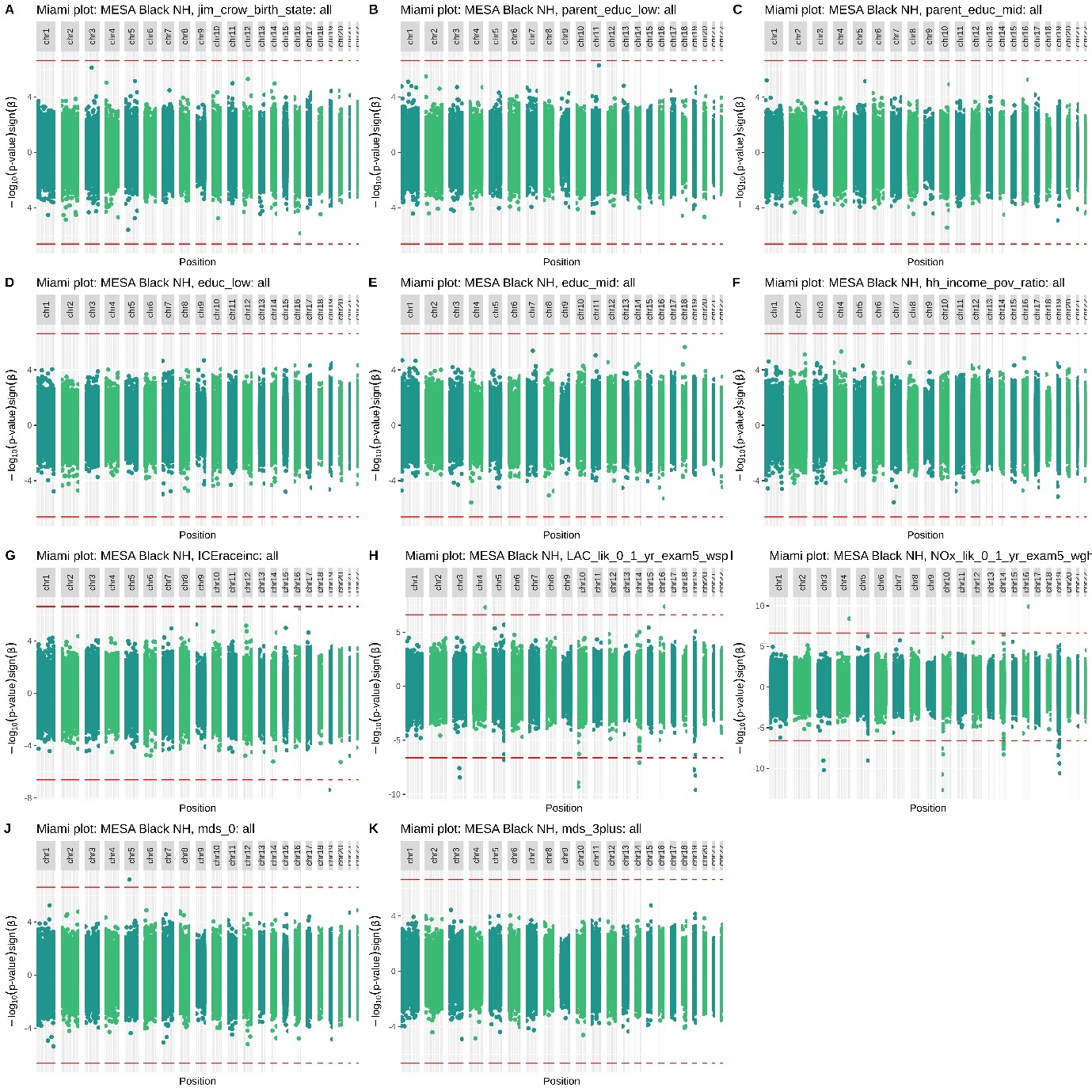

Figure 3: MESA Black NH Miami plots

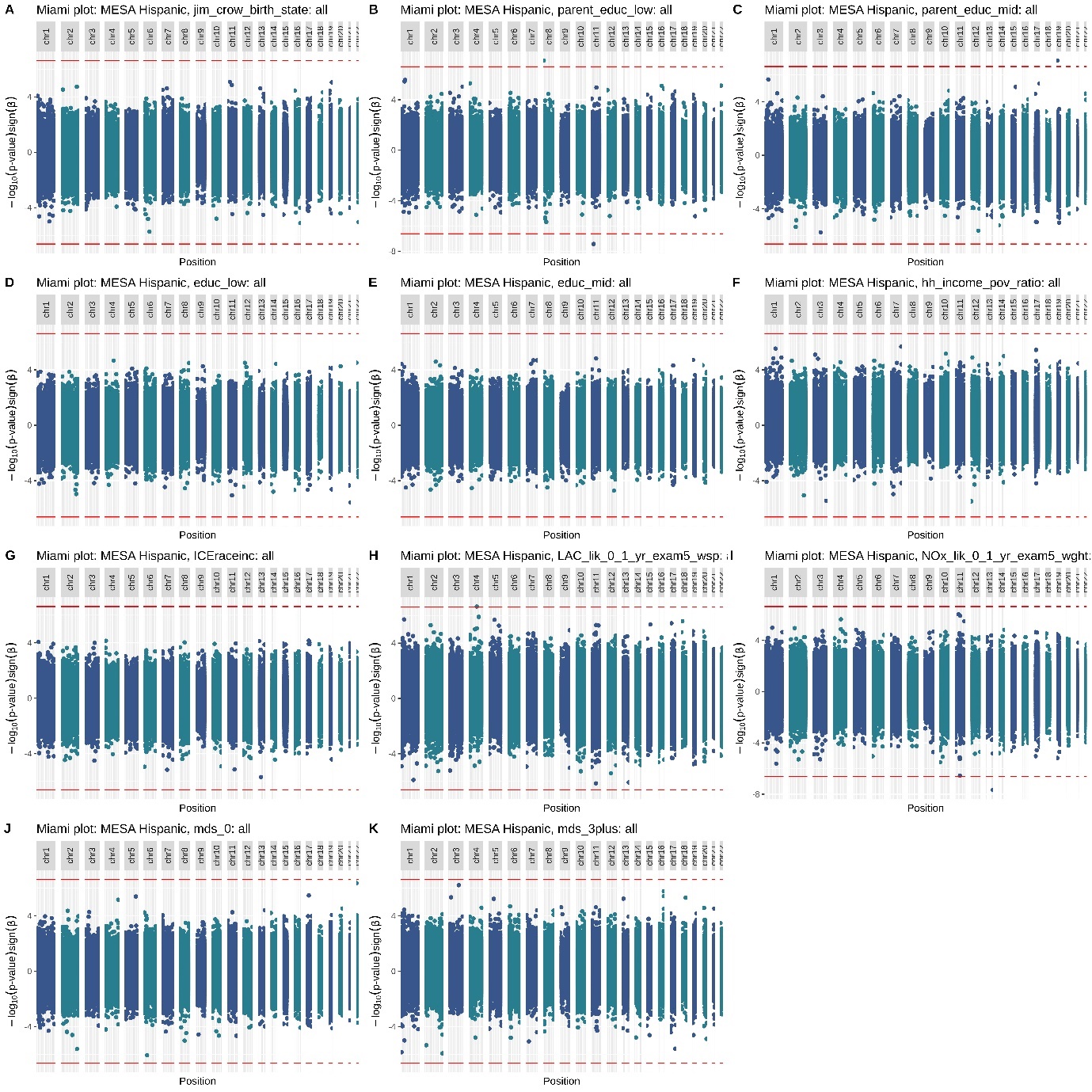

Figure 4: MESA Hispanic Miami plots

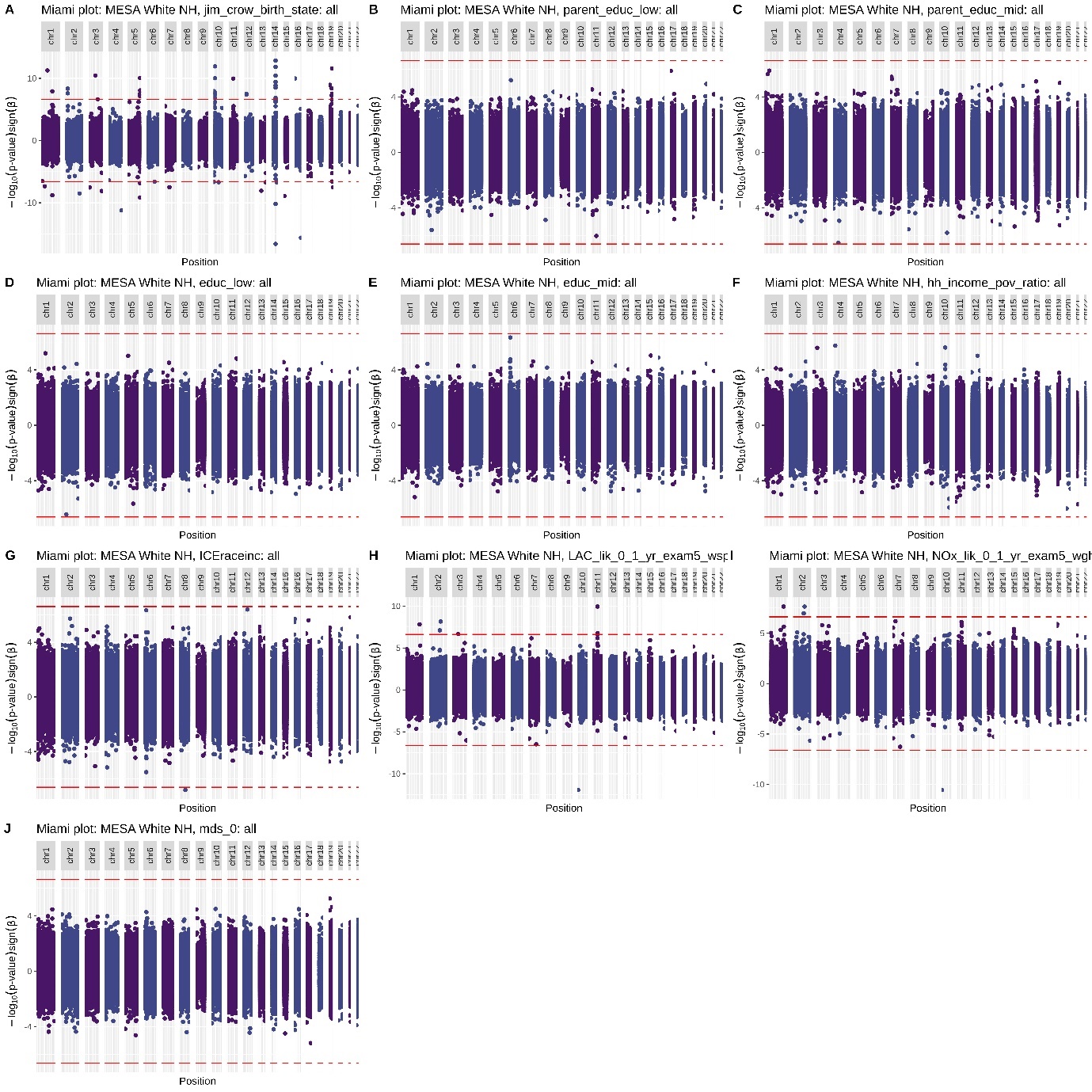

Figure 5: MESA white NH Miami plots

### MESA New York and Baltimore subset Miami plots

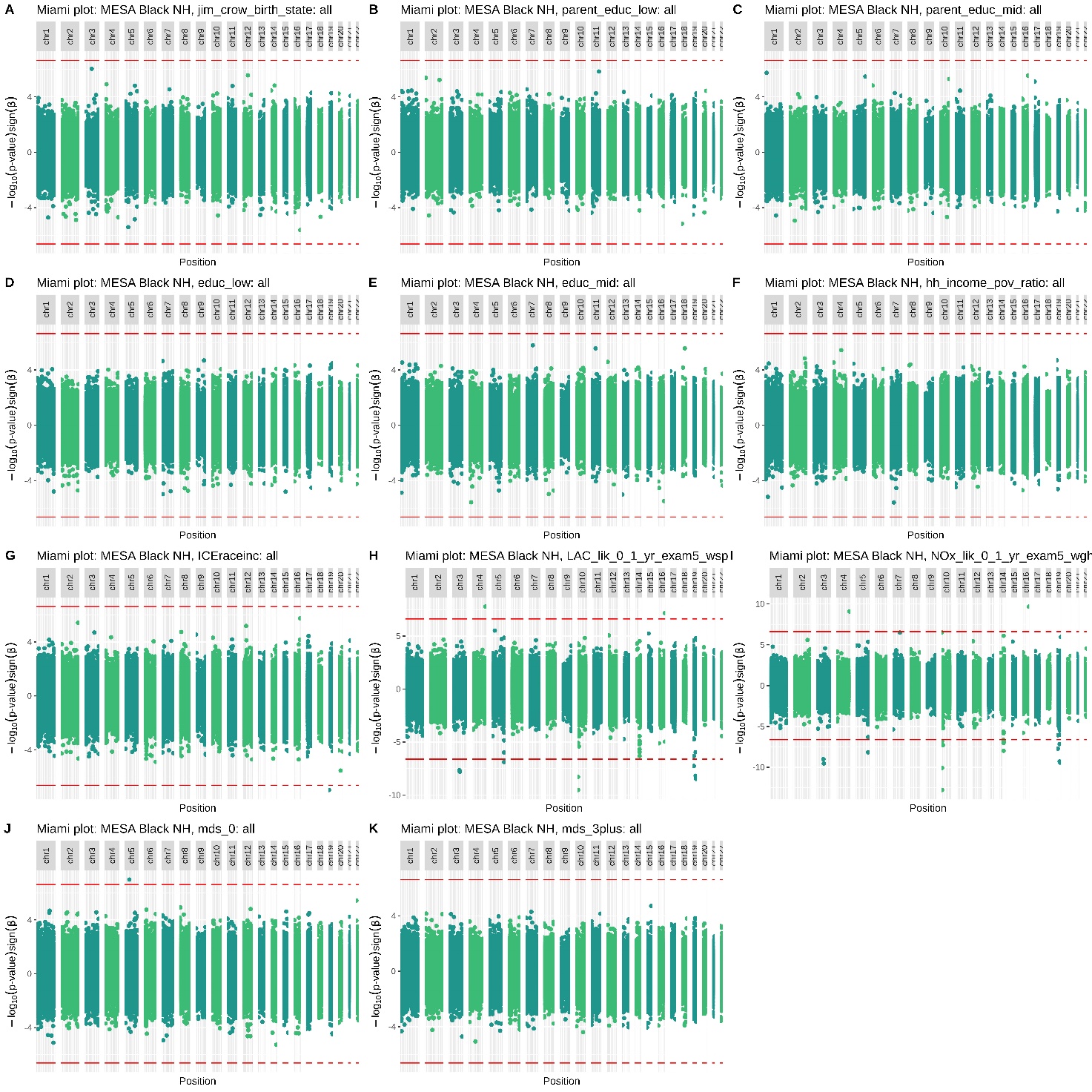

Figure 6: MESA Black NH Miami plots (JHU and COL subgroup)

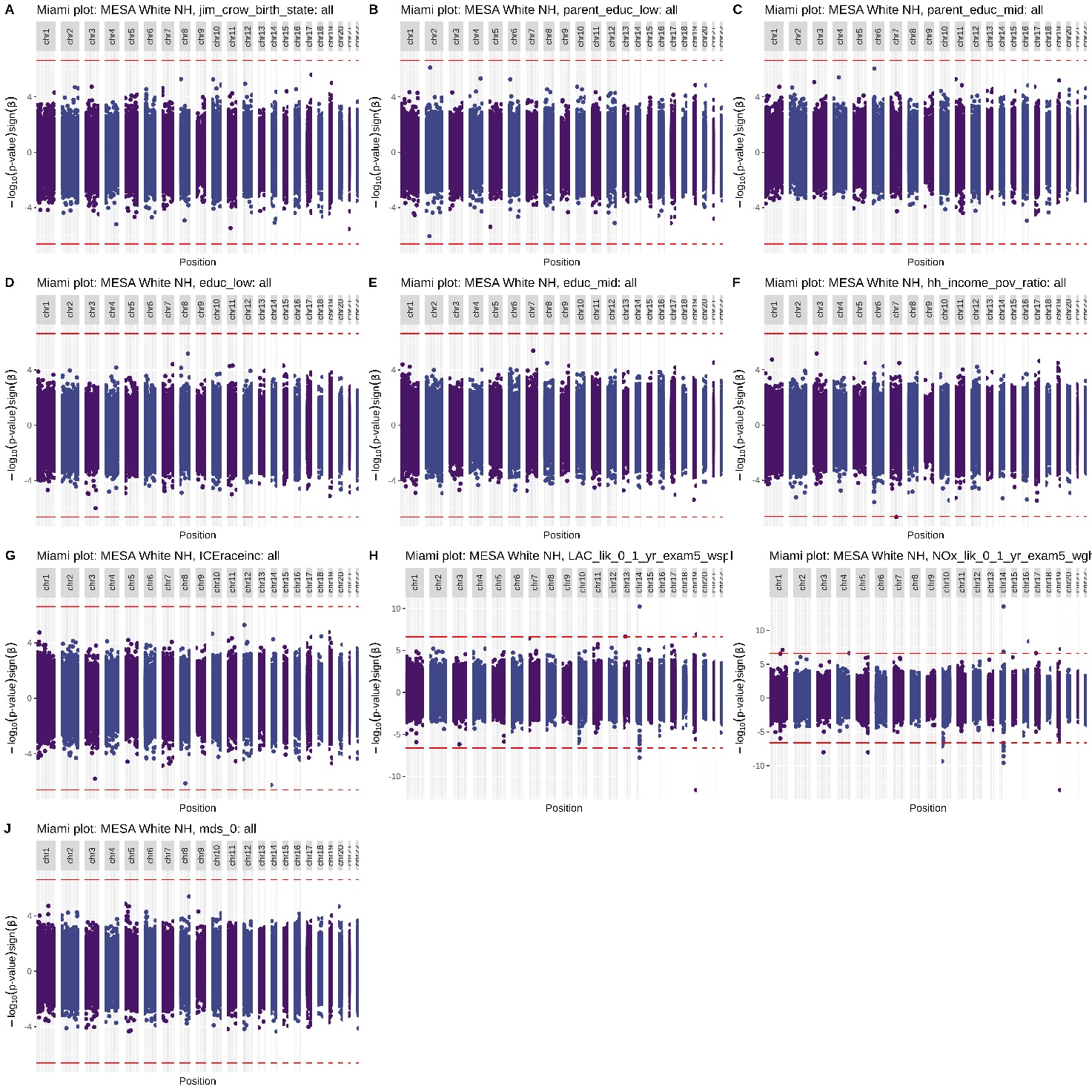

Figure 7: MESA white NH Miami plots (JHU and COL subgroup)

Figure 8: MESA white NH Miami plots (JHU and COL subgroup)

### EWAS catalog enrichment plots

#### MBMS full cohort

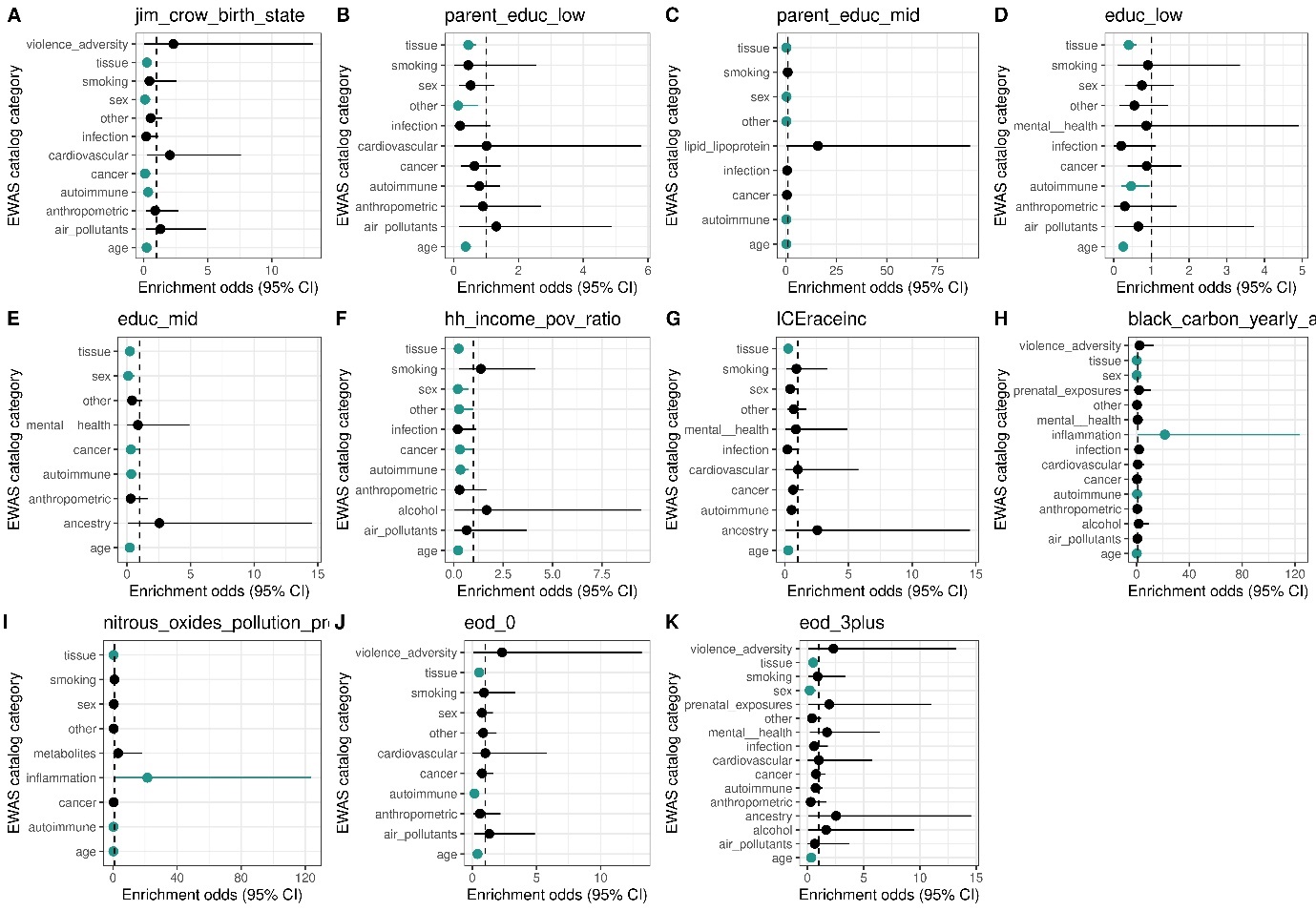

Figure 9: EWAS catalog enrichment plot: MBMS Black NH

----------------------------------------------------------------------

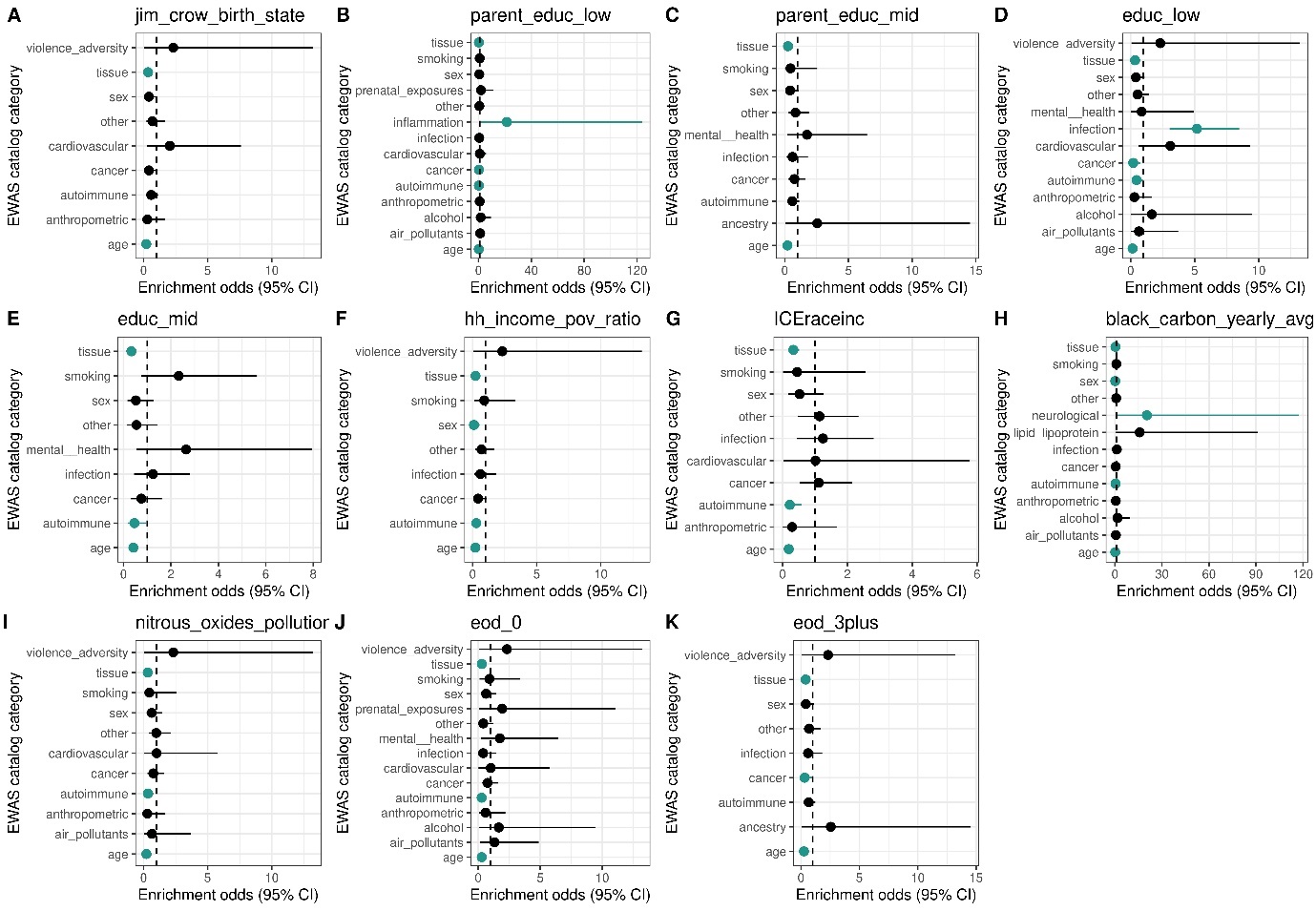

Figure 10: EWAS catalog enrichment plot: MBMS white NH

----------------------------------------------------------------------

#### MESA full cohort

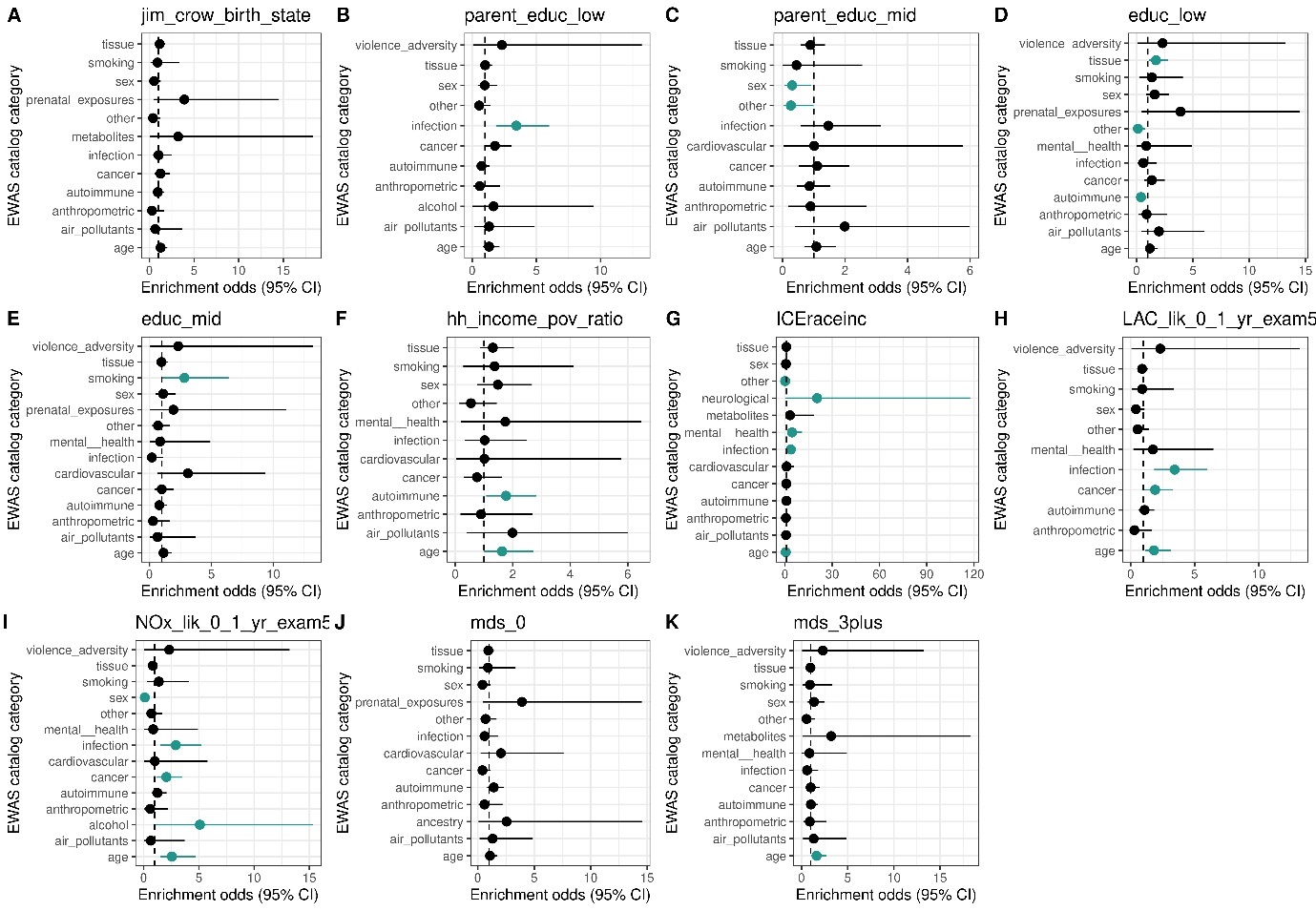

Figure 11: EWAS catalog enrichment plot: MESA Black NH

----------------------------------------------------------------------

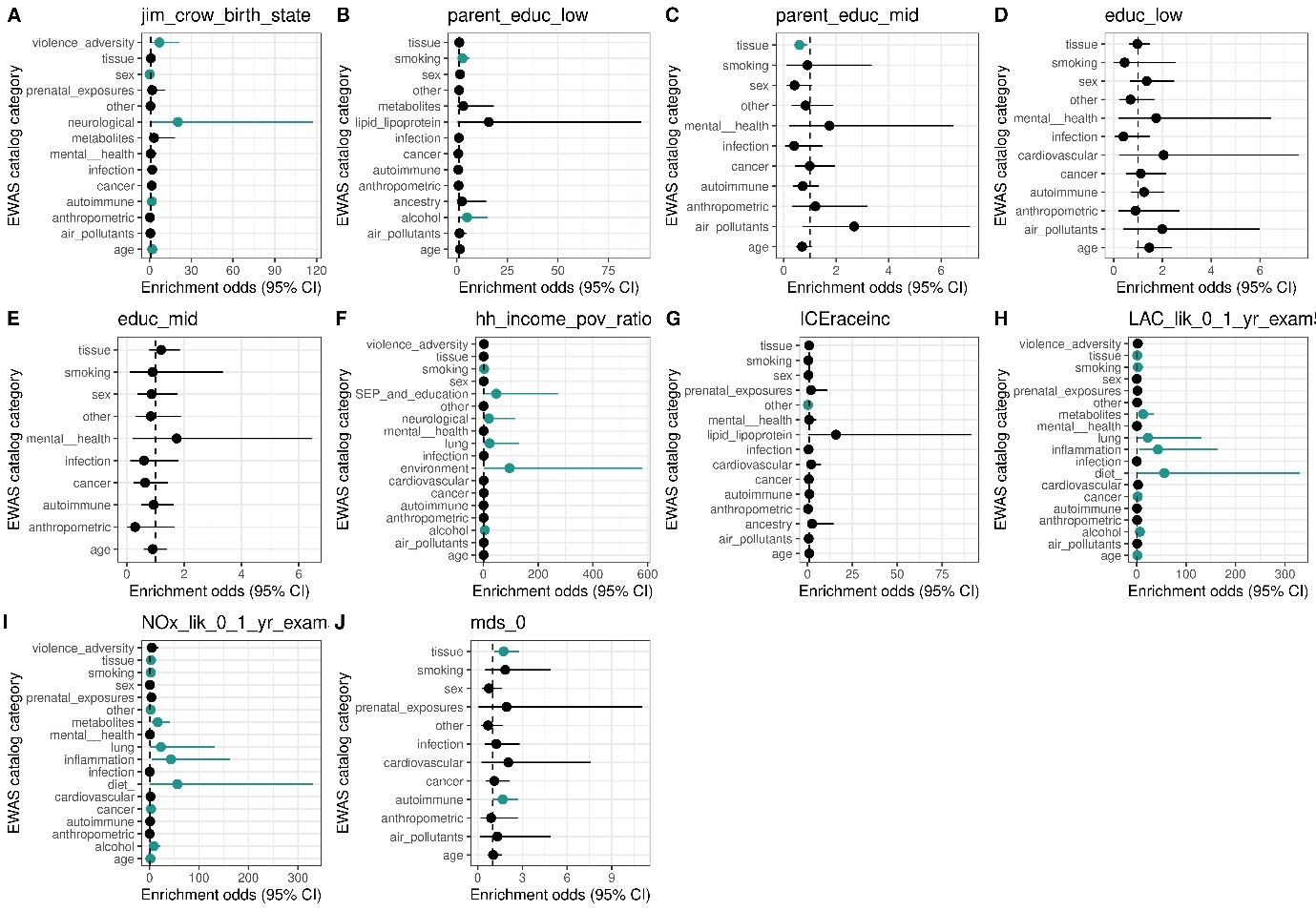

Figure 12: EWAS catalog enrichment plot: MESA white NH

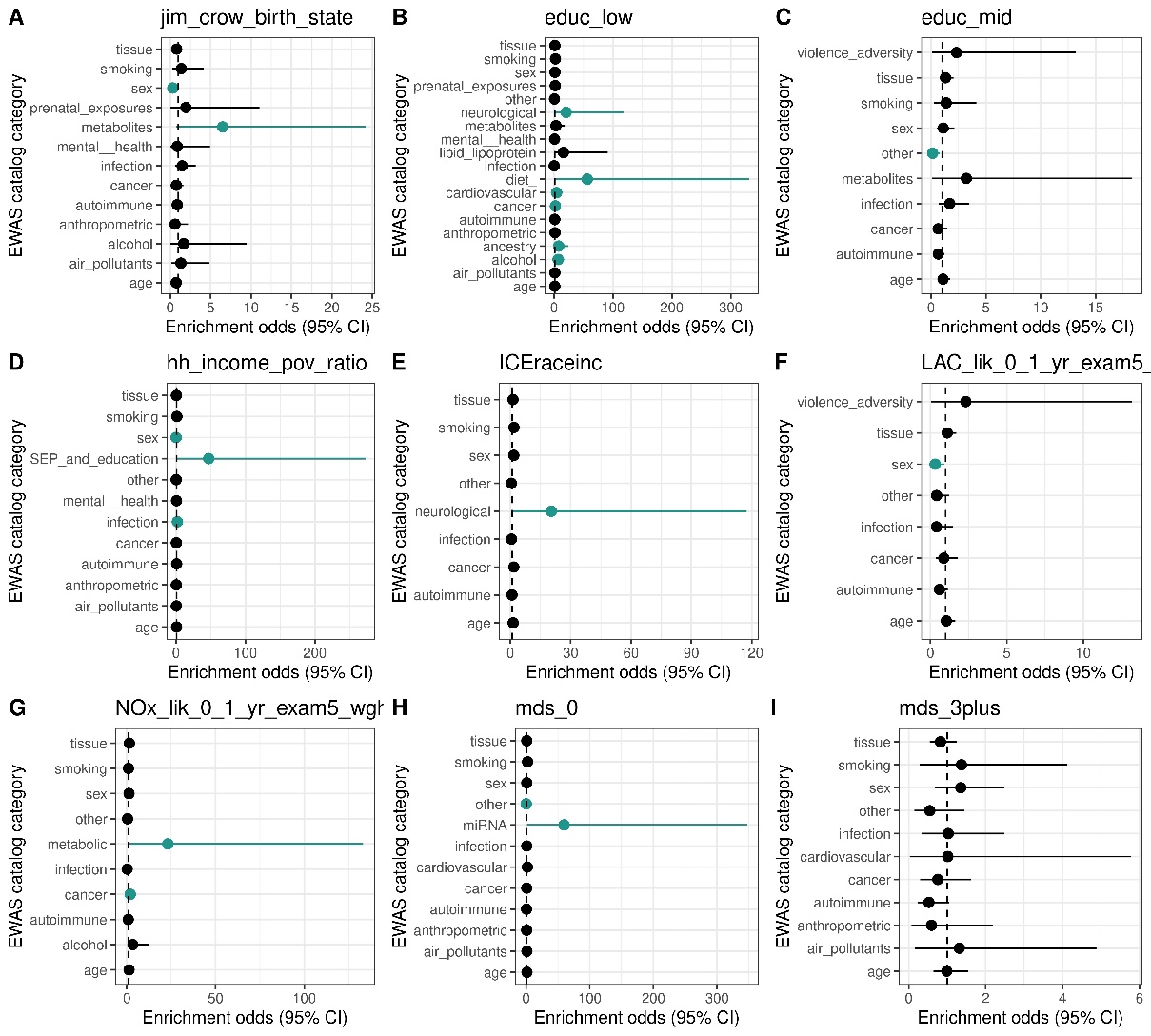

Figure 13: EWAS catalog enrichment plot: MESA Hispanic

----------------------------------------------------------------------

#### MESA JHU/COL subgroup

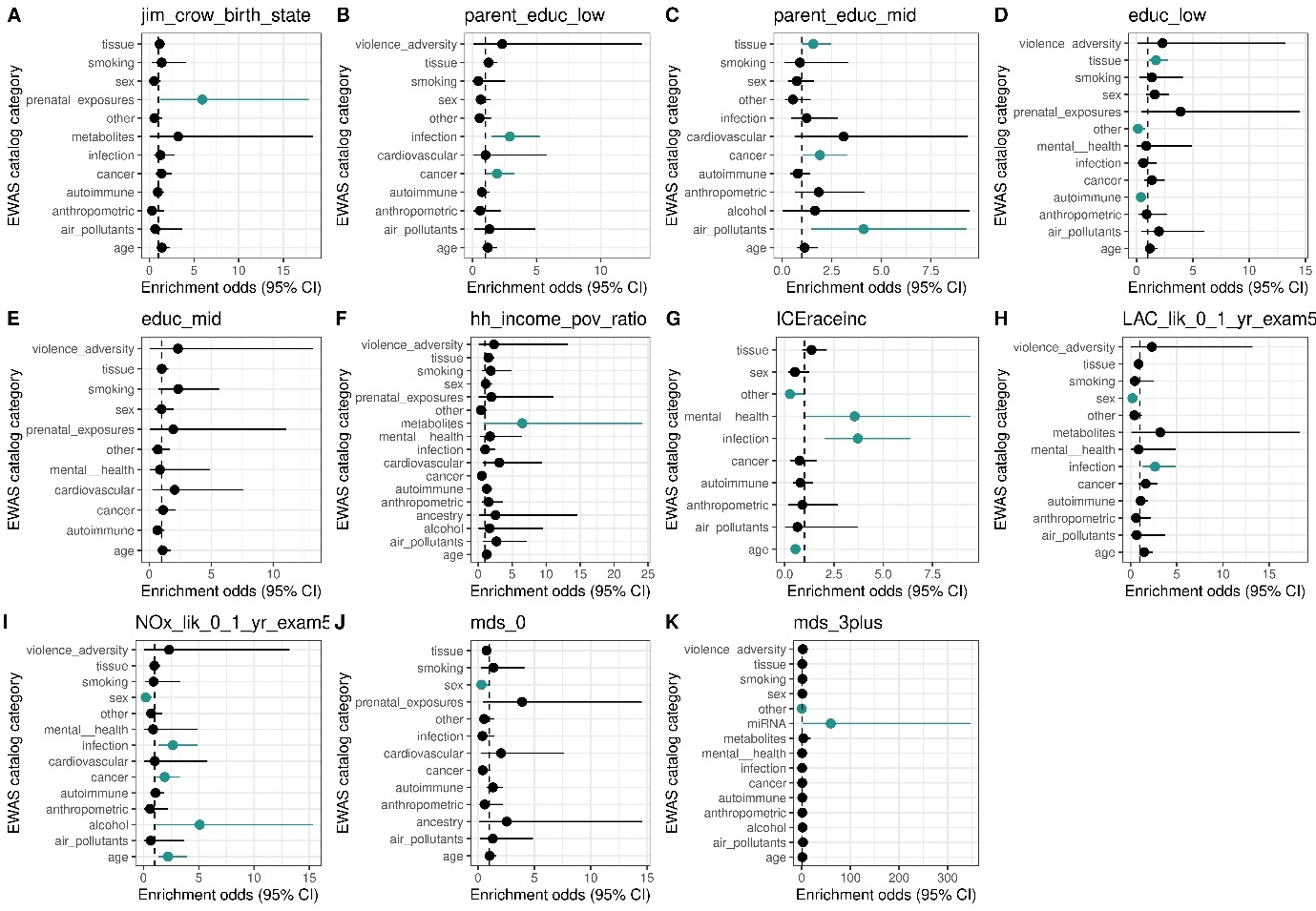

Figure 14: EWAS catalog enrichment plot: MESA Black NH JHU + COL subgroup

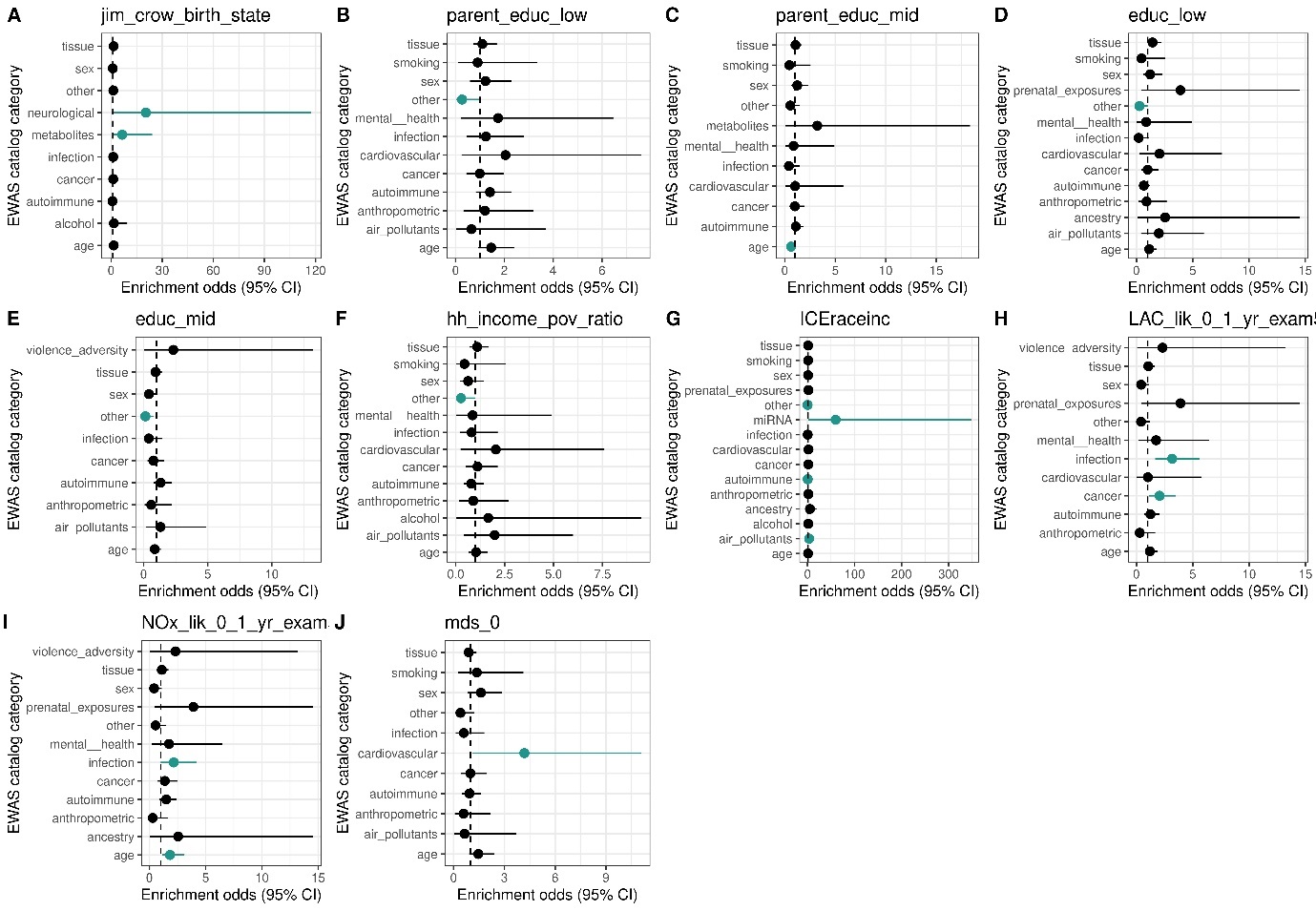

Figure 15: EWAS catalog enrichment plot: MESA white NH JHU + COL subgroup

----------------------------------------------------------------------

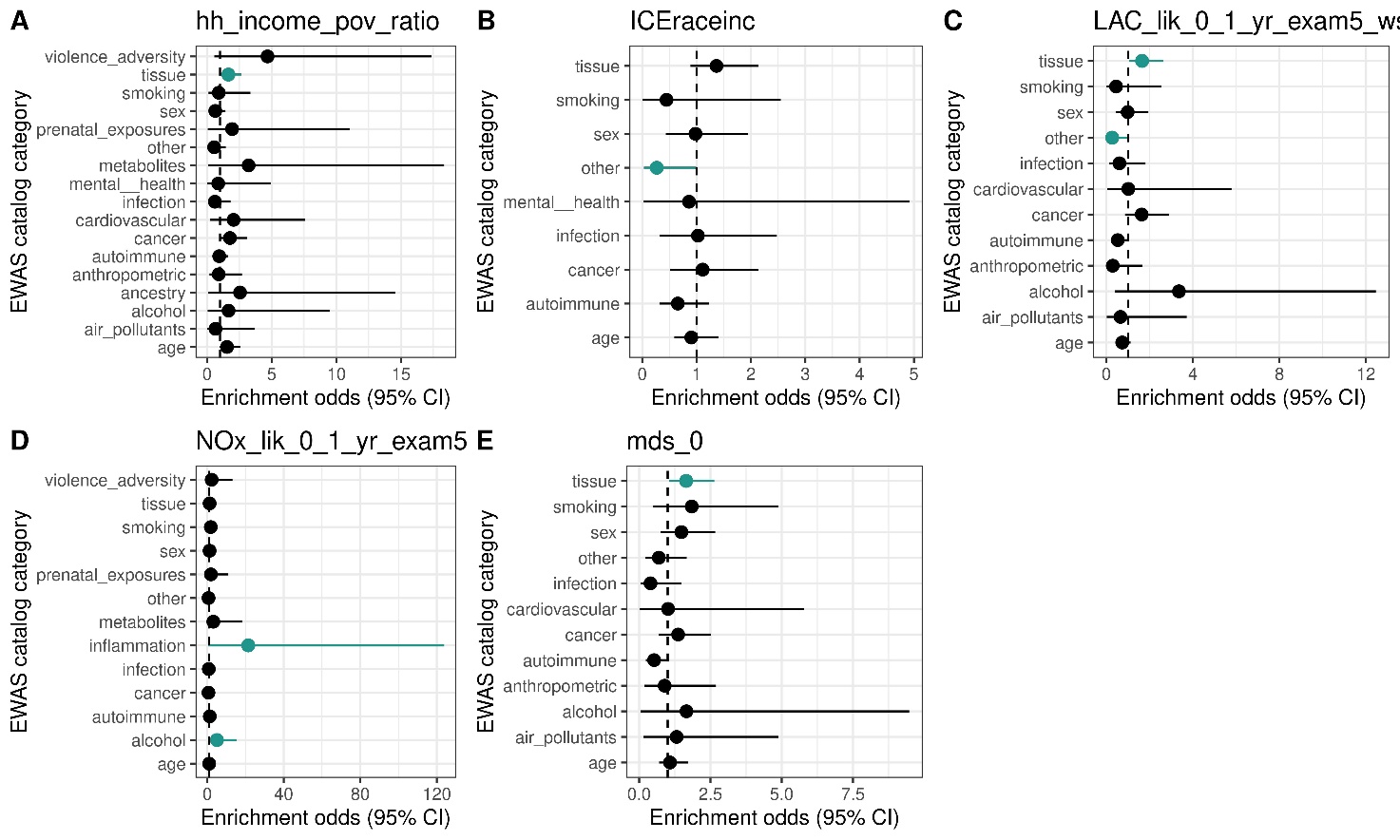

Figure 16: EWAS catalog enrichment plot: MESA Hispanic JHU + COL subgroup

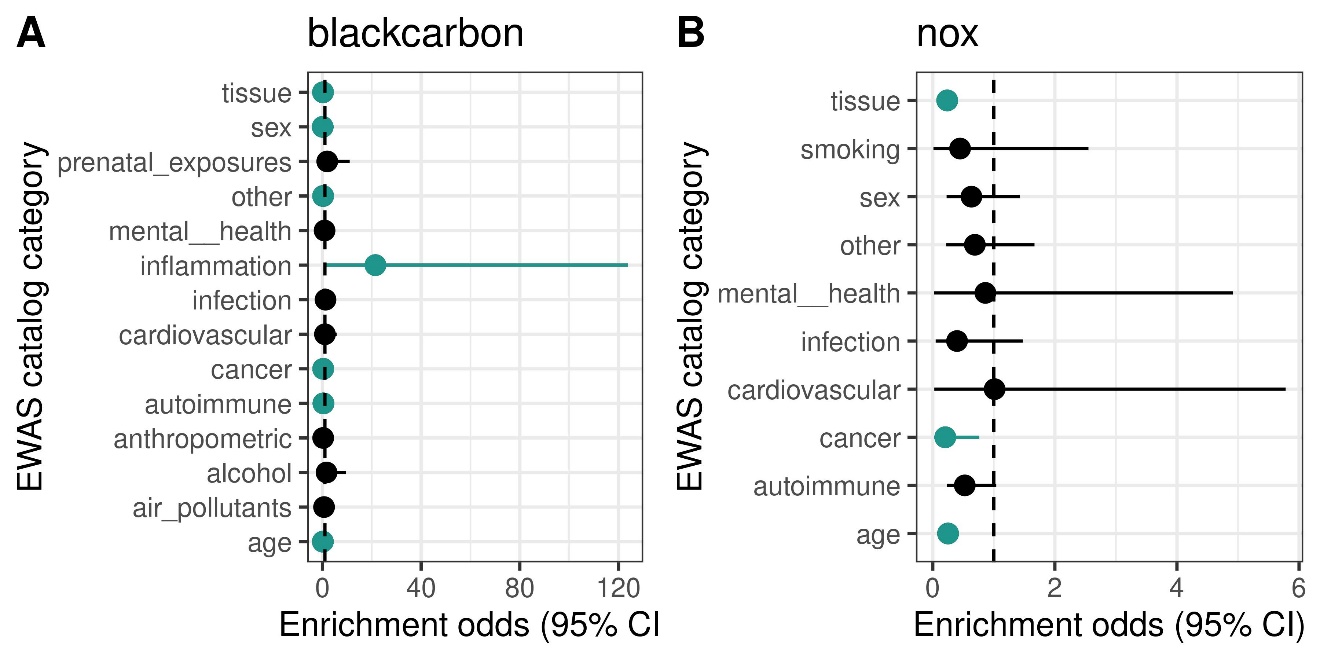

Figure 17: EWAS catalog enrichment plot: MBMS meta-analysis

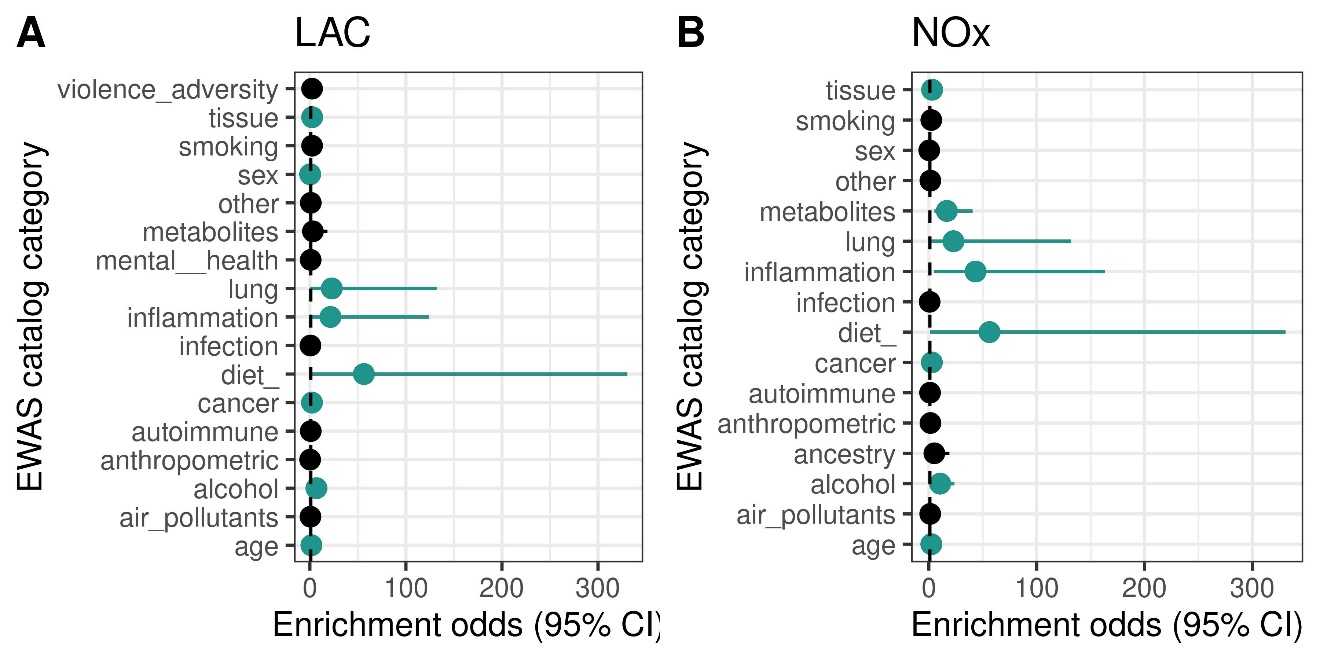

Figure 18: EWAS catalog enrichment plot: MESA meta-analysis

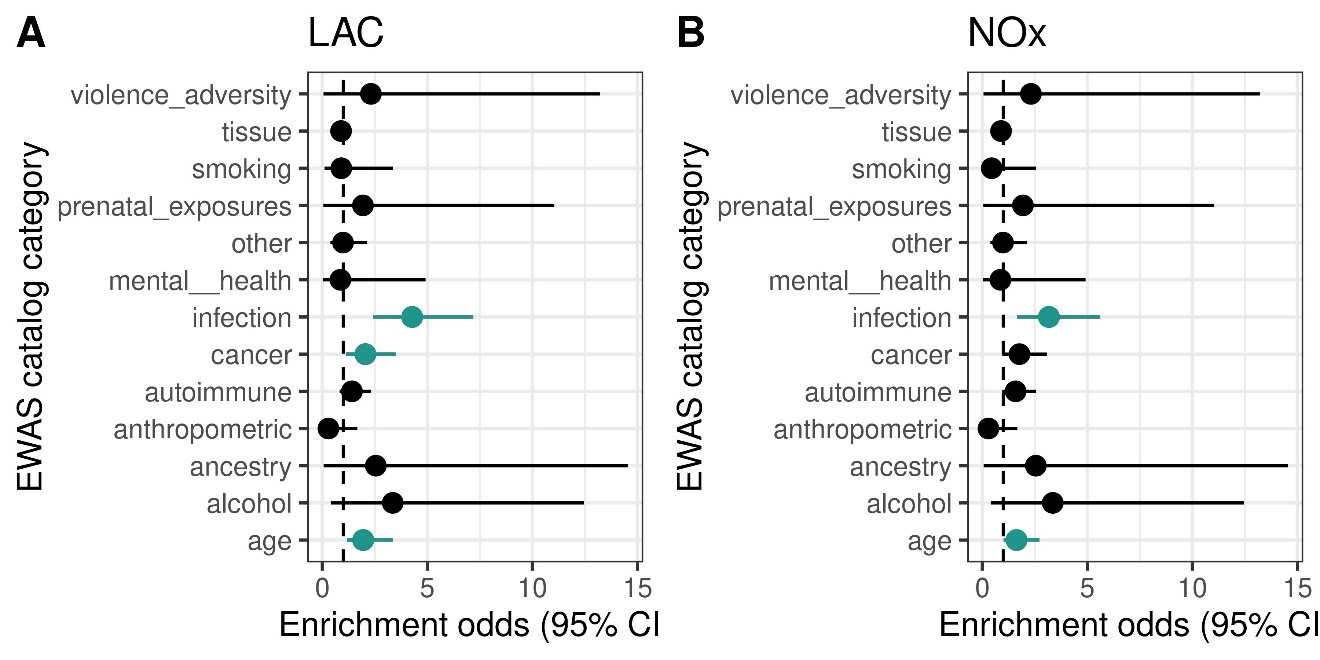

Figure 19: EWAS catalog enrichment plot: MESA meta-analysis (JHU and COL subgroup)

### Enrichment for genomic features

#### MBMS full cohort

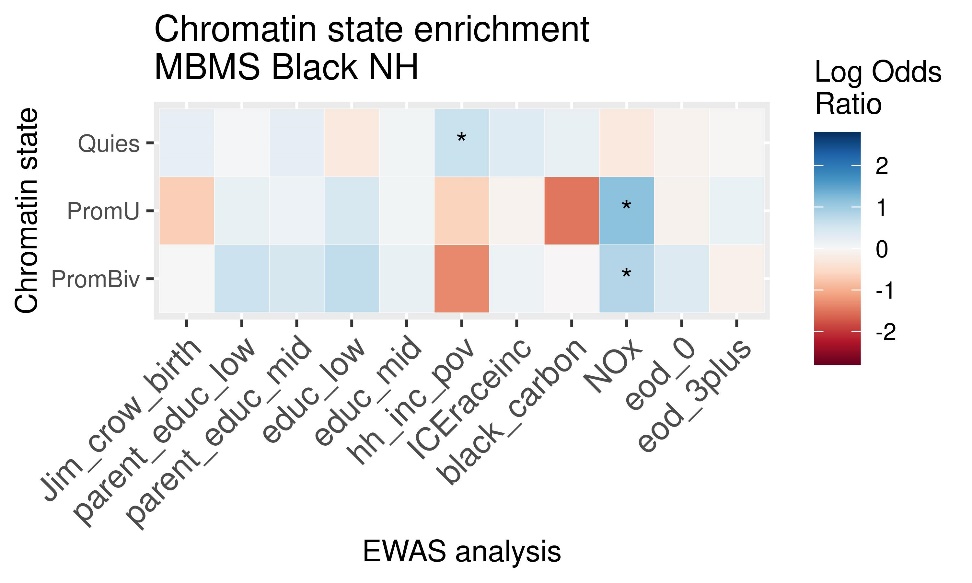

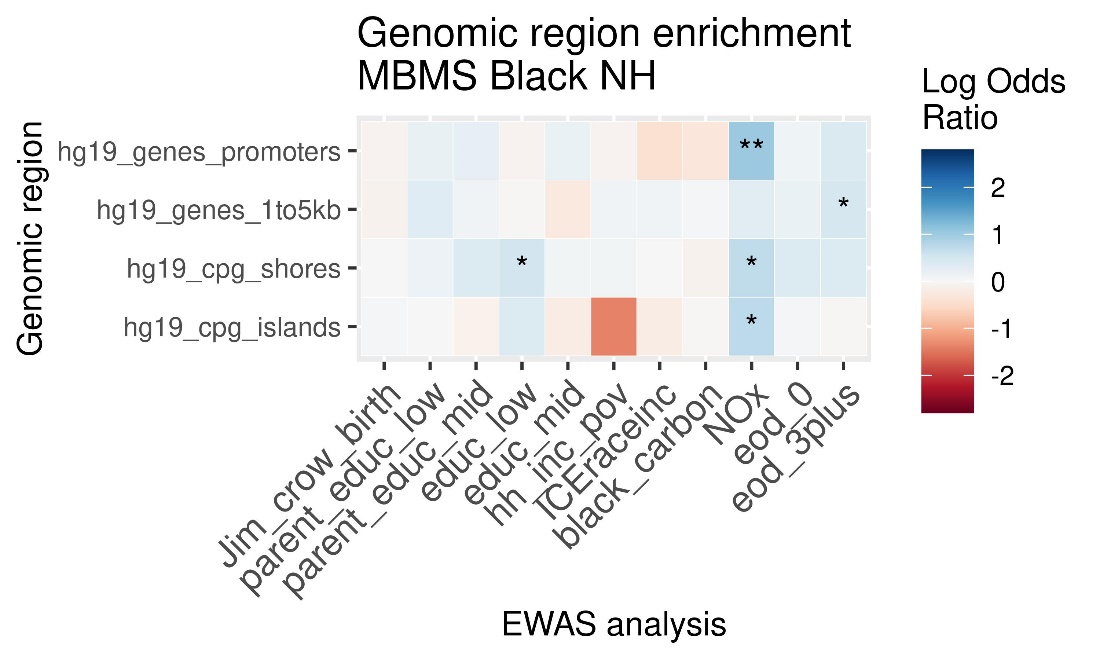

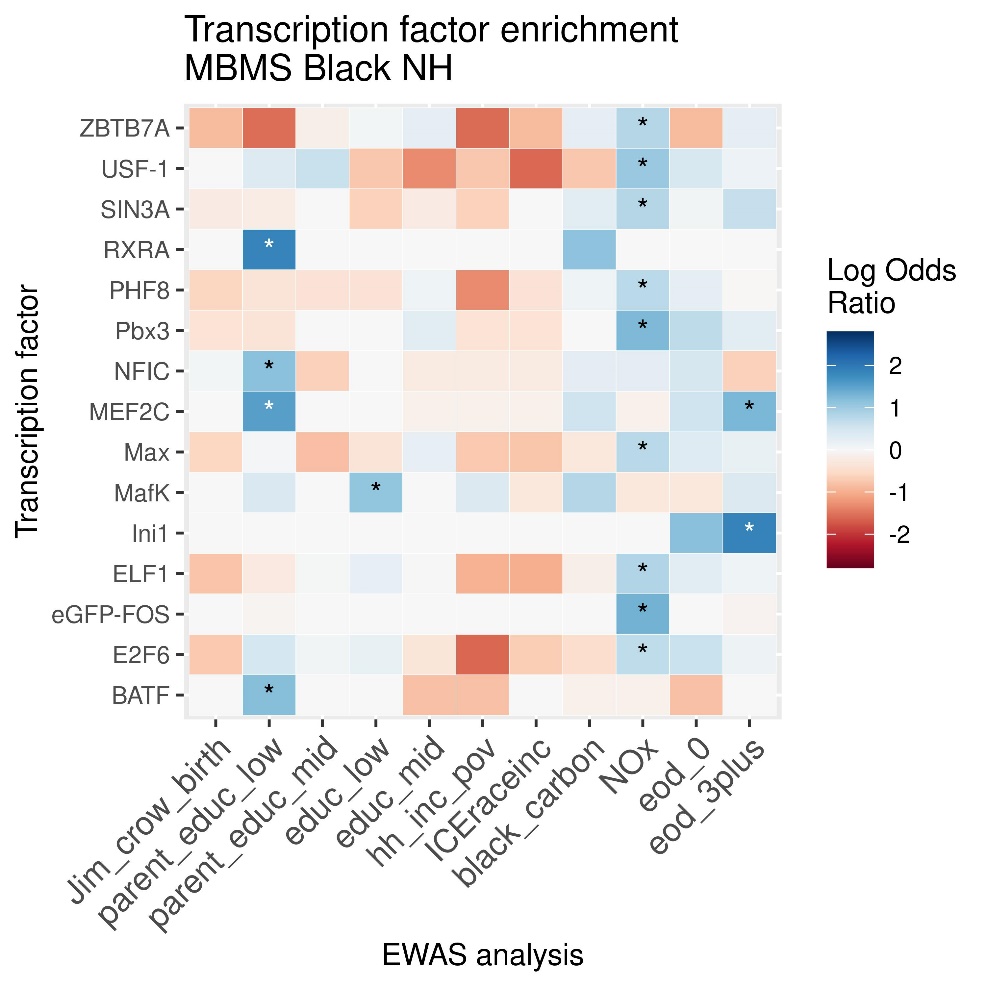

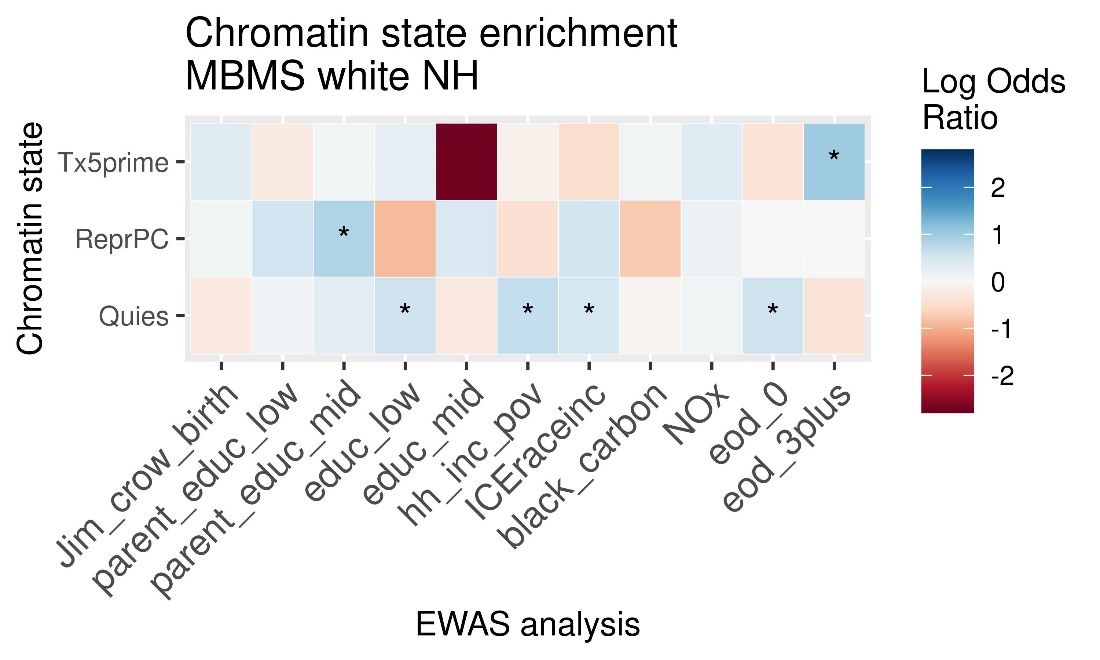

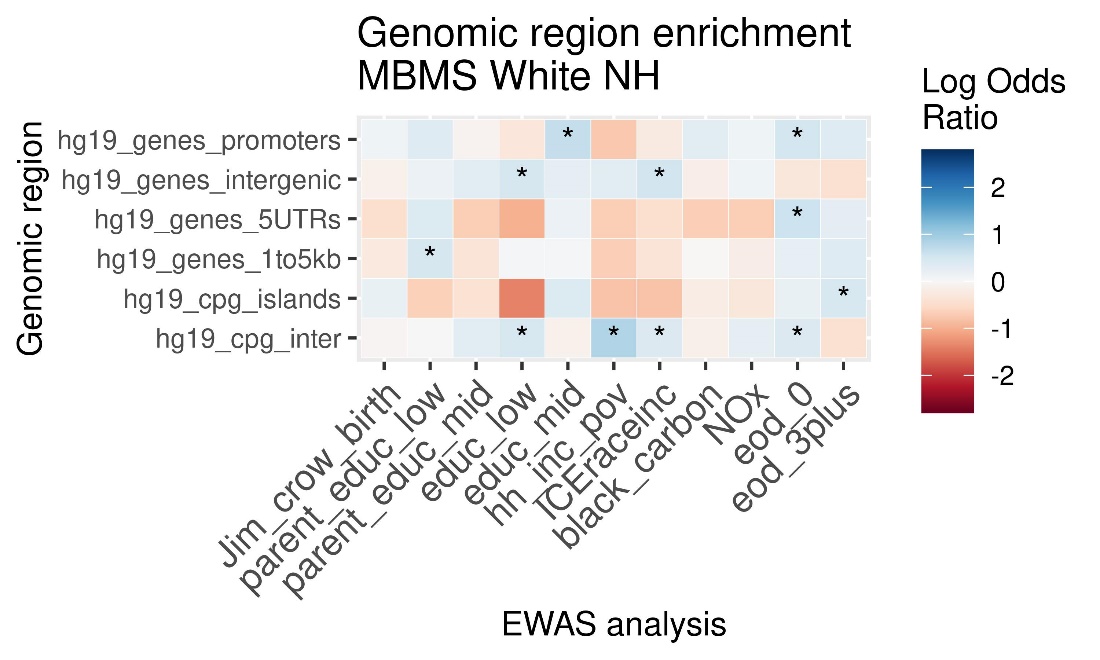

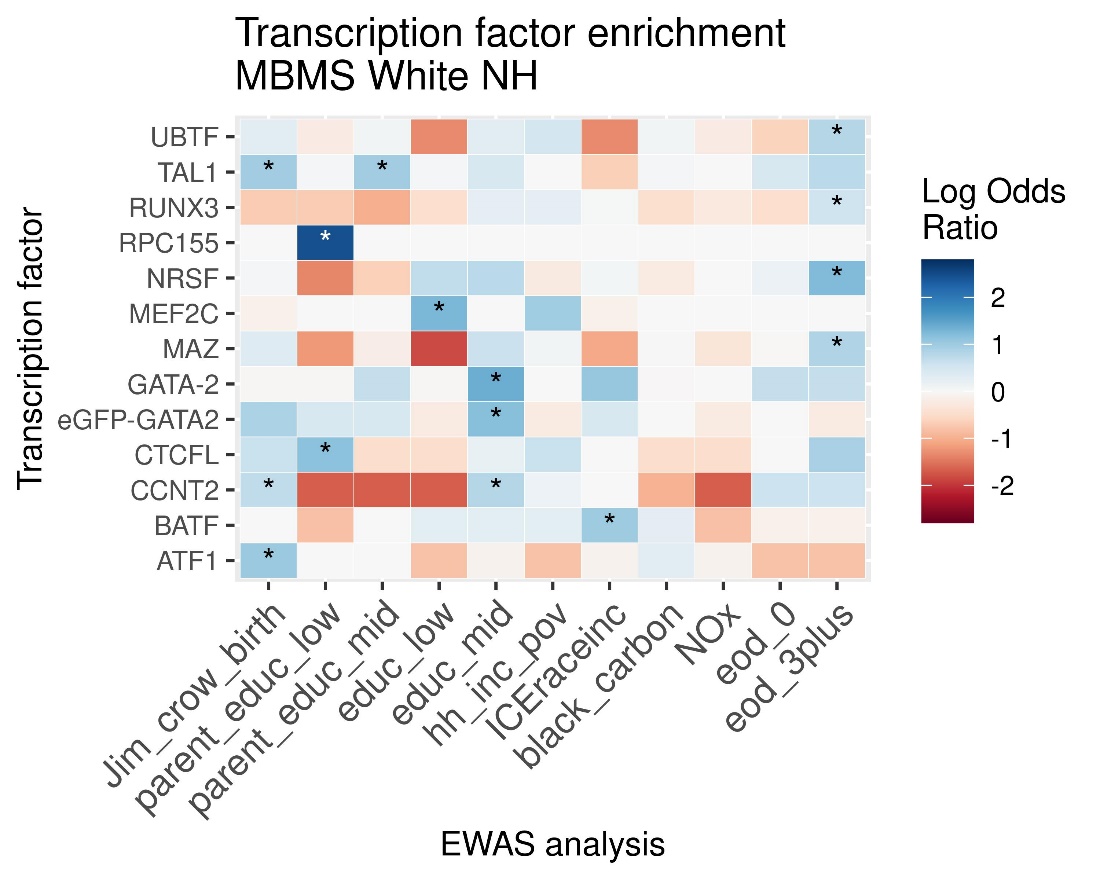

#### MESA full cohort

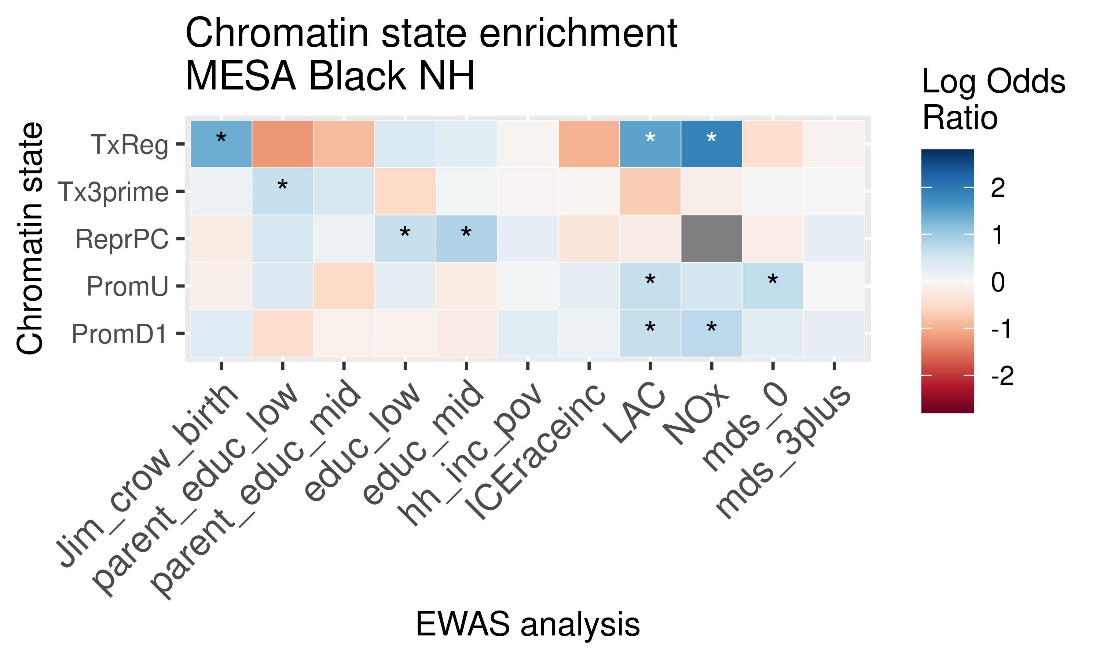

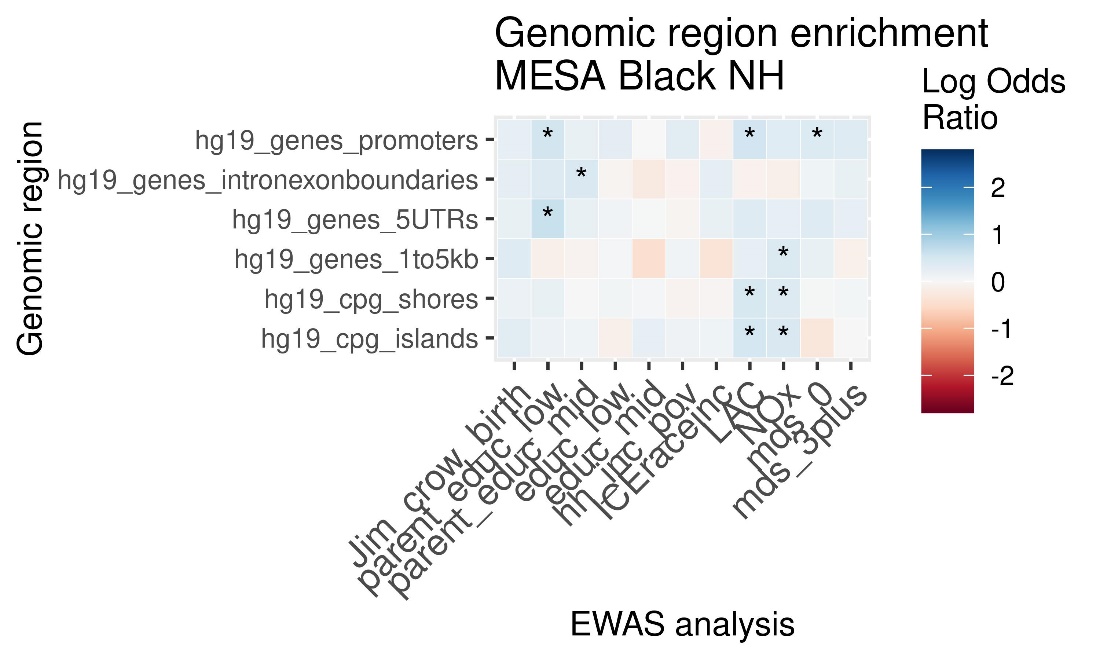

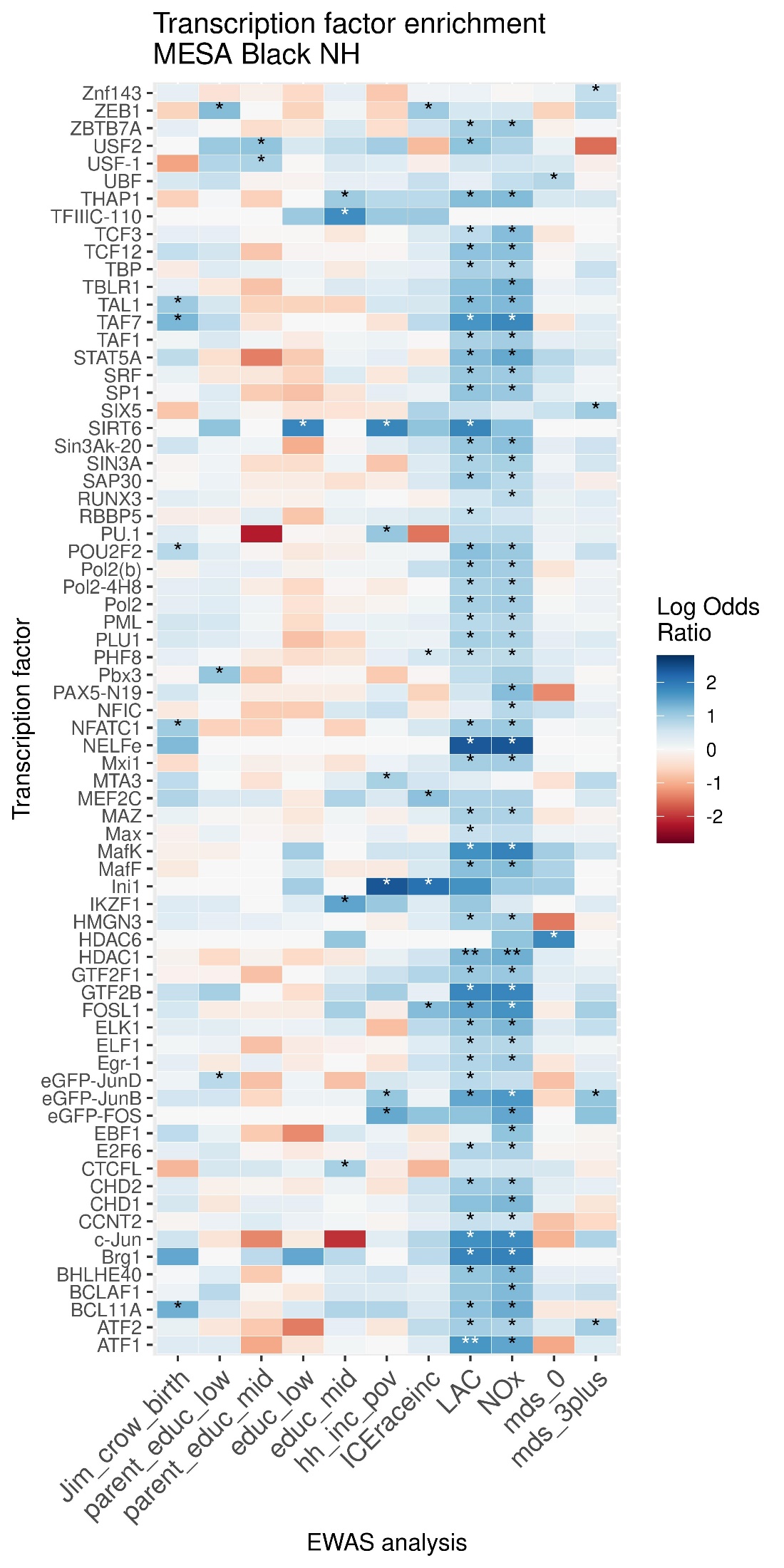

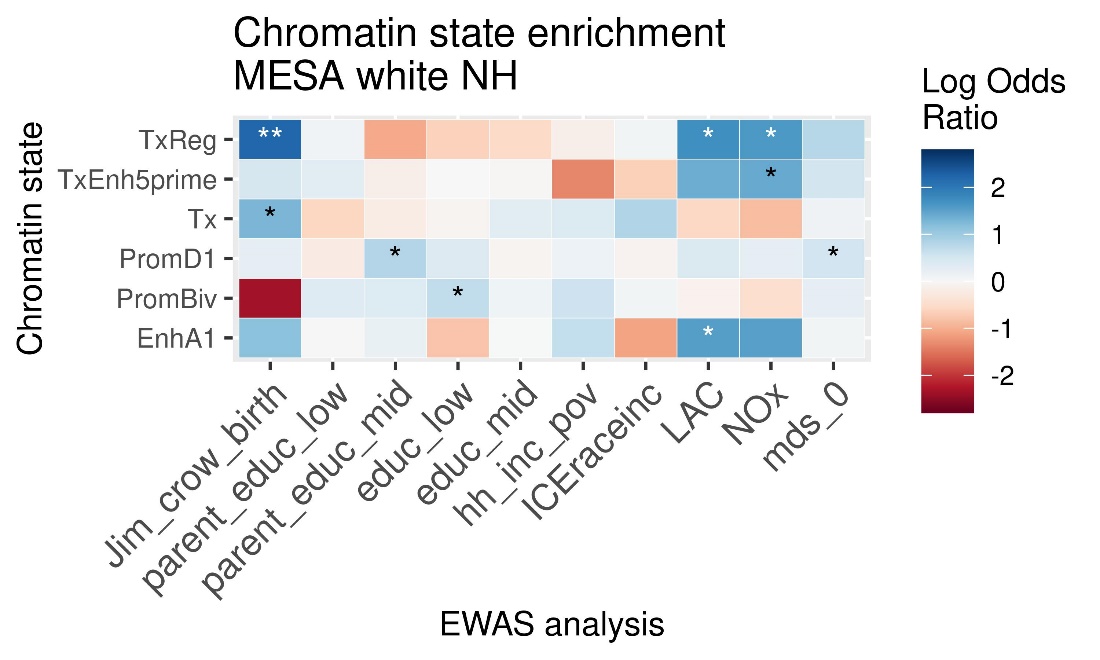

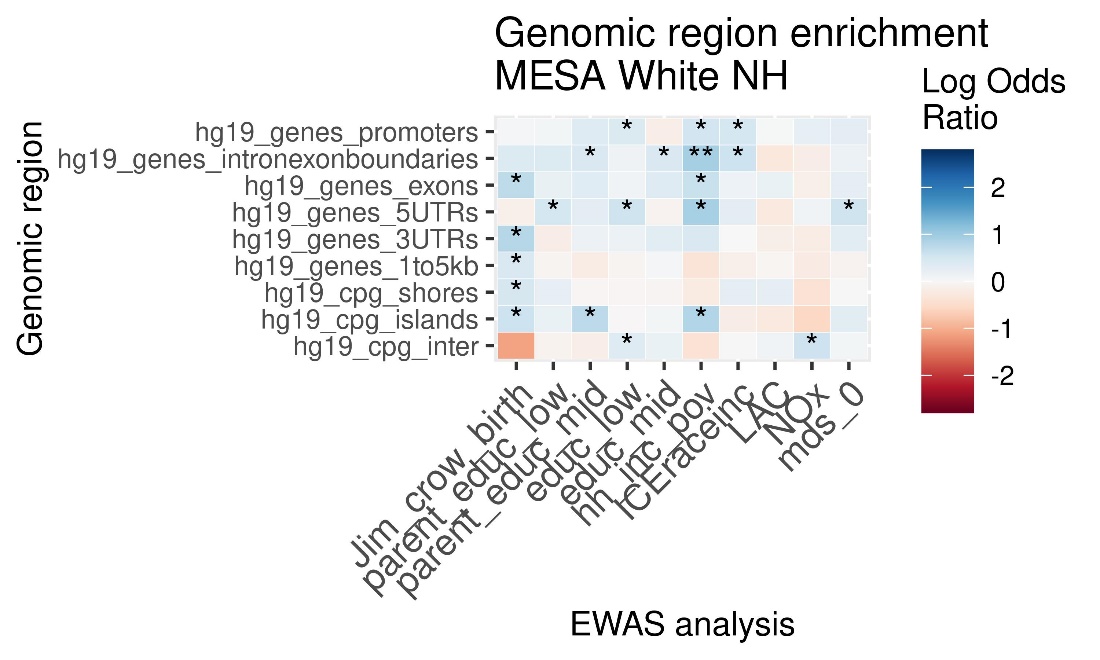

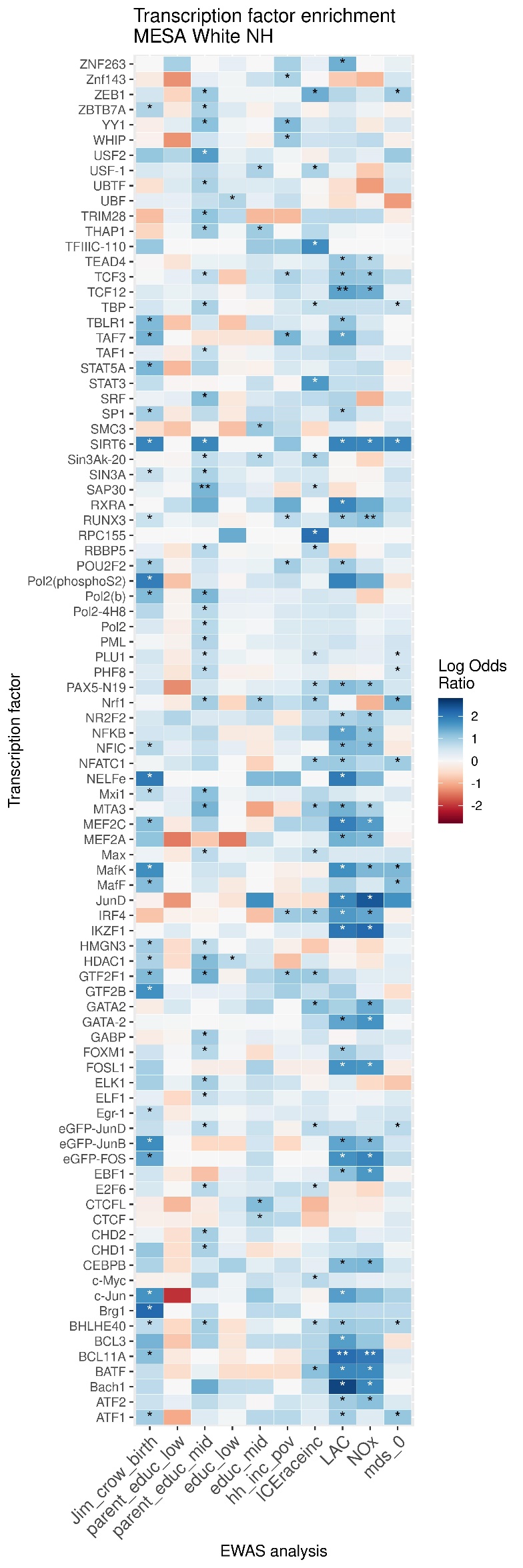

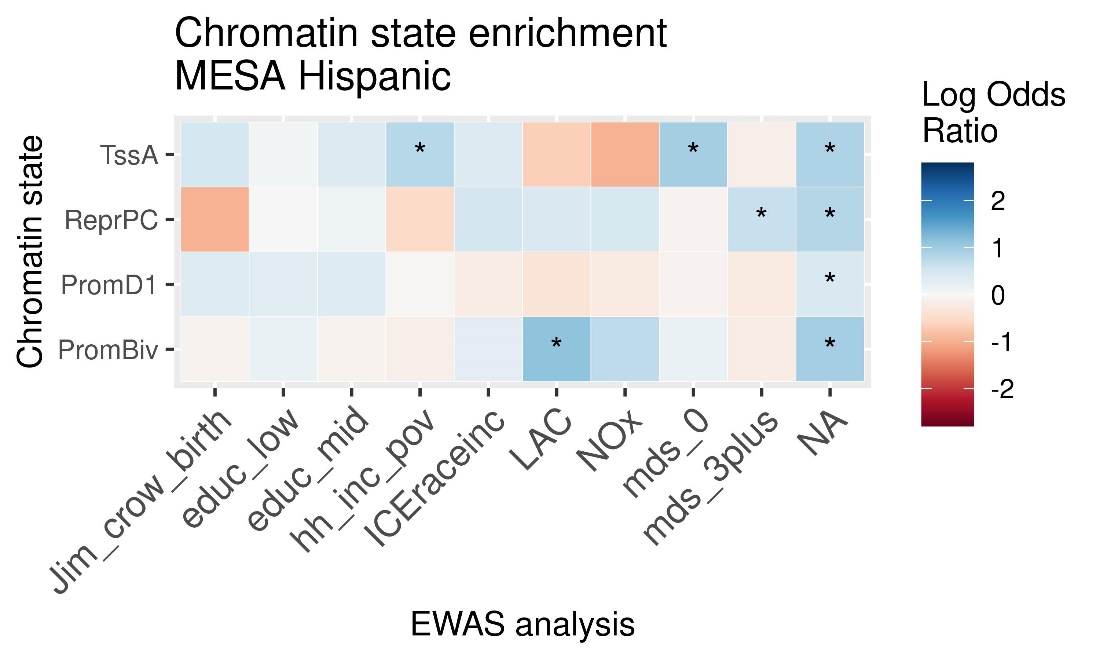

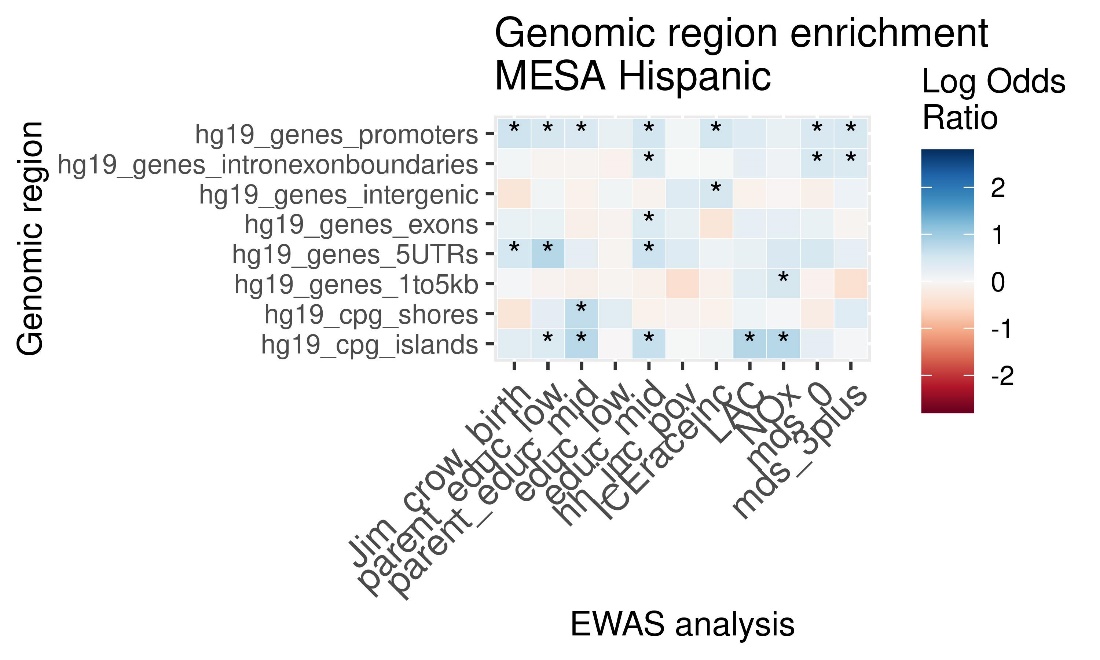

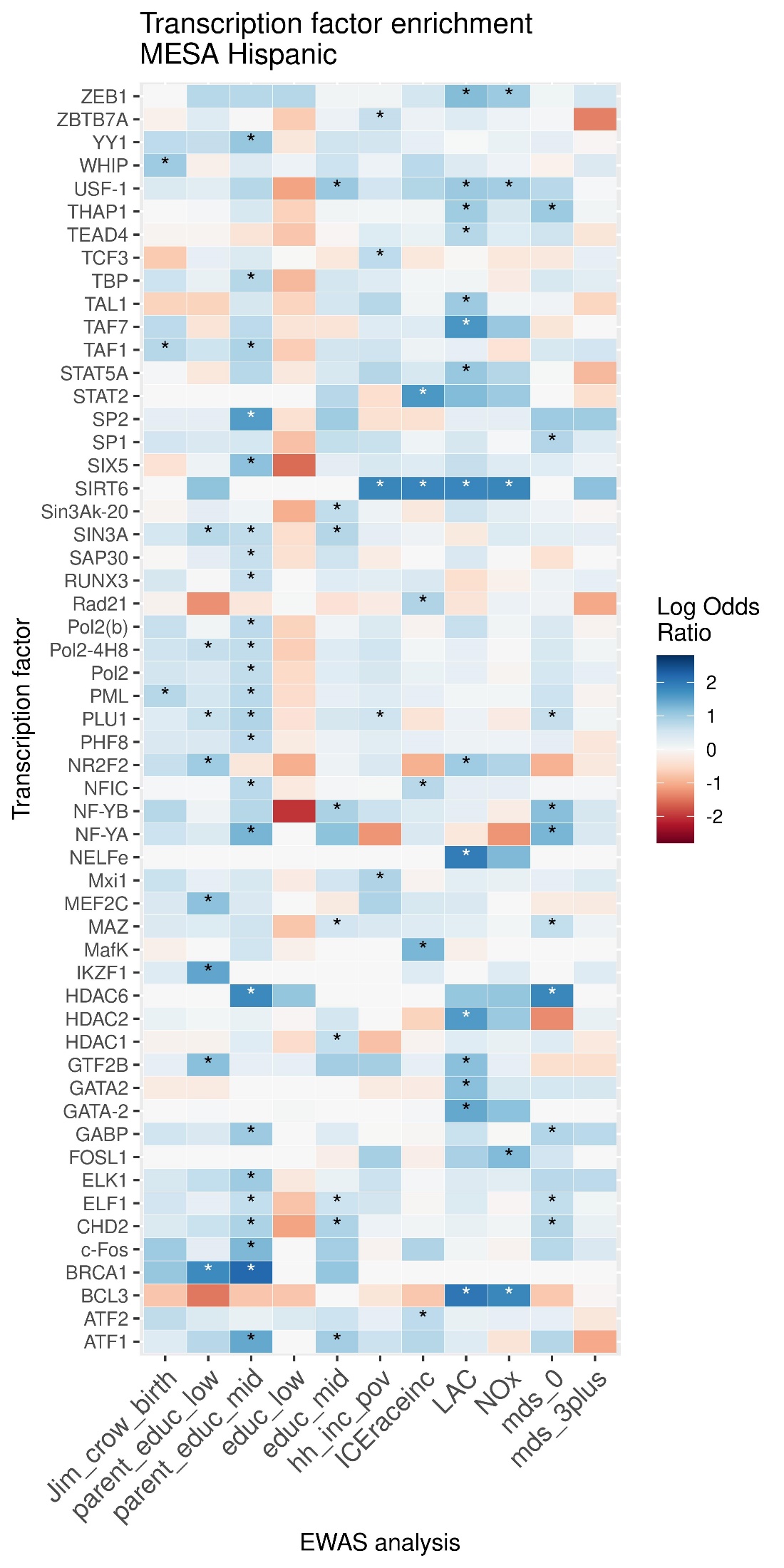

### MESA subgroup

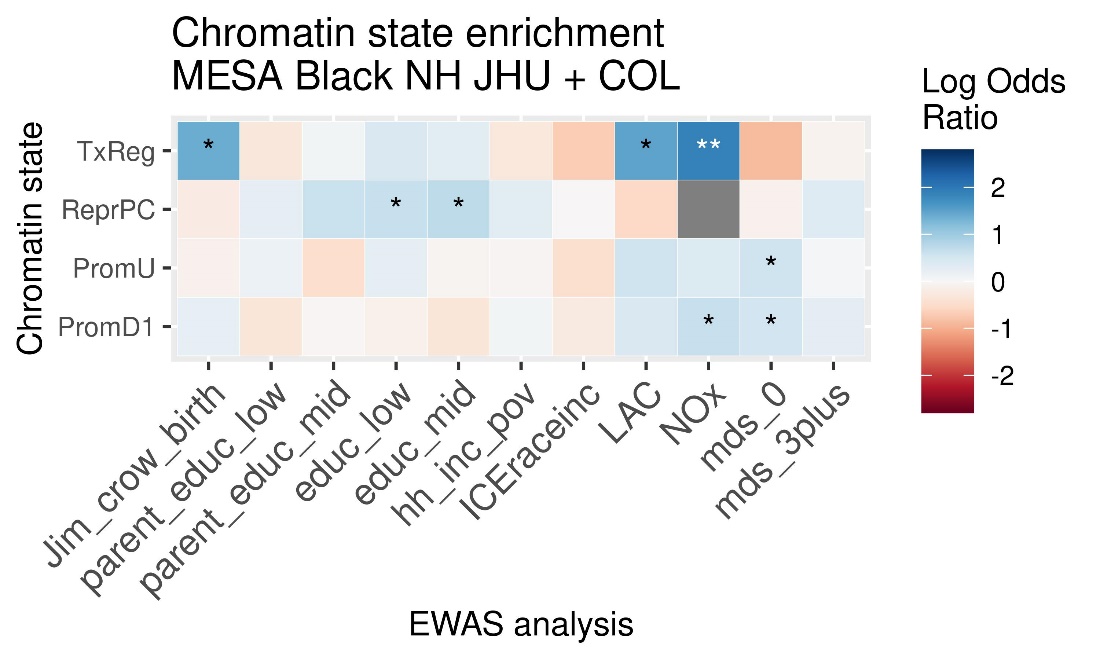

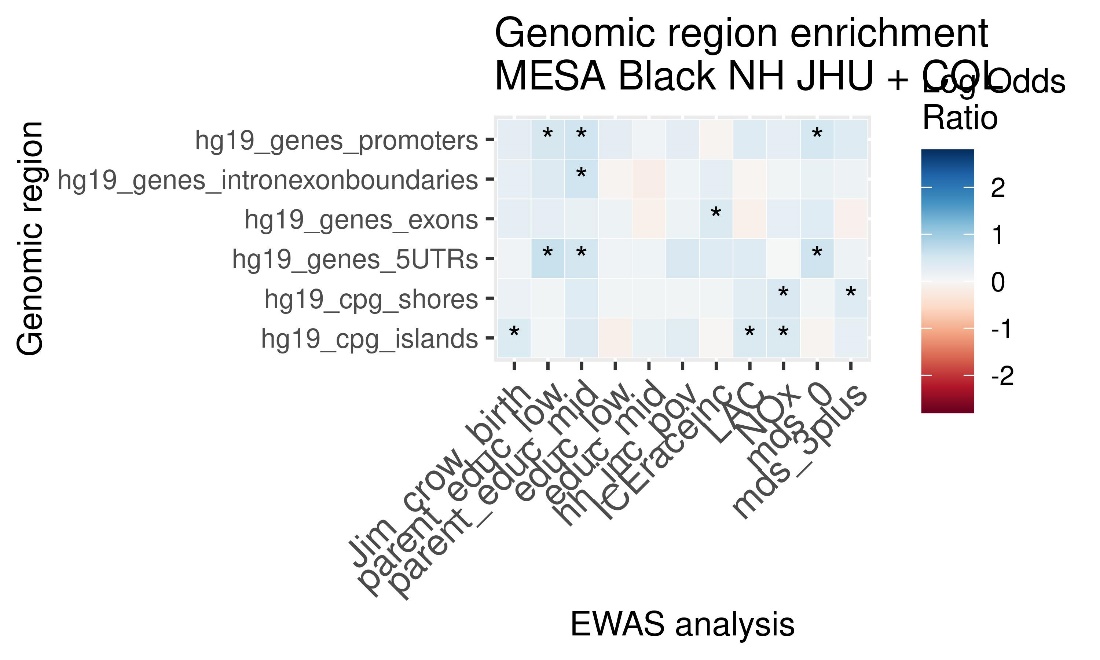

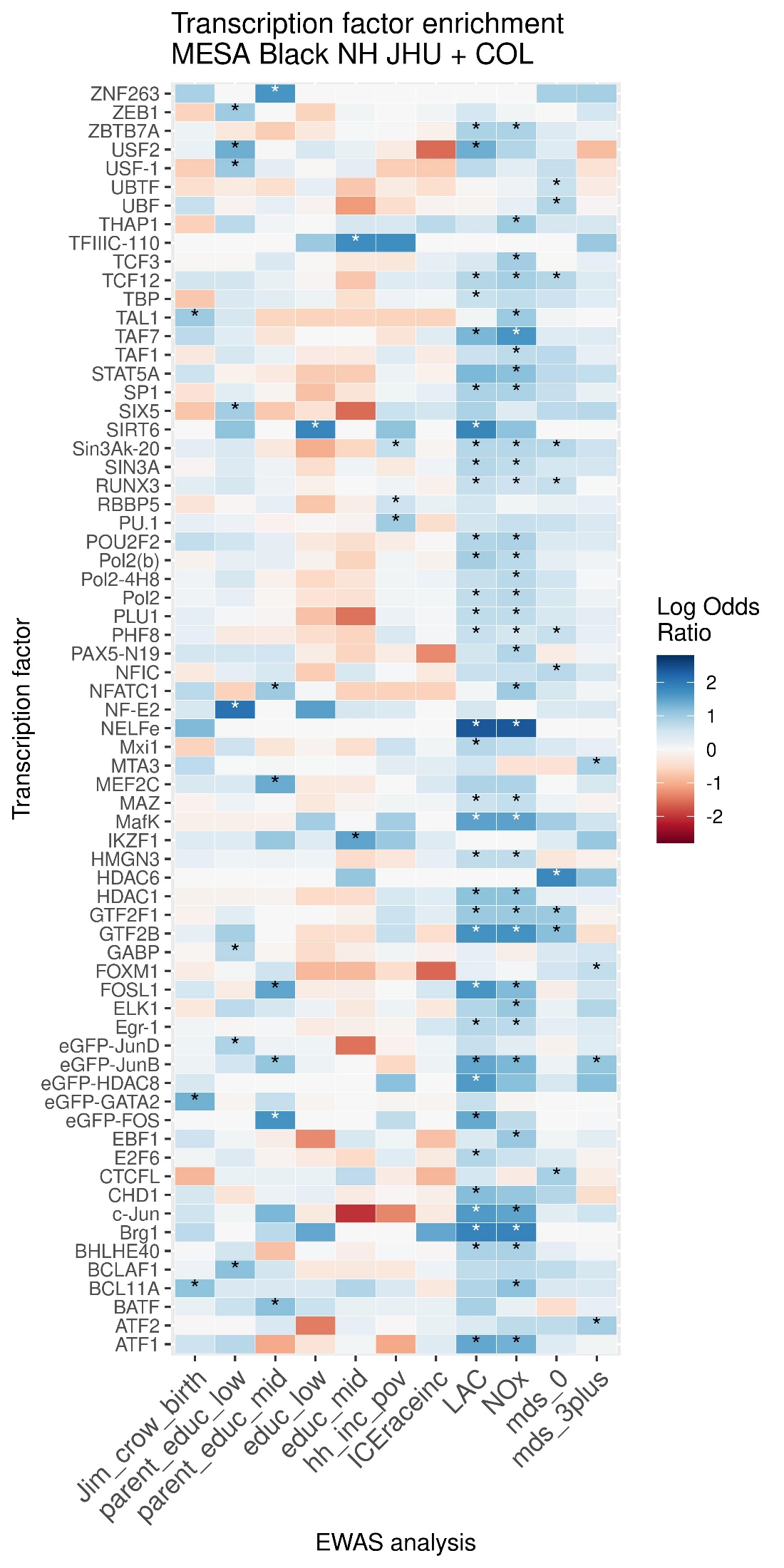

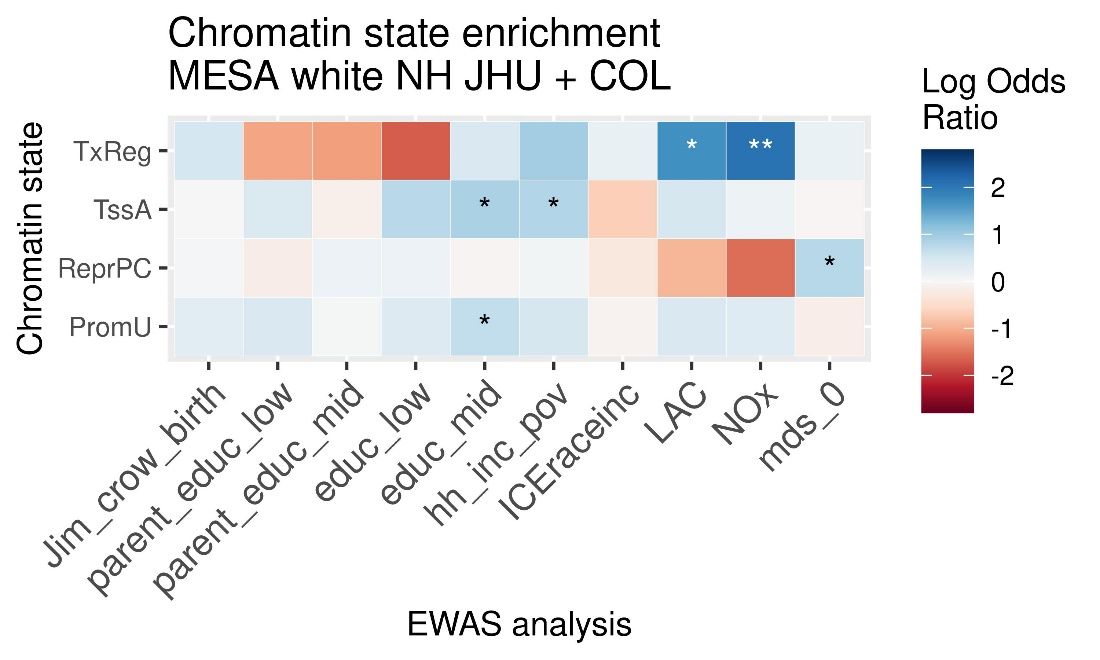

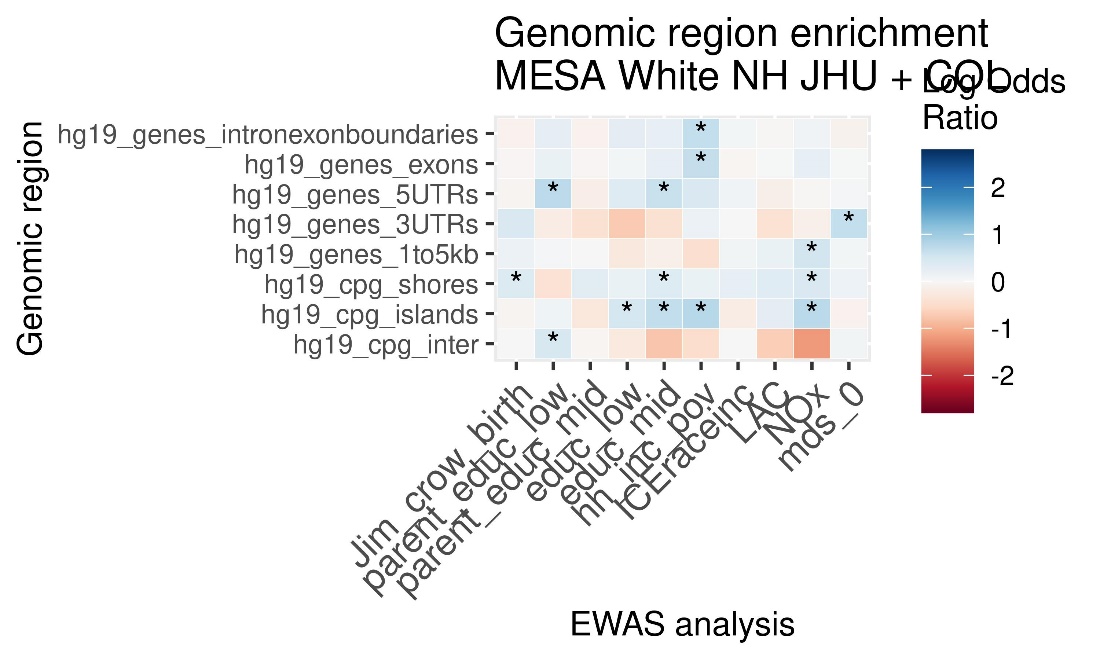

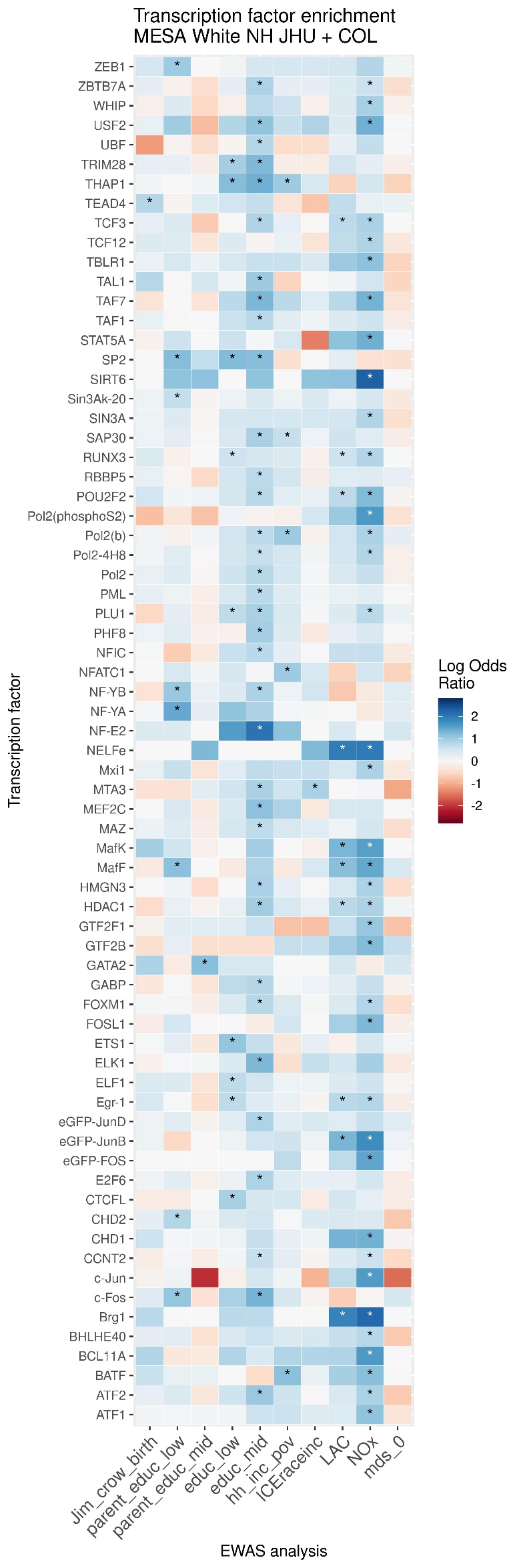

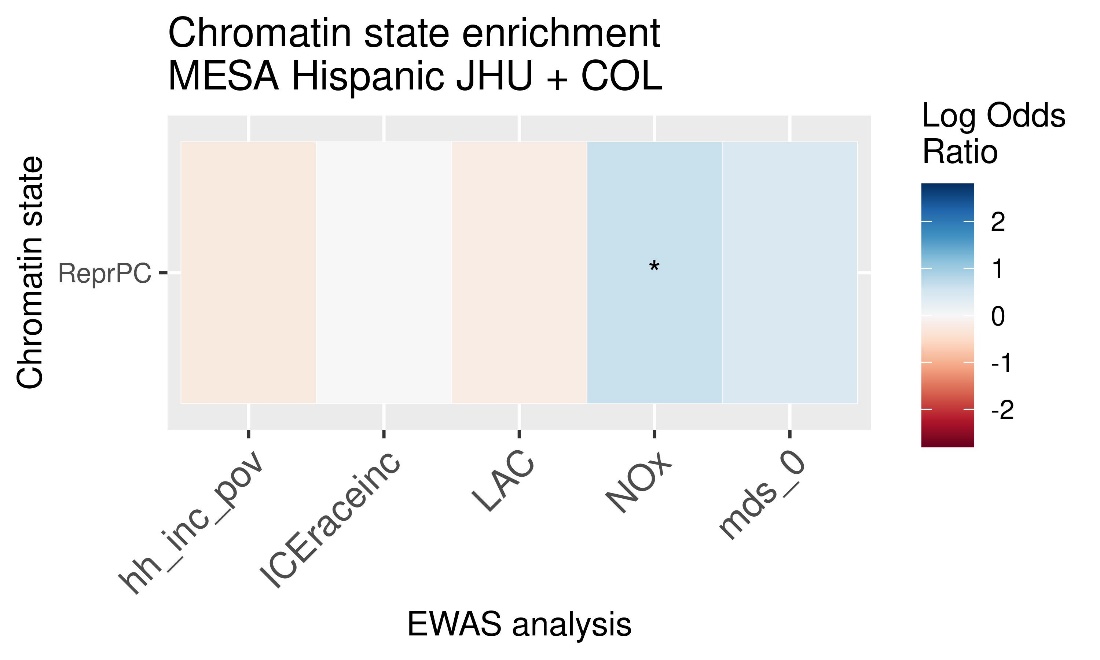

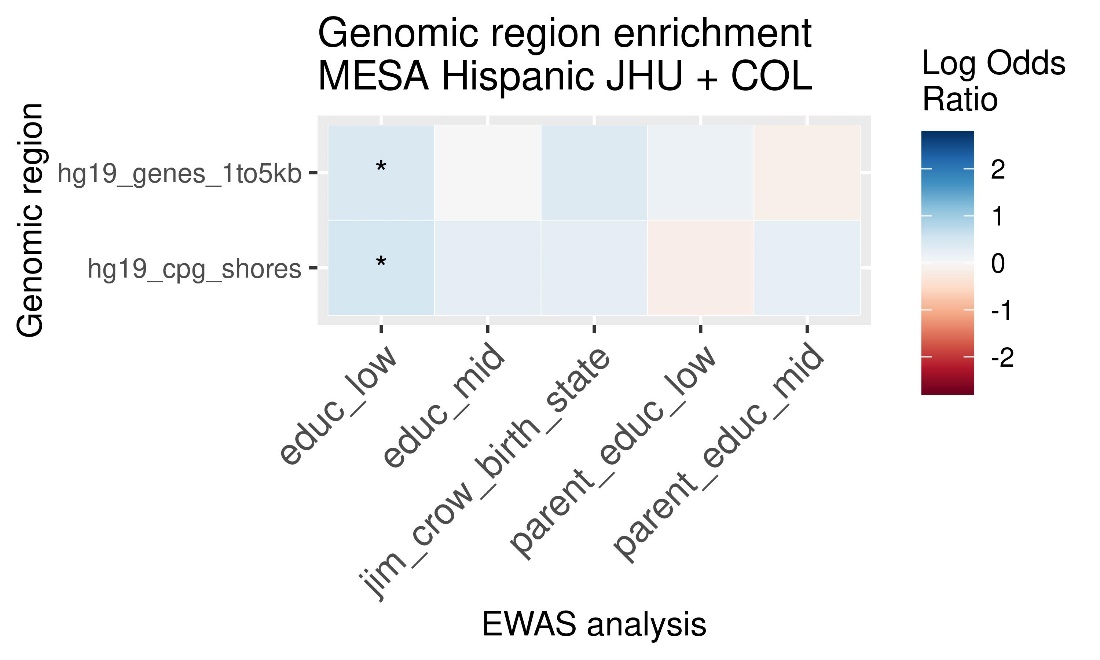

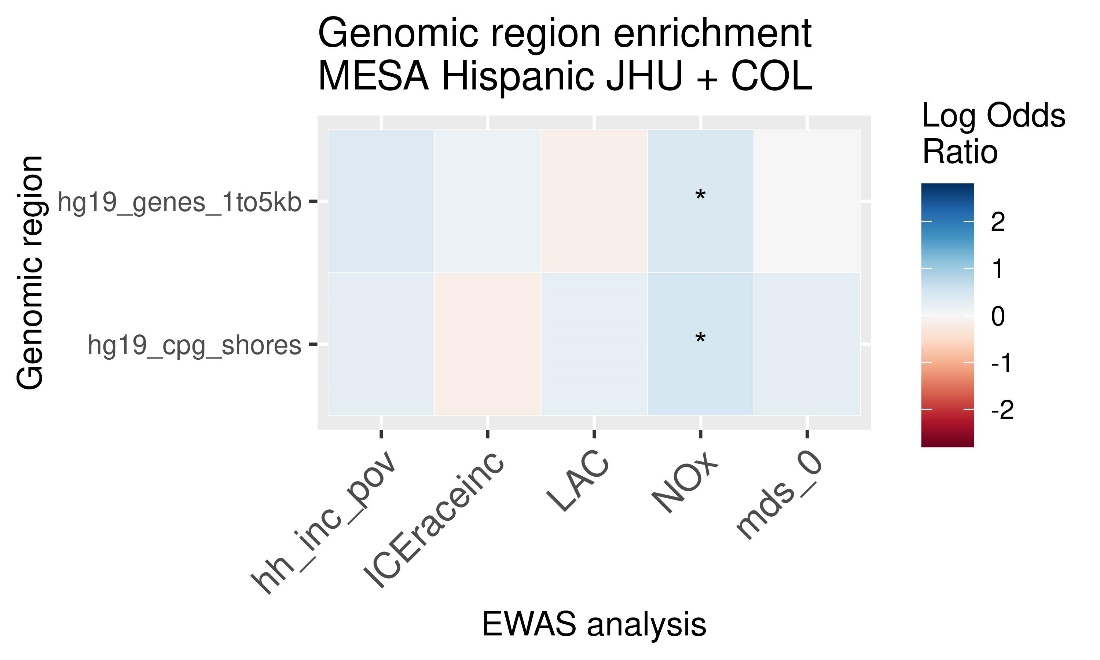

### MBMS meta-analysis

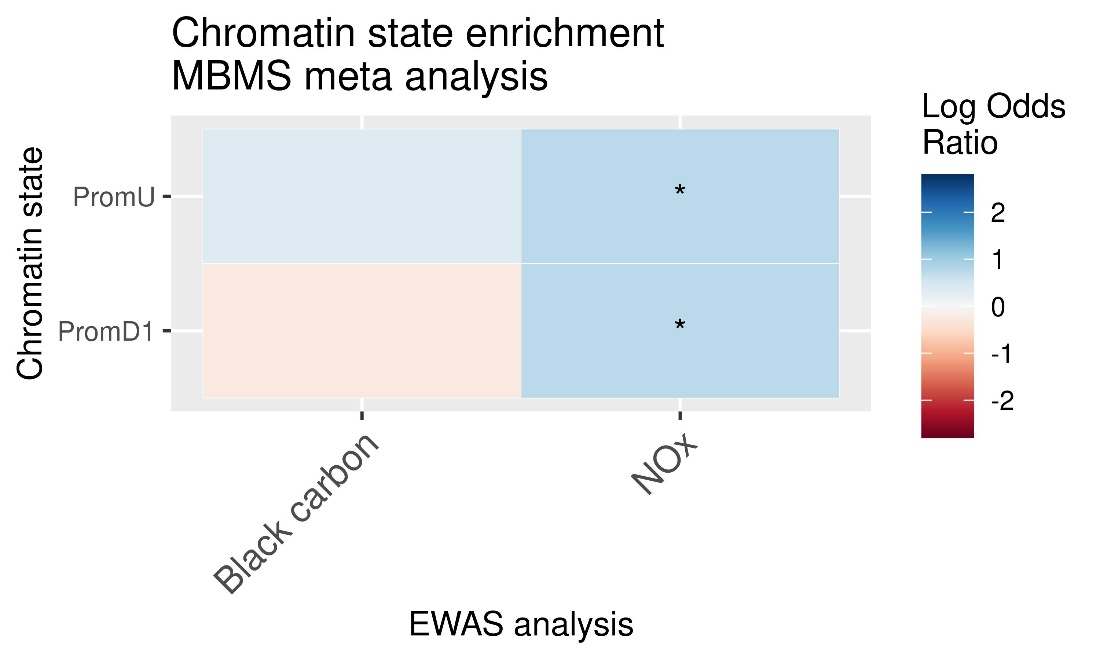

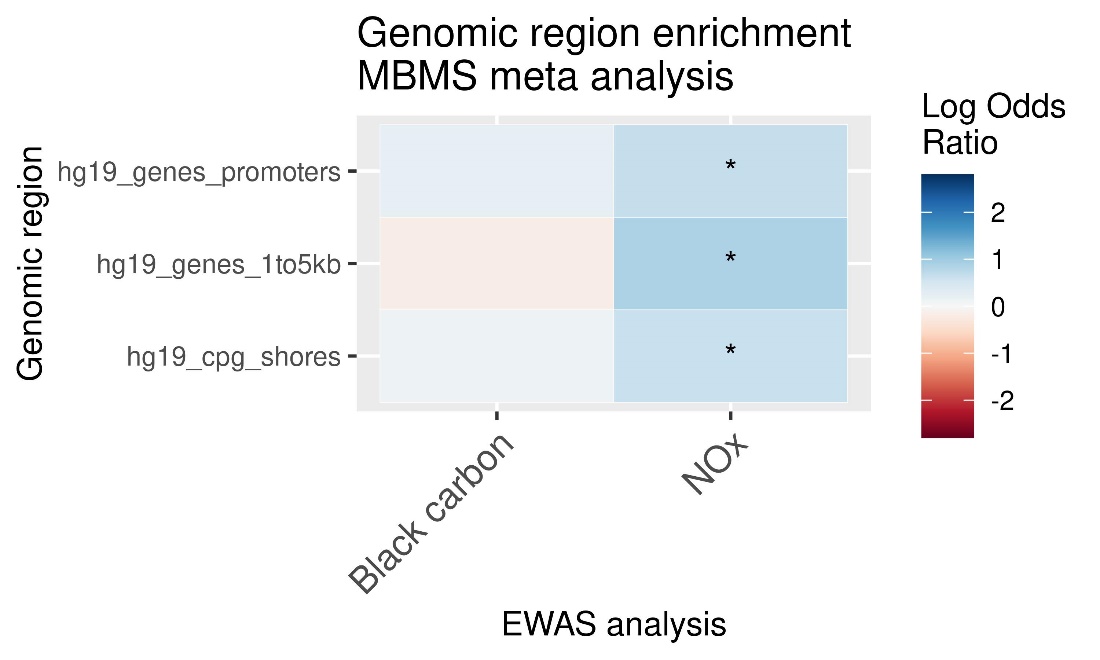

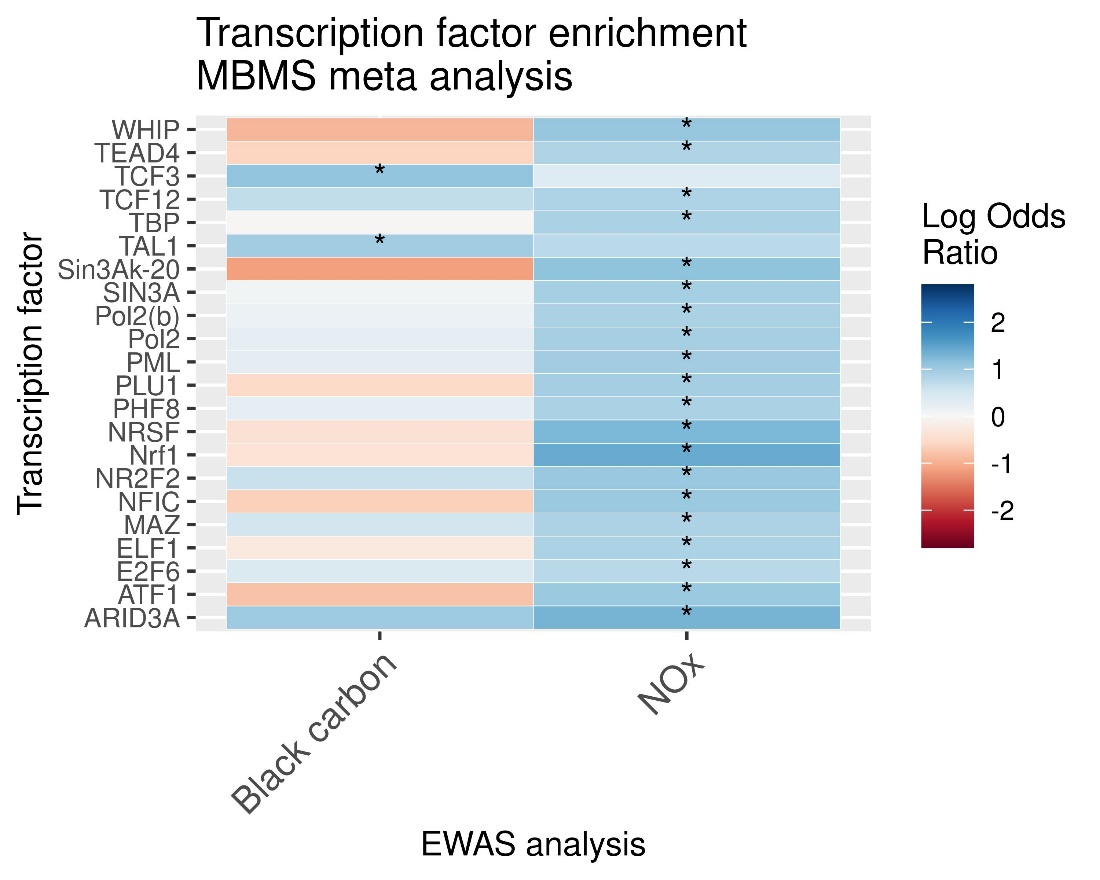

### Mesa meta-analysis (full cohort)

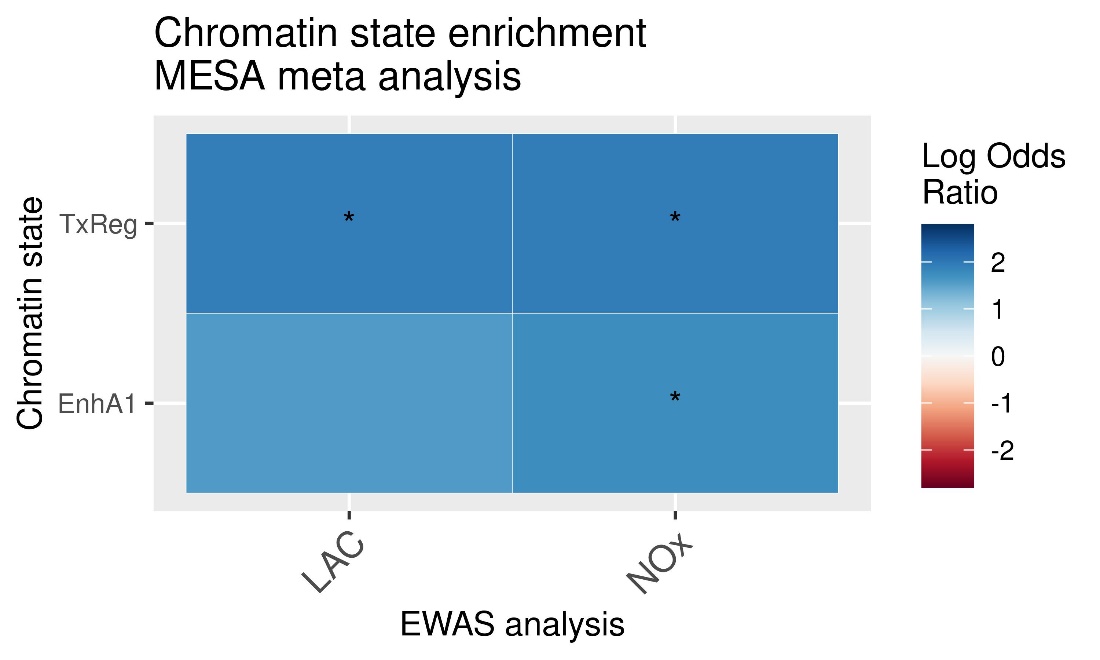

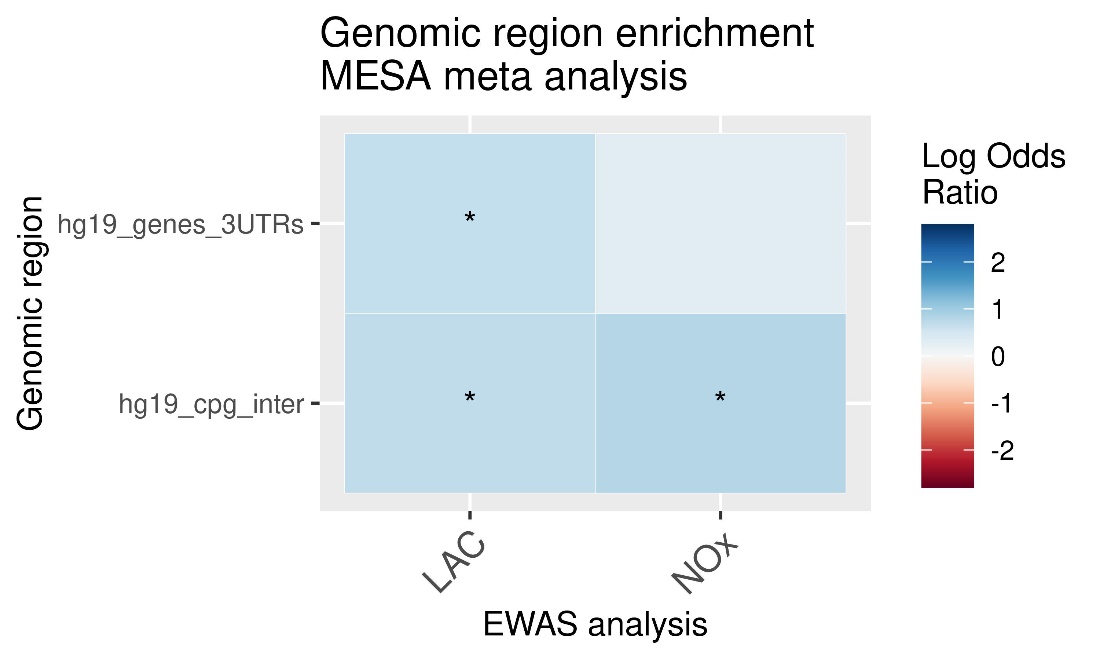

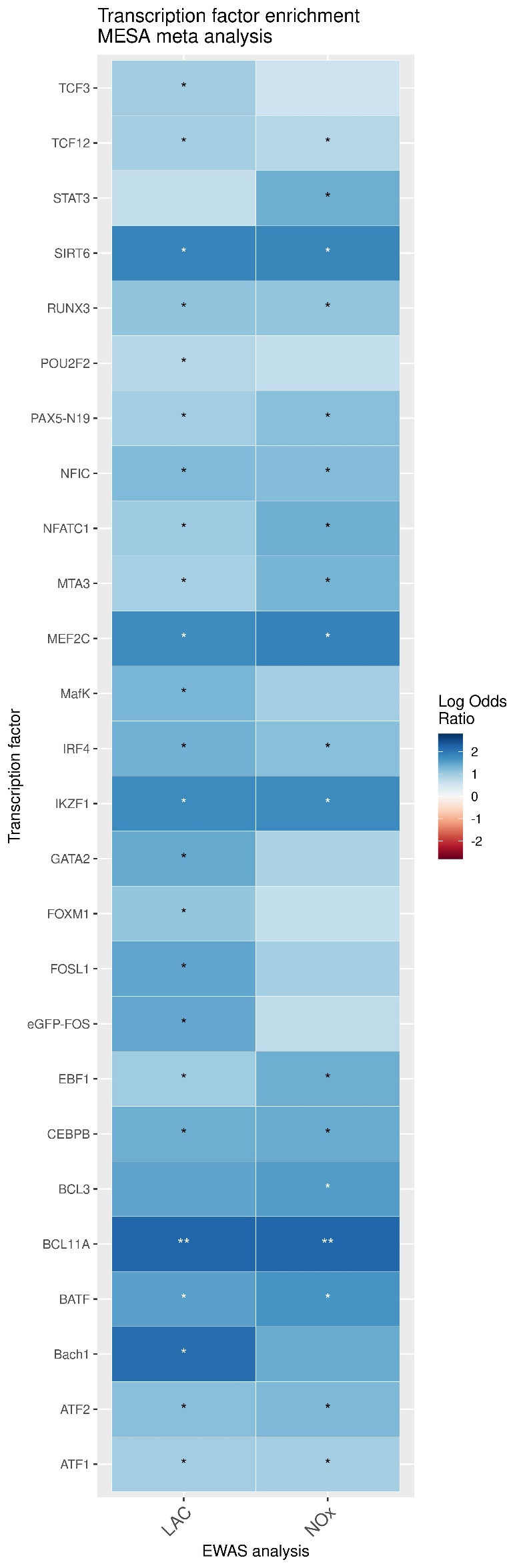

### MESA meta-analysis (subgroup)

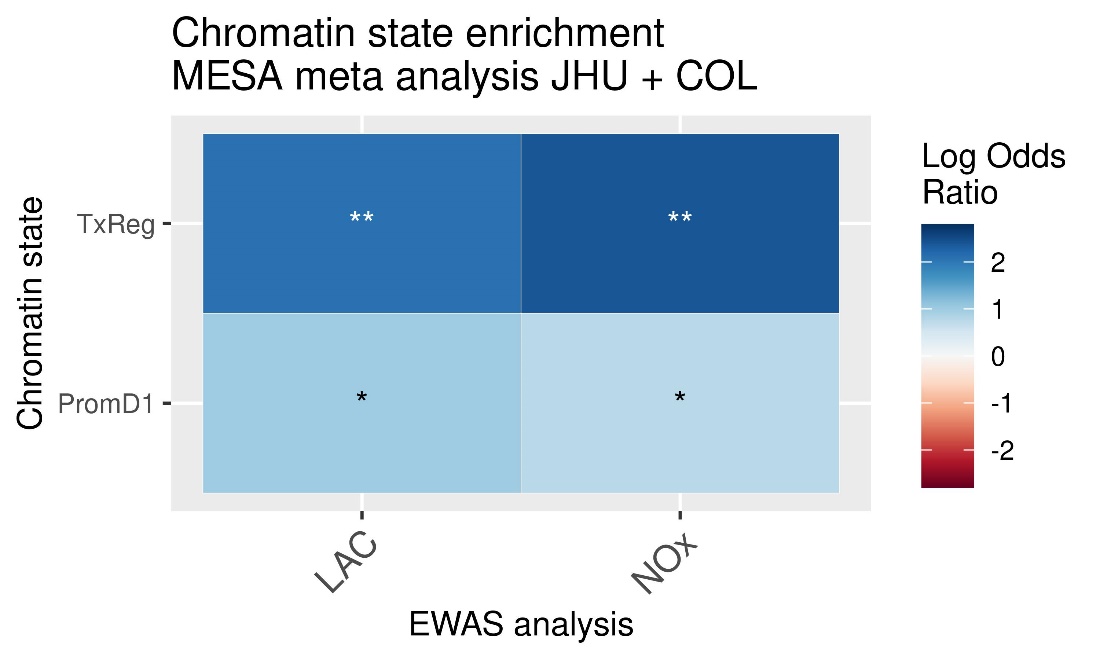

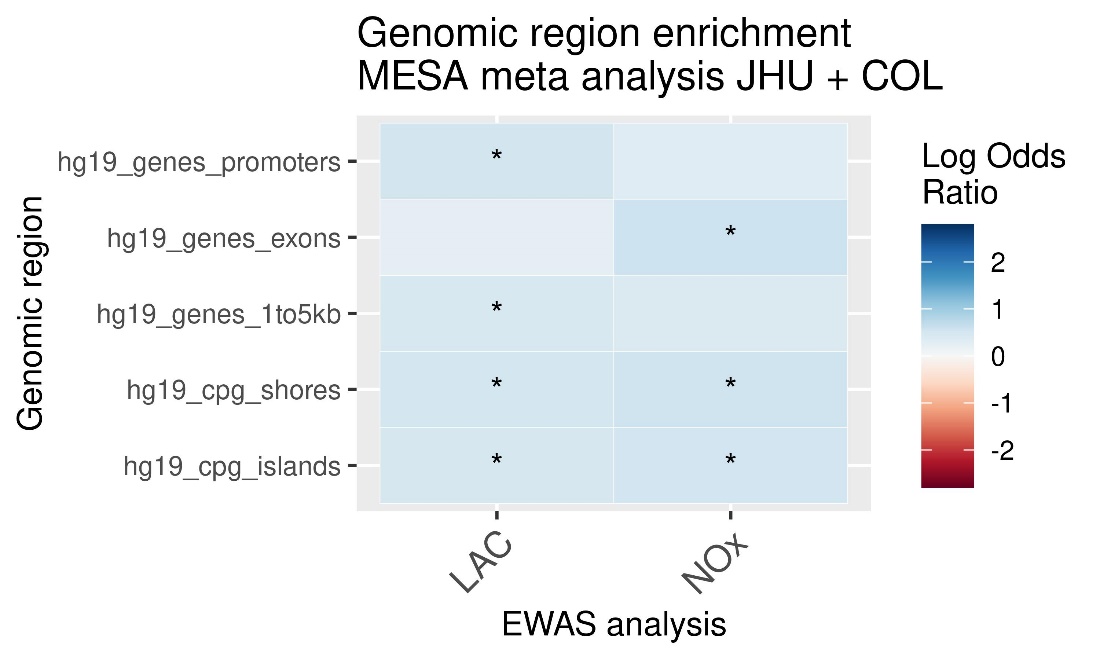

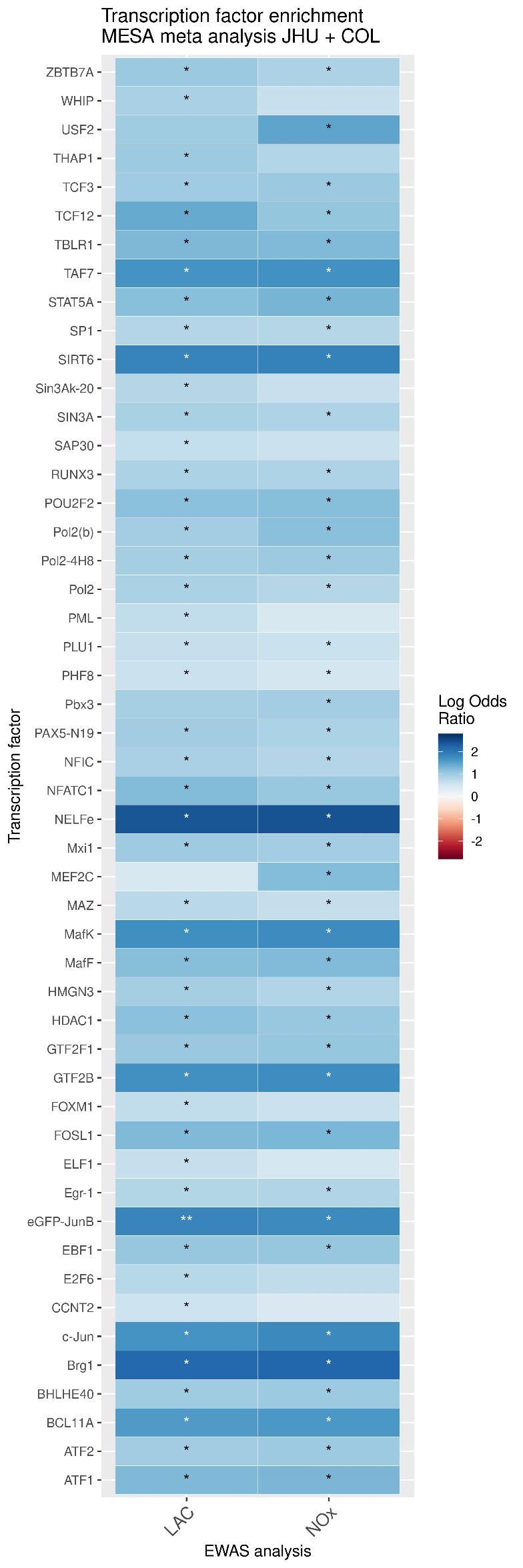
