## Supplementary Materials for "An epigenome-wide analysis of DNA methylation, racialized and economic inequities, and air pollution"

### Methods

#### Participants

MBMS includes participants recruited from four Community Health Centers (CHCs) in Boston, MA between 2008 and 2010, and was designed to investigate the association between racial discrimination and risk of cardiovascular disease, taking into account a range of social and environmental factors. The MBMS cohort and recruitment procedures have previously been described in detail (Krieger, Waterman, et al., 2011); briefly, the study recruited 1005 individuals who met study inclusion criteria and were randomly selected from the patient rosters of the CHCs. Participants were eligible if they were aged between 35 and 64 years, had been born in the US, and self-identified as white non-Hispanic or Black non-Hispanic.

Among the 1005 MBMS participants, 85% provided a finger prick blood sample on to filter paper (409 Black; 466 White), and consequently biological material was limited and in some instances of poor quality. Blood spots were stored at -20°C, and DNA was extracted from blood spots using the QIAamp DNA Investigator Kit for FTA and Guthrie cards, with samples randomised across 96 well plates. Of the 875 participants who provided blood spots, 472 of the samples were judged to be suitable for DNA extraction (blood spots judged not to be suitable were primarily collected at the first CHC where recruitment took place, whose membership was predominantly white). Of those, 48 yielded less than 40ng of DNA; we have previously determined this to be an input level at which data quality decreases (Watkins et al., 2022), so we removed them from further analysis. The amount of DNA extracted was assessed using Invitrogen Quant-iT™ PicoGreen™ (Thermo Fisher Scientific).

We generated DNAm data using the Illumina Infinium MethylationEPIC BeadChip for the remaining 424 participants. We then removed a further 96 participants from the sample set due to poor quality DNA extraction (as determined by high numbers of undetected probes on the EPIC BeadChip; these 96 samples had substantially lower levels of DNA than the 328 remaining for analysis (means of 136.8ng vs 220.7ng; t-test p=1.9e09)). Another 35 samples had a mismatch between the gender they self-reported in the study and sex as predicted by probe signal intensities targeting sites on the X and Y chromosomes. These mismatches were likely due to erroneous sample swaps. We confirmed this by showing that, by showing that, whereas chronological age and age estimated from DNA methylation were methylation highly correlated among the 293 samples whose self-reported gender matched their probe intensity (Horvath clock R=0.63, Hannum clock R=0.69), correlation among the 35 with a mismatch was very low (Horvath clock R=-0.01, Hannum clock R=0.18). This left us with 293 participants (224 Black and 69 white) with DNA methylation data for analysis.

MESA is a longitudinal US-based cohort that was set up to investigate subclinical cardiovascular disease in individuals free of clinical cardiovascular disease at recruitment. MESA comprises a total of 6814 participants aged 45-84, who were recruited to the study from six field centers across the US. We utilise data from a random subset of participants recruited from four of the field centers (Baltimore, MD; Forsyth County, NC; New York City, NY; and St. Paul, MN) who had blood samples collected at the Exam 5 data collection (2010-2012), and DNAm assayed using the Illumina Infinium HumanMethylation450 BeadChip (total n=1264, aged 55-94 at time of blood draw). DNAm was measured in monocytes isolated from whole blood with over 90% purity (Liu et al., 2013). To ensure an appropriate comparison dataset to MBMS, we included the 975 participants who were US-born and self-identified as non-Hispanic Black (n=229) and non-Hispanic white (n=555), additionally including individuals who identified as Hispanic (n=191).

We utilised only US-born MESA participants because research indicates these groups systematically differ in their self-reports of racial discrimination, with responses in the latter group differential by age at immigration and duration in the US (Brondolo et al., 2015; Dominguez, Strong, Krieger, Gillman, & Rich-Edwards, 2009; Krieger, Kosheleva, Waterman, Chen, & Koenen, 2011). Recruitment of the three racialized groups was uneven across the four recruitment sites; please see Supplementary Table 1 for a breakdown of numbers.

|  | Approximate equivalent to Boston | |  | |
| --- | --- | --- | --- | --- |
|  | New York city, NY | Baltimore, MD | St Paul, MN | Forsyth County, NC |
| Black N.H. | 93 | 132 | 0 | 4 |
| Hispanic | 67 | 0 | 124 | 0 |
| White N.H. | 71 | 168 | 270 | 46 |

Supplementary Table 1: breakdown of racialized group recruitment from the 4 MESA sites that generated DNAm data; for the 975 US-born participants only.

#### Social exposures

| **Exposure** | **Description** |
| --- | --- |
| **Childhood exposure to racialized and economic adversity** | |
| Jim Crow birth state | For both MBMS and MESA, this is a true/false variable representing whether or not the participant was born in a Jim Crow state. Jim Crow states are the 21 US states (plus the District of Columbia) which permitted legal racial discrimination prior to the 1964 US Civil Rights Act. |
| Parent’s highest education | For both MBMS and MESA, this was measured using three categories: < high school; > high school and <4 years college; >4 years college (reference level) |
| Participant’s highest education | For both MBMS and MESA, this was measured using three categories: < high school; > high school and <4 years college; >4 years college (reference level) |
| **Adult exposure to racialized and economic adversity** | |
| Household poverty to income ratio | For both MBMS and MESA, this measure is derived from participants' ratio of household income in 2010 dollars to the US 2010 poverty line given their household composition. |
| Index of Concentration at the Extremes for racialized economic segregation | For both MBMS and MESA, this measure captures extremes of privilege and deprivation, measuring both economic and racialized segregation at the census tract level (Krieger et al., 2016). Score ranges from -1 to 1 representing 100% non-Hispanic Black residents among the lowest 2 census wealth brackets, to 100% non-Hispanic white residents among the 2 highest wealth brackets. |
| Black carbon exposure | **MBMS**: measured as the cumulative average exposure to ambient black carbon (in μg/m3) at the longitude–latitude of their residential address for the 1-year prior to the individual’s study enrolment (2008-2010) (Krieger, Waterman, Gryparis, & Coull, 2015).  **MESA**: measured as light absorption coefficient (in 10^–5^/m; 1 unit approx. equivalent to 1µg/m3) for the addresses lived at during the year prior to exam 5 (Keller et al., 2015). |
| NOx exposure | **MBMS**: measured as Pollution Proximity Index (<https://datacommon.mapc.org/calendar/2020/july>), which uses NOx measurements to construct a weighted score of roadway pollution, measured on a scale of 0-5, with measurements taken over the year 2012.  **MESA**: NOx was measured in parts per billion for the year prior to recruitment (Keller et al., 2015). |
| Experiences Of Discrimination (EOD) (**MBMS only**) | A validated self-report questionnaire measuring domains of exposure to racial discrimination. Score range 0-9, categorised into 0, 1-2, 3+ based on pervious work indicating a non-linear relationship (Krieger, Smith, Naishadham, Hartman, & Barbeau, 2005). |
| Major Discrimination Scale (MDS) (**MESA only**) | A validated self-report questionnaire measuring domains of exposure to racial discrimination; combined with the attribution aspect from EDS (everyday discrimination scale) to enable comparability between EOD and MDS. Score range 0-5, categorised into 0, 1-2, 3+ based on pervious work indicating a non-linear relationship (Krieger et al., 2005). |

Supplementary Table 2: Detailed description of study exposure variables

#### DNA methylation

For MBMS, DNA was extracted from frozen blood spots in 2021 and bisulphite converted with the EZ DNA Methylation-Lightning™ Kit (Zymo Research) according to the manufacturer’s instructions. The eluant from the bisulphite-converted DNA was then applied to the Illumina Infinium MethylationEPIC Beadchip to measure DNA methylation, according to the manufacturer’s protocol. The EPIC BeadChips were scanned using Illumina iScan, with an initial quality review conducted with GenomeStudio. Sample QC and normalisation were conducted using the pipeline implemented in the *meffil* R package, which has previously been described in detail (Min, Hemani, Davey Smith, Relton, & Suderman, 2018). DNA methylation is reported in beta values; this measures methylation on a scale of 0 (0% methylation) to 1 (100% methylation).

DNA extraction and processing has previously been described in detail for MESA (Liu et al., 2013), but briefly, in 2012-2013 DNA was extracted from purified monocytes and bisulfite-converted using the EZ-96 DNA MethylationTM Kit (Zymo Research, Orange, CA, USA). The eluant from the bisulphite-converted DNA was then applied to the Illumina Infinium HumanMethylation450 BeadChip, according to the manufacturer’s protocol. The BeadChips were scanned using the Illumina HiScan reader and summarised using GenomeStudio. Sample normalisation was conducted using the R package lumi, with QC checking for ‘sex and race/ethnicity’ mismatches, and outlier identification. We converted the methylation M-values provided by MESA to beta values using the *m2beta* function in the R package *lumi* (Du, Kibbe, & Lin, 2008).

Because frozen blood spots are whole blood samples, they contain a multitude of blood cell types; resultantly MBMS DNAm data reflects a composite measure across cell types that have differing DNAm profiles. This is likely to result in averaging of DNAm across cell types at many loci. In contrast, MESA data is from a purified cell type (monocytes) that typically comprise x% of whole blood; MESA DNAm data will reflect DNAm levels in monocytes only. Thus, although we might expect some comparability between the DNAm measurements, there will be differences between the methylation profiles that arise from differing DNAm profiles of the cell types. As EWAS examines associations at single loci, there is no way to increase comparability between the datasets; but we see this as an opportunity for evidence triangulation, using differing data sources to strengthen inferences (Lawlor, Tilling, & Davey Smith, 2016). It may be that associations are easier to detect in MESA because signal will not be lost via averaging over multiple cell types.

#### EWAS

For EWAS in both MBMS and MESA we adjusted for age, reported gender (MBMS)/sex (MESA), smoking status, blood cell count proportions (estimated in meffil using the “blood gse35069 complete reference” (Houseman et al., 2012)), and batch effects (estimated using surrogate variables [SVs] calculated using the R package *sva* (Leek, Johnson, Parker, Jaffe, & Storey, 2012)). We used SVs to adjust for batch rather than batch variables directly due to small cell frequencies (due to stratifying the dataset). We chose to use 5 SVs for MBMS and 10 for MESA as these accounted for the majority of batch variation and, in MBMS, DNA input level. Their inclusion in EWAS models shifted EWAS lambda values to approximately 1. EWAS p-value thresholds were adjusted for multiple tests as recommended in the literature: as MBMS utilised the EPIC array with 850k features, the threshold was 9e-08 (Mansell et al., 2019); as MESA utilised the 450k array the threshold was 2.4e-07 (Saffari et al., 2018). Categorical exposures with more than two levels (parent’s highest education, participant’s highest education, experiences of discrimination) were run as multiple binary EWAS, with each level compared to a pre-defined reference group (reference groups are identified from previous work and are detailed in Table 1 in the main text).

We did not perform direct replication analyses between MBMS and MESA, because the substantial differences between the two cohorts would make direct comparisons too difficult to interpret. This is particularly because of the differing sample types (whole blood, which represents aggregated methylation profiles of many cell types, vs purified monocytes), as well as significant population differences including age and exposure histories.

#### Sensitivity analysis

We tested whether our results were influenced by population stratification by additionally adjusting each EWAS for the first 10 genetic principal components (PCs), and then using Pearson correlation to test effect size replication for the top 10, 25, 50, 100 and 200 sites in each EWAS. MESA used the Affymetrix Genome-Wide Human SNP Array 6.0 to genotype 8402 participants; this included 933 out of 975 of the US-born participants with DNAm data. Genetic principal components were generated by MESA: 23,428 flagged SNPs and 6,849 SNPs in long range LD were removed before principal components were computed per chromosome, and then combined across chromosomes to give the final PC values. We used the genetic PCs estimated in participants stratified by self-reported racialized group, to match our own study setup.

#### Biological enrichments of top 100 sites

These analyses included the top 100 sites from each EWAS to ensure were sufficient numbers to assess enrichment. Gene set enrichment analyses were conducted using the R package *missMethyl*. Enrichments among sites previously associated with other phenotypes and exposures were tested by comparison to published EWAS summary statistics recorded by the EWAS catalog (Battram et al., 2022) using previously described methods to create phenotype and exposure categories (Elliott et al., 2022). Enrichment of up to 26 categories could be tested (categories were not tested if no CpGs in the category were in the test set of sites); between 8 and 19 categories were tested for our exposures. Enrichments for tissue-specific chromatin states, genomic regions and transcription factor binding sites (TFBS) were assessed the R package *LOLA* (Sheffield & Bock, 2016). We tested for enrichment of chromatin states using the Roadmap Epigenomics chromHMM imputed 25 chromatin states (Ernst & Kellis, 2015; Roadmap Epigenomics et al., 2015); for genomic regions using Illumina annotations (Zhou, Laird, & Shen, 2017); and for TFBS using the Encode (Consortium, 2012; Davis et al., 2018) TFBS set, comprising ChIP-seq data on 161 TFs. All sets of genomic loci are available through <http://lolaweb.databio.org>; we reduced these to features measured using blood. For all LOLA analyses, each DNAm site was extended to a 200bp region centred at the site, removing overlapping sites to prevent inflation.

#### Lookup of associations in *a priori* specified genomic locations

We hypothesised a priori that our EWASs would detect DNAm sites that have been robustly associated with our study exposures, or factors that might relate to our exposures, in previous studies. Because some of the structural measures have not been tested via EWAS before, we also included hypothesised pathways. We grouped previous studies as follows with regard to our exposures: parent’s and participant’s education (previous EWAS of education), household poverty to income ratio (previous EWAS of individual-level socioeconomic status), ICE race plus income (previous EWAS of neighbourhood level socioeconomic measures), black carbon, LAC, and NOx (previous EWAS of air pollution measures), and EOD and MDS (previous EWAS of experiences of racial discrimination). No equivalents were available for Jim Crow birth state. DNAm sites associated with these domains in previous literature were identified through the EWAS catalog and literature searches. Where publications reported genes rather than CpG sites, we took all CpG sites within 1000bp of the gene (to incorporate transcription start sites and promotors). To ensure the sites we tested had robust associations with the exposures, we only took DNAm sites forward where they were identified in at least two separate studies, aside from racial discrimination where literature is currently limited, as this was an important focus of our study. EWAS results for each of the tested exposures were then reduced to the sites that had been identified for that particular exposure in previous literature, and tested for association at p<0.05 divided by the number of sites tested.

### Results

#### EWAS results and biological interpretation

We noted that there was a strong overlap between the genome-wide significant DNAm sites between the Jim Crow birth state and the air pollution EWASs in both the Black and white non-Hispanic MESA participants. The sites at which MESA participants were recruited strongly determined their air pollution exposure (with the Columbia site in New York having much higher pollution levels than other sites); and recruitment site was strongly related to whether or not individuals were born in a Jim Crow state (the more southern recruitment sites recruiting more individuals born in Southern states with Jim Crow laws). Suggesting that the associations in the Jim Crow EWAS in the white NH participants were very likely to have been driven by air pollution, in analyses among these participants stratified by Jim Crow birth status for which we ran the air pollution EWAS, we found that effect sizes correlated R = 0.86-0.99 with the original air pollution EWAS (which we took to be good replication given the lower numbers and reduction in variation in air pollution).

#### Sensitivity analysis

We utilised the first 10 genetic PCs in MESA to test whether population stratification might play a role in our EWAS findings. Additionally adjusting for genetic PCs in our EWAS models, we found that effect sizes for the top 10, 25, 50, 100 and 200 sites from each exposure EWAS correlated with our original EWAS: R>0.99 for white NH MESA participants; R>0.97 for MESA black NH participants; and R>0.98 for MESA Hispanic participants, except for R=0.90 for low education. We therefore concluded that population stratification did not drive the results of any of our EWAS. We could not run this sensitivity analysis for MBMS as this study does not have genetic data.

When we subset participants to those recruited at the Johns Hopkins and Columbia sites, we did not run the analysis for the black NH participants because only 4 participants were recruited outside of those two sites. For the white NH participants, we found that effect sizes for the top 10, 25, 50, 100 and 200 sites from each exposure EWAS correlated (using Pearson correlation) with our original EWAS R>0.93; aside from the low education (R>0.79) and air pollution (R>0.57 and R>0.73 for LAC and NOx, respectively) exposures.

#### Meta-analysis

Supplementary Table 3 shows the complete case numbers for the air pollution meta-analyses.

|  | **MBMS** | | **MESA** | | **MESA (JHU + COL)** | |
| --- | --- | --- | --- | --- | --- | --- |
|  | **N** | **N sites** | **N** | **N sites** | **N** | **N sites** |
| **Black carbon/LAC*** | 293 | 0 | 912 | 17 | 496 | 51 |
| **NOx** | 288 | 0 | 912 | 18 | 496 | 79 |

Supplementary Table 3: Meta-analysis results for air pollution exposure in MBMS and MESA. *Black carbon measured in MBMS; LAC (light absorption coefficient) measured in MESA.

#### EWAS catalog

In MBMS, we see enrichment for inflammation for both NOx and LAC EWAS among Black NH participants, and in the NOx meta-analysis. Among the full sample of US-born MESA participants, the LAC and NOx EWAS were enriched for infection and cancer among both Black NH and white NH participants; the NOx EWAS was enriched for metabolic factors among MESA Hispanic participants; and the LAC and NOx meta-analyses were both enriched for lung, inflammation, diet, cancer and alcohol. When we restricted MESA to participants recruited at the New York and Baltimore sites, we observed consistent enrichment for infection in Black NH and white NH individuals; with enrichment for inflammation in Hispanic individuals; and in the meta-analyses, for LAC we observed enrichment for infection and cancer, and for NOx we observed enrichment for infection.

We also found some consistent enrichments for measures of structural discrimination. Among both MESA Black NH and Hispanic participants, the racialized economic segregation EWAS was enriched for neurological traits. Among both the white NH and Hispanic participants, household poverty to income ratio EWAS was enriched for SEP and education. In the MESA subgroup analysis, enrichment for prenatal exposures was observed for the Jim Crow birth state EWAS among the Black NH and Hispanic participants. detailed results are displayed in supplementary table 5.

##### Enrichment for genomic features

When we looked at enrichment of genomic locations of the top 100 sites (using p<0.05 as a threshold), we found that among MBMS Black NH participants, NOx was the only exposure with notable associations, enriched for two chromatin states (bivalent promoter and promotor upstream of transcription start sites) in addition to being located in promoters, CpG islands and CpG island shores, and enrichment for 9 transcription factor binding sites. We did not observe similar associations for black carbon exposure; and we did not observe any striking enrichments in the MBMS white NH participants. In the MBMS meta-analyses, NOx was enriched for two chromatin states (promotor upstream of transcription start sites and promotor downstream of transcription start sites), three genomic regions (being located in promoters, CpG island shores, and 1-5kb upstream of the TSS), and 20 TFBS.

Among MESA Black NH participants, we observe very similar enrichment for LAC and NOx. NOx is enriched for two active chromatin states (transcription regulation and promotor downstream of TSS 1); three genomic locations (CpG islands, CpG shores and 1-5kb upstream of the TSS); and 53 TFBS. LAC is enriched for three chromatin states (transcription regulation, promotor downstream of TSS 1 and promotor upstream of TSS); three genomic locations (CpG islands, CpG shores, and promotors); and 50 TFBS. They overlap 2 chromatin states, 2 genomic locations, and 45 TFBS. The only structural measure with notable enrichments among the MESA Black NH participants is birth in a Jim Crow state, which is enriched for transcription regulation chromatin state, and 5 TFBS.

Among MESA white NH participants, we see slightly different enrichment patterns for air pollution in the full cohort. Enrichment is instead seen for chromatin states related to transcription regulation, and enhancer for LAC. We only observe enrichment for genomic regions for NOx (intergenic CpG islands); LAC and NOx are enriched for a similar set of TFBS. There are some strong enrichments for Jim Crow birth state, which as discussed above was an artefact of air pollution differences. Among MESA Hispanic participants, LAC exposure shows some associations with active genomic regions, enriched for bivalent promotor chromatin states, location in CpG islands, and 15 TFBS. In the MESA full cohort meta-analysis for LAC and NOx we observe enrichment for chromatin states related to transcription regulation; genomic regions related to intergenic CpG islands; and 24 and 18 TFBS. When we restrict MESA to the New York and Baltimore sites, among white NH participants we observe similar chromatin state enrichments; NOx enrichments for CpG island regions; and a reduction in the number of TFBS enrichments for LAC. In the MESA subset meta-analysis we observe LAC and NOx enrichment for chromatin states related to transcription regulation and promotors; location in CpG islands; and for 48 and 41 TFBS.

Notably, genomic feature enrichments for NOx among both MBMS and MESA Black NH participants involved similar genomic locations (CpG islands and shores) and chromatin states (related to promotors), as well as 6 of a possible 9 TFBS; suggesting that this higher-level analysis may illustrate potential overlap of biological mechanisms or genome regulation between the two cohorts, even though association with specific DNAm sites was not observed between the two studies. This similarity of genomic feature enrichments was not apparent for black carbon. However we think this strengthens implications of our findings, given the heterogeneity between the two cohorts.
